# Genomic and eco-geographic features of locally adapted inversions in wild sunflowers

**DOI:** 10.64898/2026.08.04.742882

**Authors:** Yue Yu, Eric Gonzalez Segovia, Ji Wang, Alexandra Legendre, Veronique Gautier, Stéphane Munos, Marco Todesco, Loren H. Rieseberg

**Affiliations:** Department of Botany and Biodiversity Research Centre, University of British Columbia, Vancouver, British Columbia, Canada; Key Laboratory for Bio-resource and Eco-environment of Ministry of Education, College of Life Sciences, Sichuan University, Chengdu, P. R. China; Laboratoire des Interactions Plantes Microbes-Environnement (LIPME), Université de Toulouse, INRAE, CNRS, LIPME, Castanet-Tolosan, France; Gentyane Platform UMR INRAE/UCA GDEC 5 chemin de Beaulieu 63000 Clermont-Ferrand, France; Institut de Recherche en Santé Digestive (IRSD), INSERM, INRAE, Université de Toulouse, France; Michael Smith Laboratories, University of British Columbia, Vancouver, British Columbia, Canada; Department of Biology, University of British Columbia, Kelowna, British Columbia, Canada

## Abstract

Chromosomal inversions are increasingly recognized as important drivers of local adaptation and ecological divergence because they suppress recombination and maintain adaptive allele combinations despite ongoing gene flow. However, the eco-evolutionary conditions favouring the establishment of such indirectly adaptive inversions, as well as the genomic features that distinguish them from other inversions remain incompletely understood. In this study, we investigated these questions in a wild sunflower system comprising two species: *Helianthus debilis* and *Helianthus praecox*, which exhibit diverse ecotypes and varying degrees of geographic overlap across Texas and Florida in the USA. To resolve the evolutionary relationships between and within these species, and identify potentially adaptive inversions, we generated haplotype-resolved reference assemblies and integrated comparative and population genomic analyses. We identified three major genetic clusters that only partially corresponded to taxonomic classifications. We further detected 156 inversions across the genome, 11 of which showed signatures consistent with a role in local adaptation. Notably, nine of the 11 putatively adaptive inversions were found in sympatric Texas populations. Together with a similar enrichment of inversions in genome assemblies from sympatric versus allopatric populations, our results suggest that inversions are more likely to evolve in heterogeneous environments with ongoing gene flow than in allopatry. Lastly, locally adaptive inversions were generally larger, contained more genes, and showed greater sequence divergence between haplotypes than other types of inversions. Our findings provide empirical support for the role of gene flow in promoting inversion establishment and identify genomic characteristics associated with indirectly adaptive inversions.

## 1. Introduction

Local adaptation is common in widespread plant species, and often leads to the formation of ecotypes, which are populations with distinct suites of morphological and physiological traits that increase fitness in local environments. Chromosomal inversions and other recombination modifiers play an important role in the formation of ecotypes and in the process of speciation in the presence of gene flow by limiting recombination among alleles contributing to local adaptation and assortative mating (Dobzhansky and Sturtevant 1938; Wellenreuther and Bernatchez 2018; Huang and Rieseberg 2020; Todesco et al. 2020; Felsenstein 1981; Trickett and Butlin 1994; Servedio 2009). Inversions also facilitate the accumulation of Bateson-Dobzhanksy-Muller (BDM) incompatibilities in the presence of gene flow (Navarro and Barton 2003; Noor et al. 2001). Such incompatibilities would otherwise require allopatry for establishment.

Geographical context is important for the establishment of adaptive inversions. In the most widely accepted model, Kirkpatrick and Barton (2006) found that inversions containing two or more locally adapted alleles have a higher likelihood of establishment when there is gene exchange between sympatric or parapatric populations adapted to different environments. This is because the inverted chromosome avoids recombination with incoming, maladapted immigrant alleles, preserving high-fitness, locally adapted gene combinations. These adaptive combinations increase in frequency along with the inversion haplotypes housing them. In contrast, recombination suppression is not advantageous in the absence of maladaptive gene flow, so adaptive inversions are less likely to become established in allopatry. Recent simulation studies further support this prediction by showing that inversions are most strongly favoured when gene flow swamps locally adapted alleles, whereas inversions provide little advantage when locally adapted loci are already resistant to swamping (Schaal et al. 2022).

In addition to inversions that are indirectly adaptive through their effects of recombination suppression (Kirkpatrick and Barton 2006), inversions can also be neutral, underdominant (heterozygous disadvantage), overdominant (heterozygote advantage), or directly beneficial due to breakpoint effects (Dagilis 2022; Villoutreix et al. 2021; Berdan et al. 2021; Connallon and Olito 2022). These evolutionary models are not mutually exclusive. For example, an inversion established because it facilitates local adaptation may subsequently accumulate deleterious mutations and later be maintained by overdominance (Berdan et al. 2021). Nonetheless, theory predicts that inversions arising under different selective regimes should differ in several genomic characteristics (Dagilis 2002; Connallon and Olito 2022). One of the most important of these is inversion length. Underdominant, neutral, and directly beneficial inversions are expected to be relatively small, on average, mainly because smaller inversions arise more frequently and are less likely to incur substantial fitness costs associated with recombination. In contrast, locally adapted inversions are predicted to be larger, contain more genes, and occur in high recombination regions of the genome because the benefits of recombination suppression increase with the number of adaptive alleles captured by the inversion and the levels of recombination between them (Kirkpatrick and Barton 2006; Charlesworth and Barton 2017; Connallon and Olito 2022). Such inversions are also expected to exhibit greater sequence divergence between haplotypes because they are retained due to balancing selection (Wellenreuther and Bernatchez 2018) and to occur more frequently in ecotypes or species with sympatric or parapatric distributions, where divergent selection acts in the presence of gene flow (Huang and Rieseberg 2020; Todesco et al. 2020). An inversion’s location relative to the centromere may also influence its fate. Paracentric inversions, which do not include the centromere generally suffer less underdominance than pericentric inversions, which span the centromere (Kirkpatrick 2010). Thus, paracentric inversions are likely to establish more readily and be more common than pericentric inversions, making them more likely to contribute to local adaptation (Kirkpatrick 2010; Wilkinson et al. 2024).

Recent advances in sequencing and computational biology have made it possible to tackle inversion detection using both comparative and population genomic methods. On the comparative genomics side, the combination of long-read and Hi-C sequencing permits the development of high-quality reference assemblies, which can be compared to enable comprehensive inversion detection. Such inversions were previously difficult to identify in some species because of highly repetitive breakpoints and flanking regions (De Coster et al. 2021; Mérot 2020). Conversely, newly developed population genomic methods permit identification of long haplotype blocks (which often represent inversions) from resequencing data (Li and Ralph 2019). Accurate identification of inversions provides a foundation for studying their origins, geographic distributions, and genomic features. Here we leverage these advances to further investigate the natural history of inversions in wild sunflowers, an evolutionary model system.

Previous studies of inversions in wild sunflowers have focused on three species (*Helianthus annuus, H. argophyllus, and H. petiolaris*) and demonstrated that large inversions play a key role in ecological divergence and speciation in this system (Huang et al. 2020; Todesco et al. 2020; Huang et al. 2025). However, these studies lacked reference genomes for the wild species and only examined large inversions but did not investigate how geographic range overlap influences inversion establishment. In this study, we extend our analyses of inversions to two additional wild species, *H. debilis* and *H. praecox*. These closely related species encompass diverse ecotypes and subspecies and exhibit varying degrees of geographic overlap across Texas and Florida, providing an opportunity to examine how geographic context shapes inversion evolution.

Hybridization between subspecies, combined with uncertainty in subspecies identification due to phenotypic similarity in contact zones, has obscured evolutionary relationships in this system. To resolve these complexities and comprehensively identify inversions, we employ a combination of HiFi sequencing, Hi-C scaffolding, and population-level genotyping-by-sequencing (GBS) data. We first assemble high-quality haplotype-resolved reference genomes for both species and use comparative genomics analyses to identify structural variation within and between species, with a particular focus on inversions. We then conduct population-level analyses to characterize polymorphic inversions and assess their role in environmental adaptation. Results from these analyses allow us to clarify patterns of genetic structure, refine taxonomic classification, and identify adaptive inversions across the system. In doing so, we test two central hypotheses: (1) locally adaptive inversions are more likely to arise and persist in the presence of gene flow across heterogeneous environments; and (2) locally adapted inversions are expected to be longer, contain more genes, exhibit greater sequence divergence, and exclude centromeres.

## 2. Results

### 2.1 Haplotype resolved reference genomes

We generated PacBio HiFi and Hi-C data for one individual of *H. debilis* ssp*. tardiflorus* from Northwest Florida and for one individual of *H. praecox* ssp. *hirtus* from western Texas (Figure 2a; Table S1). Using a combination of long reads and long-range interaction data and the assembly pipelines described previously, we produced final phased chromosome-level assemblies for *H. debilis*: HDebH1, HDebH2 and *H. praecox*: HPraH1, HPraH2, each with 17 chromosomes.

HDebH1 had a total length of 3.37 Gbp, a contig N50 of 21.79 Mbp, an assembly N50 of 194.30 Mbp, and a BUSCO score of 98.5%. Annotation indicated that the HDebH1 genome is mostly composed of repeats (84.22%), with 43,903 genes (Table 1). Assembly statistics for HDebH2 are very similar (Table 1), with a slightly higher BUSCO score of 99%, which is reflected in a higher number of annotated genes of 44,695 and a lower proportion of repeats of 83.83% (Table 1). Unassigned fragments for HDebH1 and HDebH2 are 71.95 Mbp (90.23% repeats) and 42.12 Mbp (93.43% repeats) in length, respectively, with BUSCO scores of 0.6% and 0.1%, and 3661 and 2831 BRAKER-annotated genes, respectively.

**Table 1:** Genome assembly and genome annotation statistics for the two haplotypes of *H. debilis* (HDebH1 and HDebH2) and *H. praecox* (HPraH1 and HPraH2).

| Assembly Features | HDebH1 | HDebH2 | HPraH1 | HPraH2 |
| --- | --- | --- | --- | --- |
| HiFi coverage | 20X | 20X | 27X | 27X |
| Hi-C coverage | 18X | 18X | 24X | 24X |
| Assembly size (Gb) | 3.37 | 3.35 | 3.04 | 3.17 |
| Chromosome Number | 17 | 17 | 17 | 17 |
| Contig N50 size (Mb) | 21.79 | 23.39 | 31.33 | 26.12 |
| Assembly N50 size (Mb) | 194.30 | 194.77 | 178.22 | 187.12 |
| BUSCO (%) | 98.5 | 99 | 99.1 | 99 |
| LAI | 22.48 | 22.41 | 22.97 | 23.12 |
| QV value | 65.79 | 65.57 | 68.8 | 69.48 |
| Total repeats (%) | 84.22 | 83.83 | 81.52 | 81.28 |
| Transposable elements (%) | 70.77 | 68.39 | 66.65 | 59.47 |
| LTR retrotransposons (%) | 65.79 | 63.04 | 60.44 | 53.29 |
| — Gypsy(%) | 39.79 | 37.44 | 35.73 | 41.29 |
| — Copia(%) | 14.66 | 15.11 | 14.34 | 11.62 |
| DNA transposons (%) | 2.47 | 2.02 | 2.19 | 2.93 |
| Unclassified repeats (%) | 13.45 | 15.44 | 14.87 | 21.81 |
| Tandem repeats (%) | 0.58 | 0.61 | 0.74 | 1.02 |
| Number of genes in chromosome | 43,903 | 44,695 | 43,995 | 45,588 |
| Number of genes Liftoff HA412v2 | 39,741 | 39,714 | 39,797 | 39,796 |
| BUSCO (%) | 97.7 | 98.1 | 98.3 | 98.2 |
| Single-copy BUSCOs (%) | 70.9 | 71.2 | 72.5 | 72 |
| Duplicated BUSCOs (%) | 26.8 | 26.9 | 25.8 | 26.2 |
| Fragmented BUSCOs (%) | 0.8 | 0.9 | 0.7 | 0.8 |
| Missing BUSCOs (%) | 1.5 | 1.1 | 1 | 1 |

For the *H. praecox* assembly, HPraH1 had a total length of 3.04 Gbp, a contig N50 of 31.33 Mbp, an assembly N50 of 178.22 Mbp, and a BUSCO score of 99.1% (Table 1). Annotation revealed a slightly lower proportion of repeats (81.52%) than in *H. debilis*, likely reflecting its smaller genome, but about the same number of genes (43,995). HPraH2 has a longer length of 3.17Gbp, a lower contig N50 of 26.12Mbp, but a higher assembly N50 of 187.12 Mbp and a similar BUSCO score of 99%. The total proportion of repeats was similar, but the number of annotated genes was higher, 45,588 (Table 1). Unassigned fragments for HPraH1 and HPraH2 were much smaller than in *H. debilis*: 0.49Mbp (16.48% of repeats) and 1.49 Mbp (38.47% of repeats), respectively, with very low BUSCO scores of 0 % and 0.1%, respectively. Given their short length, it is unsurprising that no genes were annotated with our pipeline.

The distribution of the annotated repeats, genes and predicted centromeres along each chromosome is displayed in Figure 1. Some repeat hotspots (green line; Figure 1) align with predicted centromeres, and genes are observed to cluster around the two ends of each chromosome (blue line, Figure 1). We also found a 59bp repeat: “TGATCAATTAACCCCAATATGGTCTATTATGACATATATGCTCTTCTATTGCCCGTT GTG” that is present in most of the centromeres of all assemblies.

**Figure 1:**
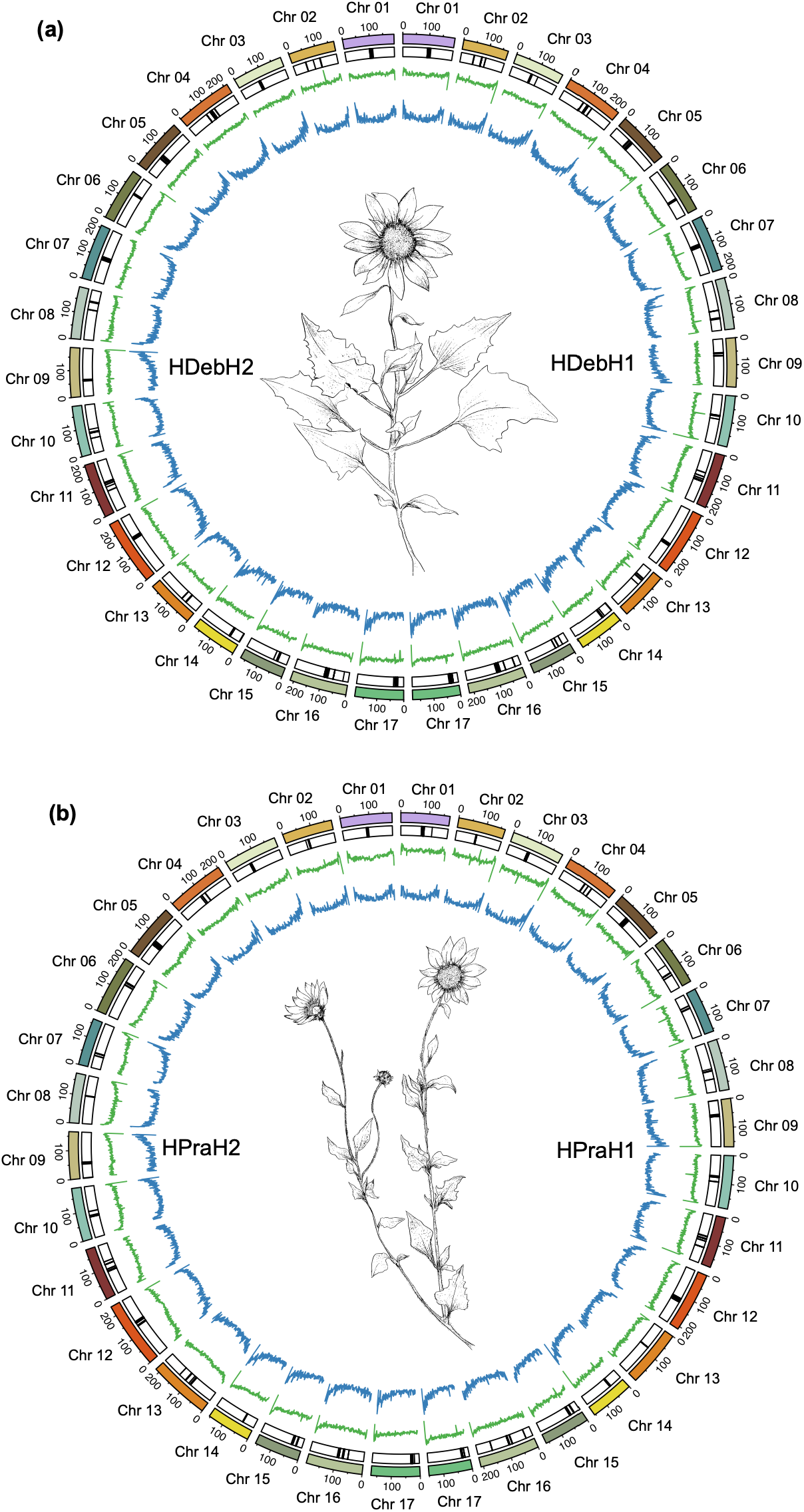
Circos plot for *H. debilis* (a) and *H. praecox* (b). From the outer to inner circle are chromosome number, chromosome position key, predicted centromeres, repeat density (green line) and gene density (blue line) in 1 Mbp windows. Illustration of the assembled species by Yi-Xuan Ye.

### 2.2 SNP calling and identification of major genetic clusters

SNP calling across the 416 high-quality samples yielded 50,511 filtered SNPs. Admixture analysis revealed 12 distinct genetic clusters across all samples, as well as many admixed samples (Figure S1). Genetic relationships in Florida align closely with the subspecific classification, but not in Texas, where subspecies of both *H. debilis* and *H. praecox* can be found in the same clusters. PCA suggests a simpler relationship, with the first two principal components (PCs) explaining 20.51% (PC1) and 9.36% (PC2) of variation and separating the 416 samples into three distinct major genetic clusters (Figure 2c). These three clusters were also confirmed for *K* = 3 in the admixture analysis (Figure 2d). The first cluster (A) consists of all allopatric/parapatric subspecies of *H. debilis* (all populations of DT, DV and DD) in Florida. The second (B) cluster consists of most of the *H. praecox* populations in Texas (PH1, PP1-3, PR4-8). Lastly, all remaining populations of both *H. debilis* and *H. praecox* (PH2, PR1-3, DC2,3,5,6 and DS1-5) are in the third cluster (C).

**Figure 2.**
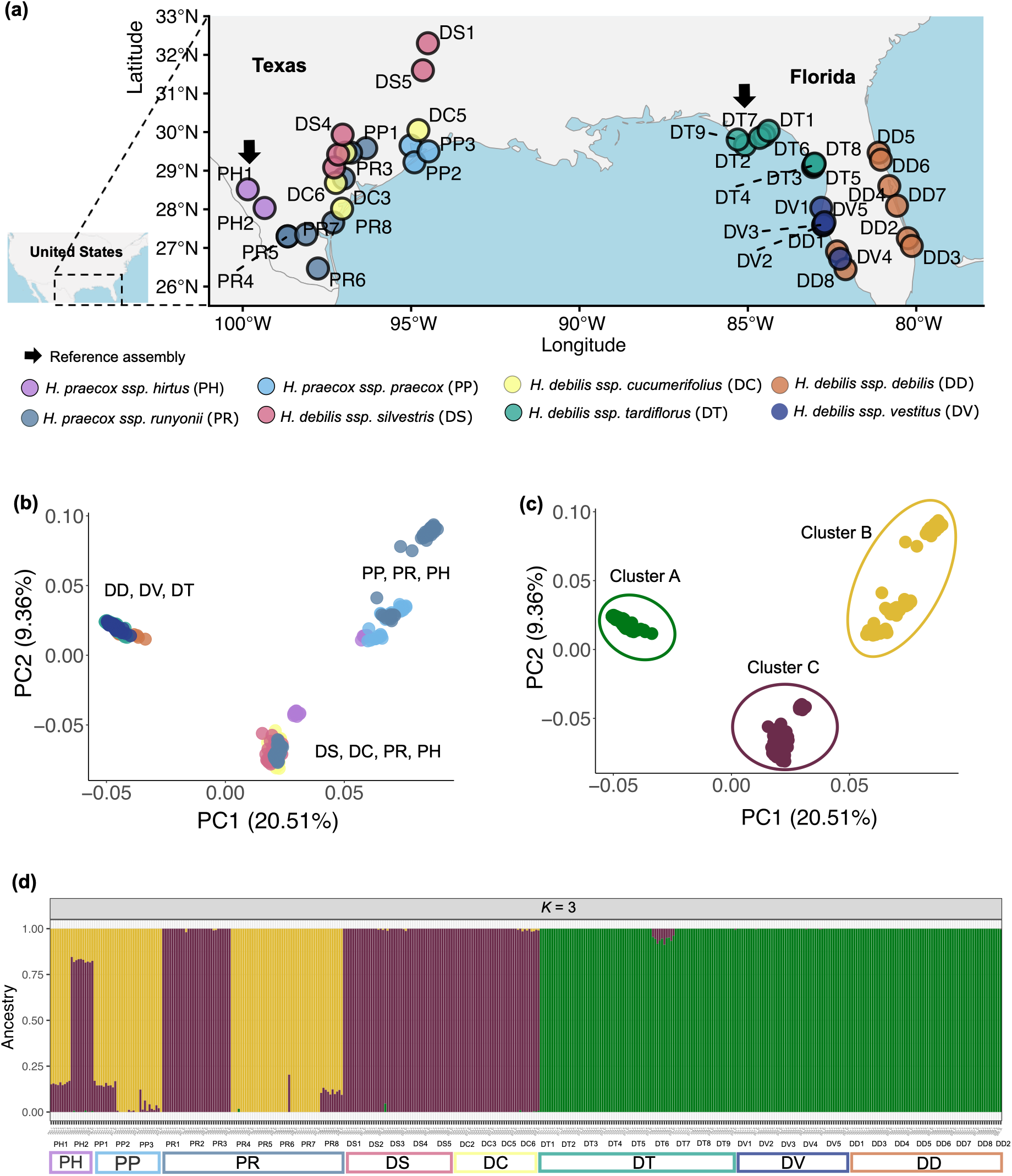
Spatial distribution map and identified genetic clusters of *Helianthus praecox* and *H. debilis* subspecies in Texas and Florida, Southern USA. (a). Locations of the 44 populations in Texas and Florida in this study are represented by each point, colored per subspecies. Arrows in the map point out the collection sites for reference assembly individuals (Table S1). Three populations of the initial 47 populations were excluded because they are outside of the geographical range of interest (DC1, 4 and 7). Location information was obtained from USDA (https://npgsweb.ars-grin.gov/gringlobal/search, accessed December 2021). (b). Principal Component Analyses (PCA) of the first two PCs for all 44 populations, with point colors corresponding to subspecies. (c). PCA of the first two PCs clustered into three major genetic clusters with point color corresponding to admixture result when *K* = 3. (d). Admixture plot of *K* = 3; each column represents an individual, with the corresponding population and subspecies shown below the plot.

### 2.3 Identifying locally adapted inversions

We identified a total of 11 haploblocks ranging from 8.1 to 44.0 Mbp in size using *lostruct* (Table 2). For nine of these, we were able to confirm that they were inversions from the reference genome comparisons. For the other two haploblocks, we relied solely on population genomic analyses to infer that they likely represent inversions. For example, a 29.4 Mbp haploblock, pra17.02, was found on chromosome 17 (Figure 3a). While the three haploblock genotypes (0, 1 and 2) were not clearly separated in a PCA plot (Figure 3b), the intermediate genotype had significantly higher heterozygosity (Figure 3c), as expected if the haploblock were caused by a structural variant. Candidate inversion frequency increases with higher latitude and lower longitude, showing a pattern from southwest to northeast (Figure 3d).

**Figure 3.**
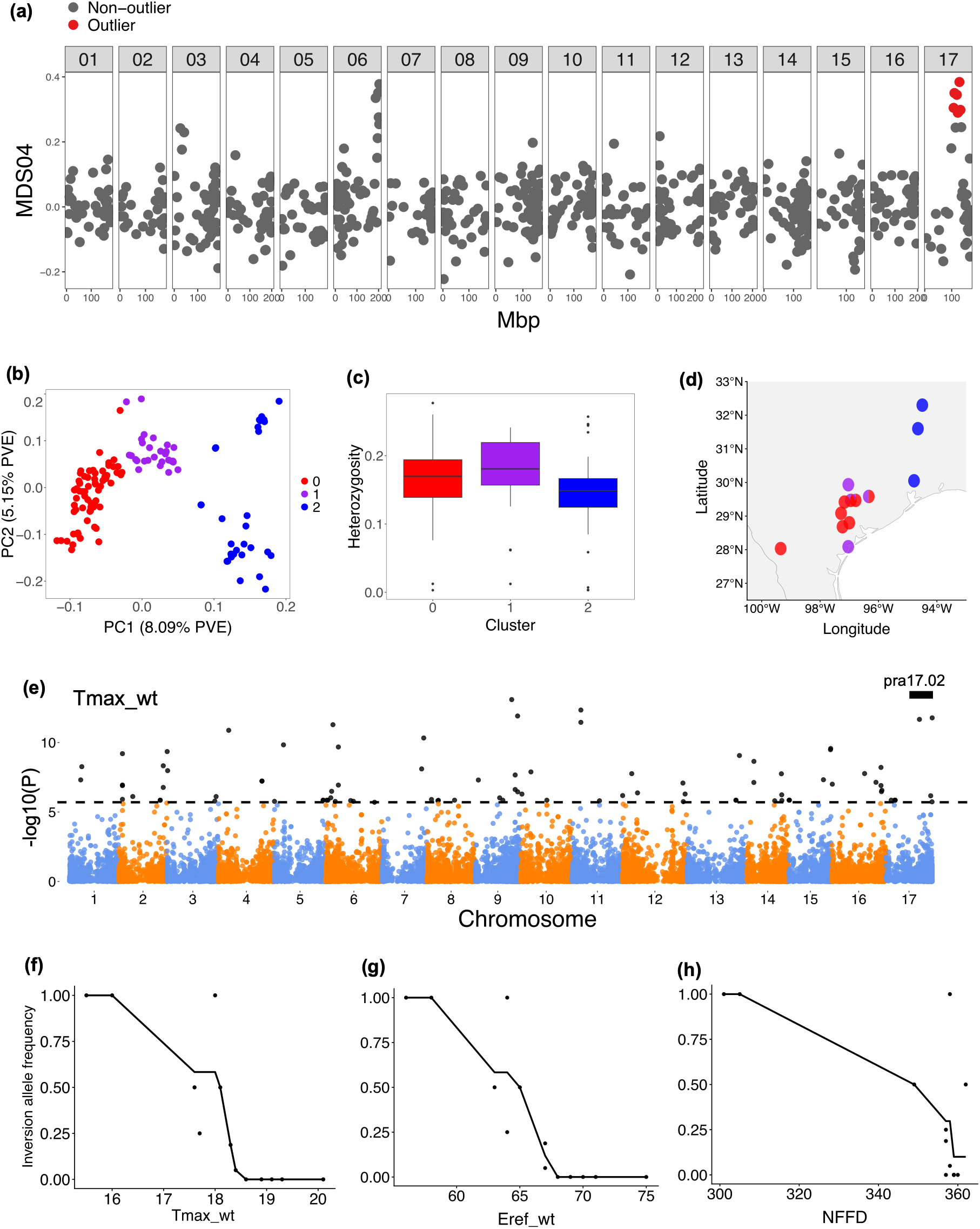
Putative polymorphic inversion pra17.02 among 12 populations in Texas. (a). MDS04 score for all 50-SNP windows (each point) along *H. praecox* chromosome 17, with outlier windows (pra17.02) highlighted in red. (b) Principal component analysis (PCA) of outlier regions, separated into three major clusters based on PC1 values, used to infer inversion genotypes, either homozygous (red and blue) or heterozygous (purple). (c). Mean heterozygosity of each defined cluster from PCA in a boxplot. (d) Inversion genotype frequencies in each population in Texas. (e). Manhattan plot of genome-environment association along all chromosomes for Tmax_wt: Mean maximum temperature for winter. Each point is a locus, the dotted horizontal line is the Bonferroni threshold of significance. Any locus with a -log10(P) above the threshold is marked in black, indicating it is significantly associated with the focal environmental variable. Black bar represents inversion position. Inversion allele frequency change along the gradient of Tmax_wt (f), Eref_wt (g) and NFFD (h), with each point representing a population.

**Table 2:**
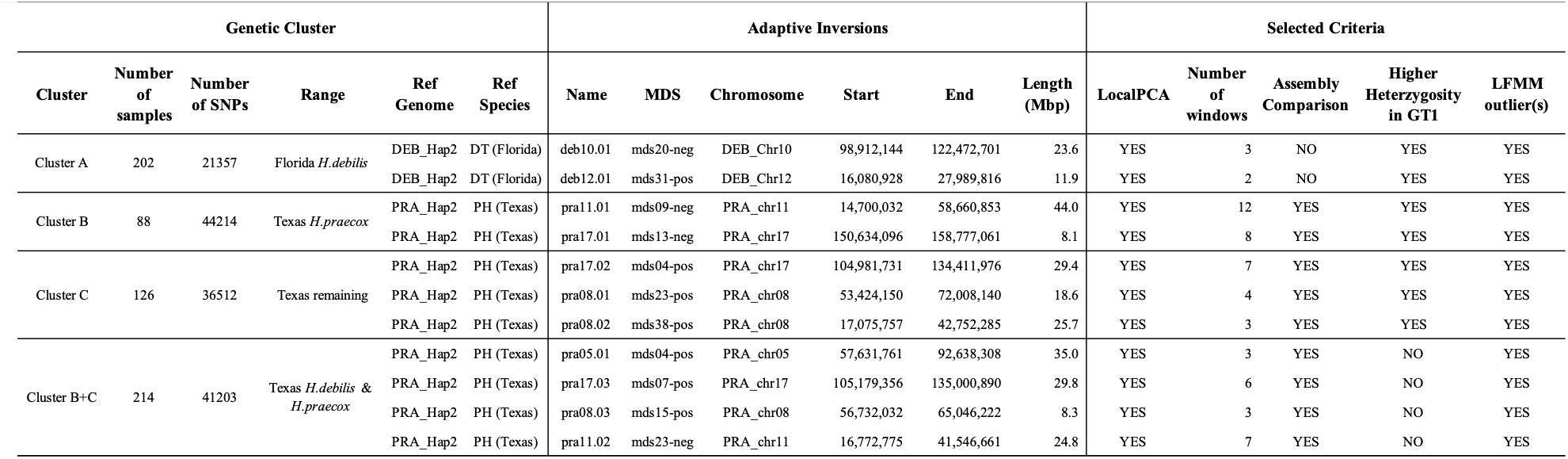
Locally adaptive inversions identified from genetic cluster A, B, C, and B plus C. Three main categories of information are summarized, including: basic summaries of genetic cluster and putative inversions within each cluster, followed by a list of criteria used to determine if these inversions likely contributed to local adaptation.

We provided further evidence of environmental adaptation using LFMM (Figure 3e) to account for background structure and visualized a monotonic cline of pra17.02 for variables affecting growing season temperature, moisture and length: Tmax_wt (spearman’s rho = −0.920; Figure 3f), Eref_wt (spearman’s rho = −0.931; Figure 3g), NFFD (spearman’s rho = −0.565; Figure 3h). All LFMM-outlier loci that overlapped with adaptive coordinates were summarized and reported in Table S2.

Detection signal visualization for all 11 putative locally adapted inversions can be found in Figure S2-S12. Inversions found in cluster A are mostly associated with variation in precipitation and available water content in soil, with clear inversion frequency changes from the west to east coast of Florida. Inversions in sympatric populations in Texas (cluster B and C) overall show a strong association signal with growing season temperature and late-season heat and drought stress. Most inversion frequency changes from coastal to interior Texas, similar to inversion patterns found in *H. argophyllus,* which is also in southeastern Texas (Todesco et al. 2020).

### 2.4 Genome assembly comparisons reveal structural variation and many inversions

For the genome assembly comparison, we first identified all syntenic regions and structural variants (Figure 4). Between HDebH1 and HDebH2, large blocks of syntenic regions were observed, comprising approximately 2.78 Gbp in both haplotypes, which is over 82% of the assembled genome size. In terms of structural variants, we identified 93 inversions, of which 17 were >500 Kbp in length, 1,174 translocations (size range 223 - 548,198 bp), and 3,638 indels between haplotypes (Table 3).

**Figure 4.**
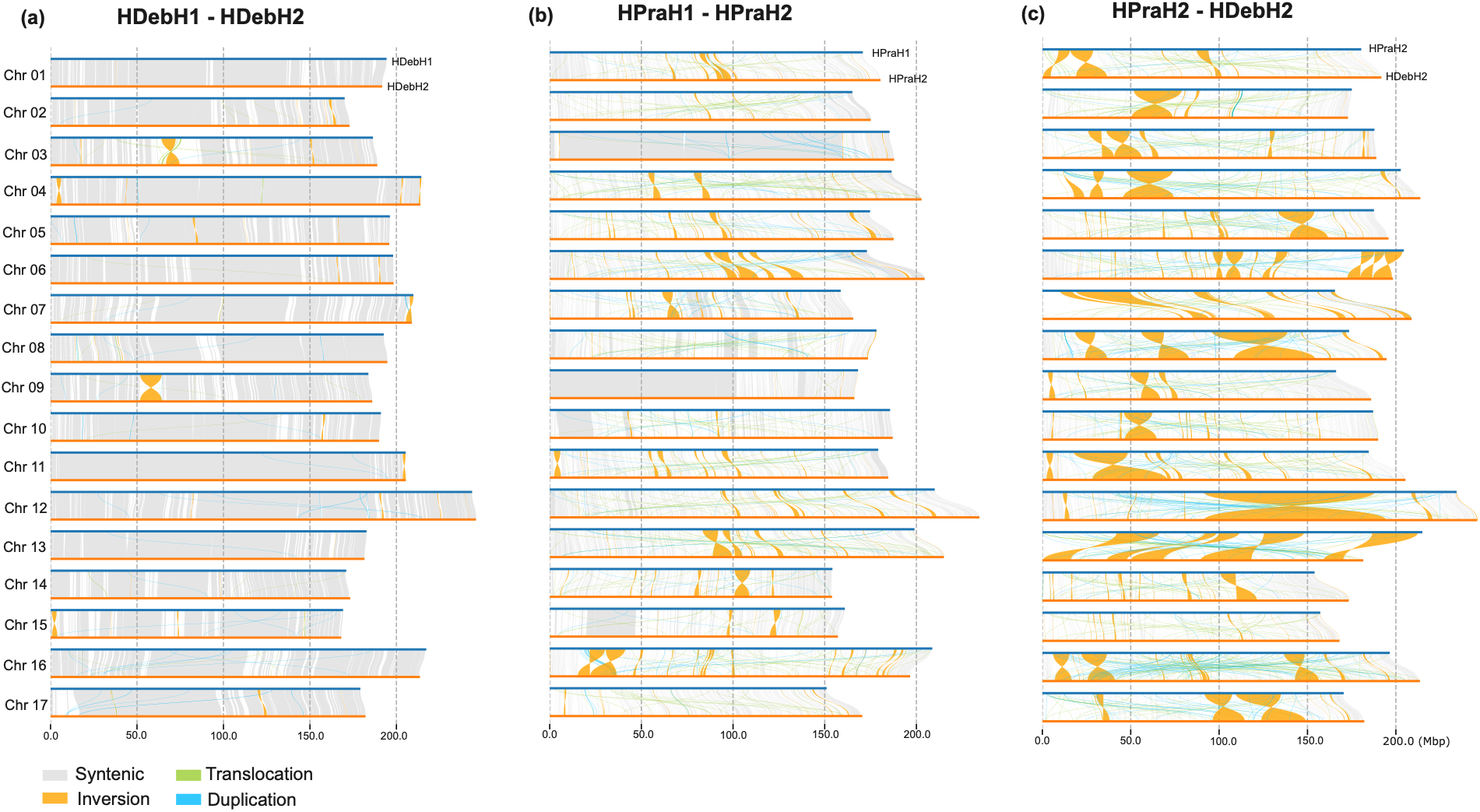
Syntenic regions, inversions, translocations and duplications identified using minimap2 and SyRI, visualized by plotSR between (a) HDebH1 - HDebH2, (b) HPraH1 - HPraH2 and (c) HDebH2 - HPraH2 (between species) for fine scale nucleotide base pair comparison.

**Table 3:**
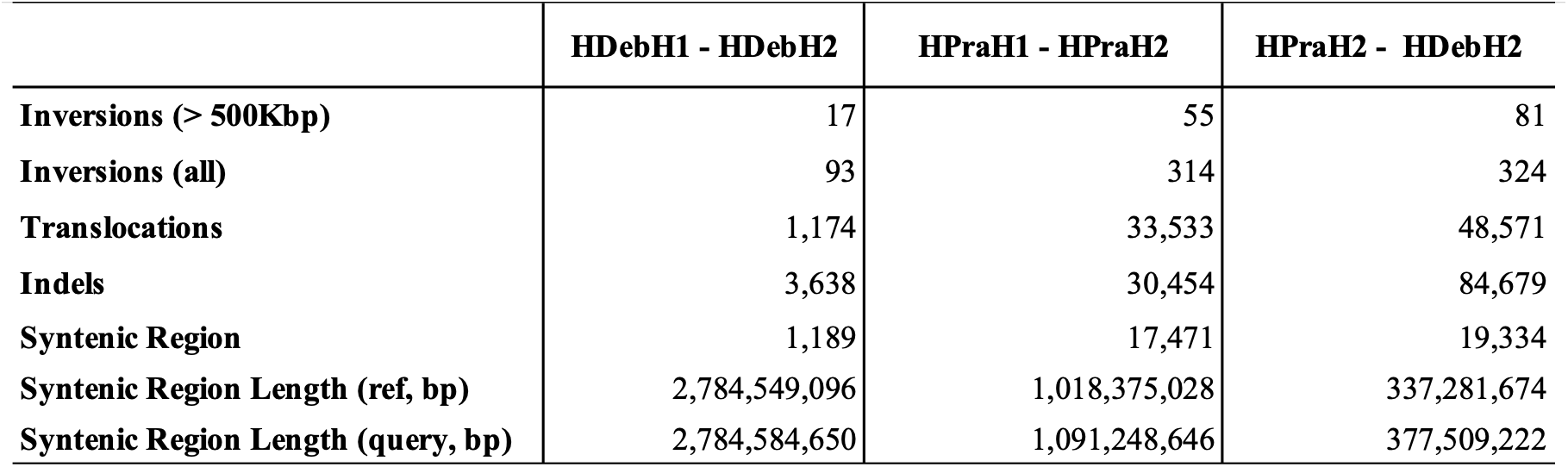
Summary statistics of number and length of syntenic regions, number of structural variations between HDebH1 - HDebH2, HPraH1 - HPraH2 and HDebH2 - HPraH2. In each comparison, the former is the reference assembly, and the latter is the query.

**Table 4:**
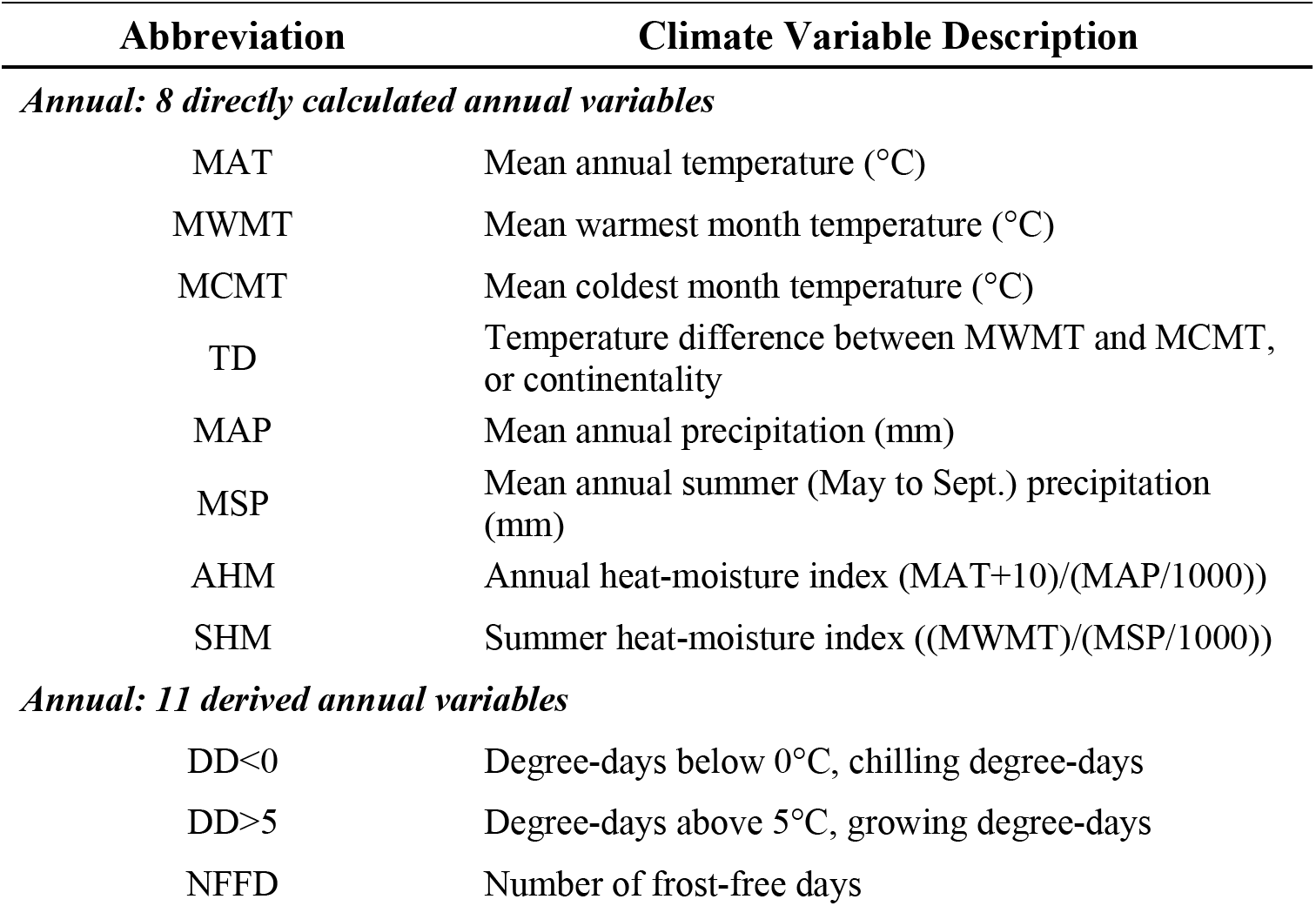
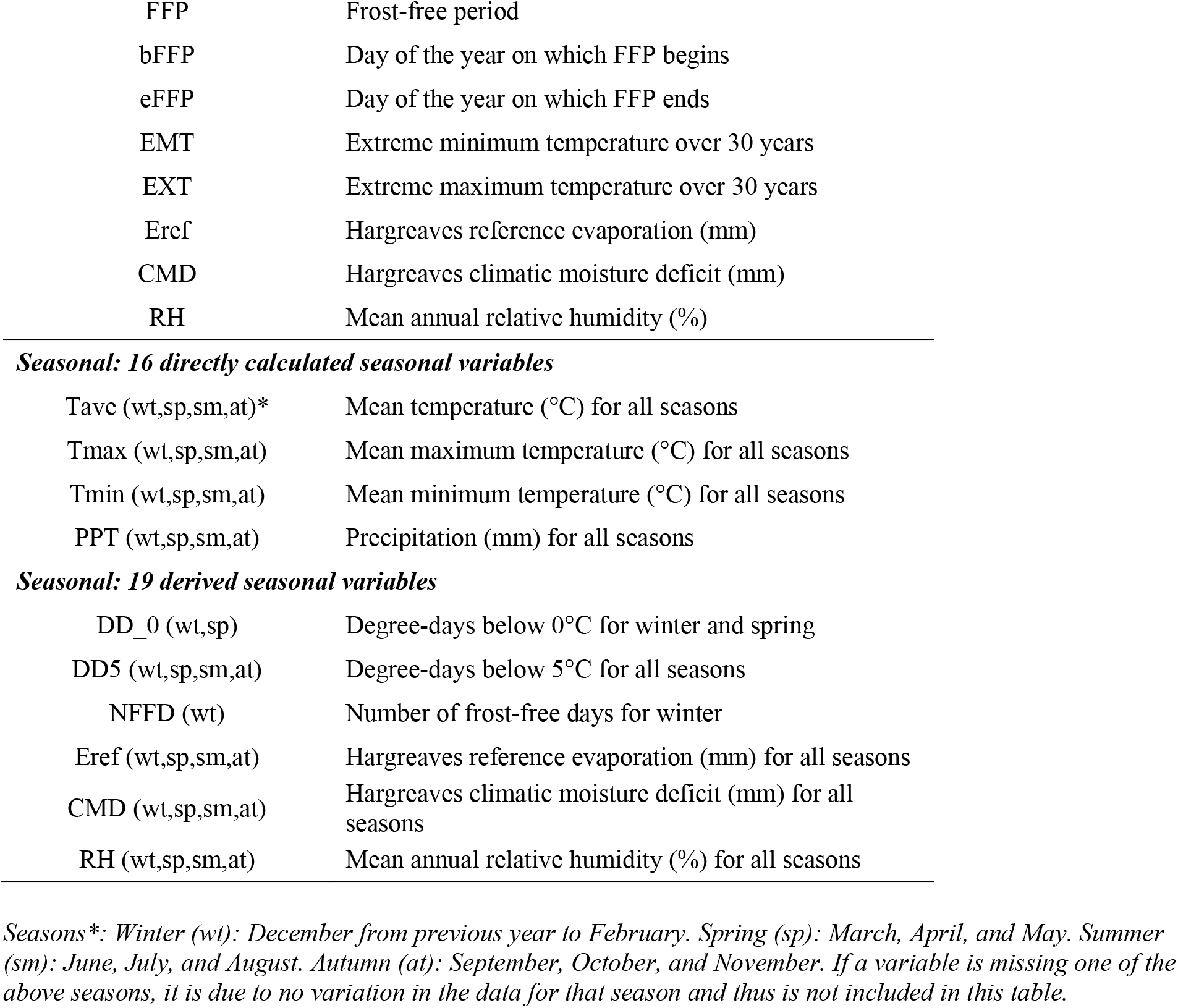
List of the 19 annual climate variables and 35 seasonal climate variables from *ClimateNA* and a brief description of each (Wang et al. 2016).

| Abbreviation | Climate Variable Description |
| --- | --- |
| *Annual: 8 directly calculated annual variables* |  |
| MAT | Mean annual temperature (°C) |
| MWMT | Mean warmest month temperature (°C) |
| MCMT | Mean coldest month temperature (°C) |
| TD | Temperature difference between MWMT and MCMT, or continentality |
| MAP | Mean annual precipitation (mm) |
| MSP | Mean annual summer (May to Sept.) precipitation (mm) |
| AHM | Annual heat-moisture index (MAT+10)/(MAP/1000)) |
| SHM | Summer heat-moisture index ((MWMT)/(MSP/1000)) |
| *Annual: 11 derived annual variables* |  |
| DD<0 | Degree-days below 0°C, chilling degree-days |
| DD>5 | Degree-days above 5°C, growing degree-days |
| NFFD | Number of frost-free days |
| FFP | Frost-free period |
| bFFP | Day of the year on which FFP begins |
| eFFP | Day of the year on which FFP ends |
| EMT | Extreme minimum temperature over 30 years |
| EXT | Extreme maximum temperature over 30 years |
| Eref | Hargreaves reference evaporation (mm) |
| CMD | Hargreaves climatic moisture deficit (mm) |
| RH | Mean annual relative humidity (%) |
| **Seasonal: 16 directly calculated seasonal variables** |  |
| Tave (wt,sp,sm,at)* | Mean temperature (°C) for all seasons |
| Tmax (wt,sp,sm,at) | Mean maximum temperature (°C) for all seasons |
| Tmin (wt,sp,sm,at) | Mean minimum temperature (°C) for all seasons |
| PPT (wt,sp,sm,at) | Precipitation (mm) for all seasons |
| **Seasonal: 19 derived seasonal variables** |  |
| DD_0 (wt,sp) | Degree-days below 0°C for winter and spring |
| DD5 (wt,sp,sm,at) | Degree-days below 5°C for all seasons |
| NFFD (wt) | Number of frost-free days for winter |
| Eref (wt,sp,sm,at) | Hargreaves reference evaporation (mm) for all seasons |
| CMD (wt,sp,sm,at) | Hargreaves climatic moisture deficit (mm) for all seasons |
| RH (wt,sp,sm,at) | Mean annual relative humidity (%) for all seasons |

Seasons\*: Winter (wt): December from previous year to February. Spring (sp): March, April, and May. Summer (sm): June, July, and August. Autumn (at): September, October, and November. If a variable is missing one of the above seasons, it is due to no variation in the data for that season and thus is not included in this table.

**Table 5:**
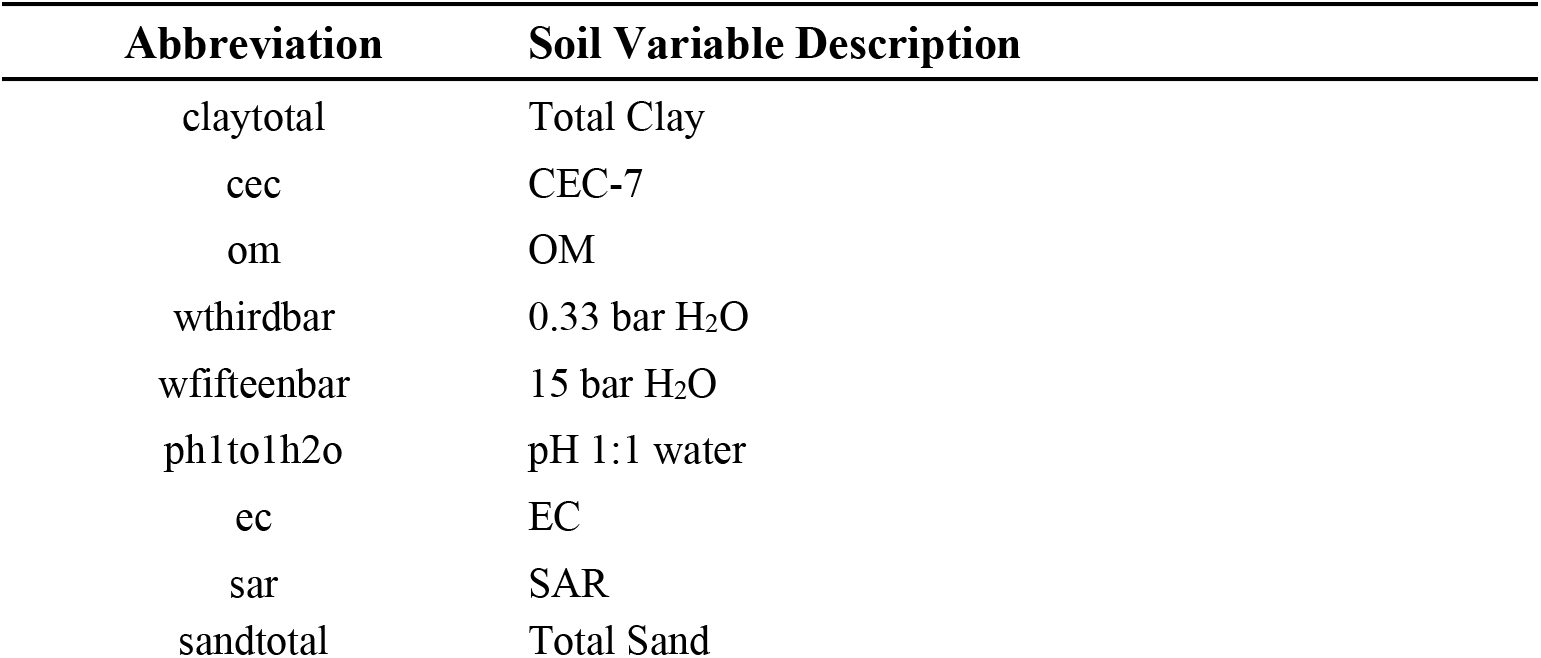
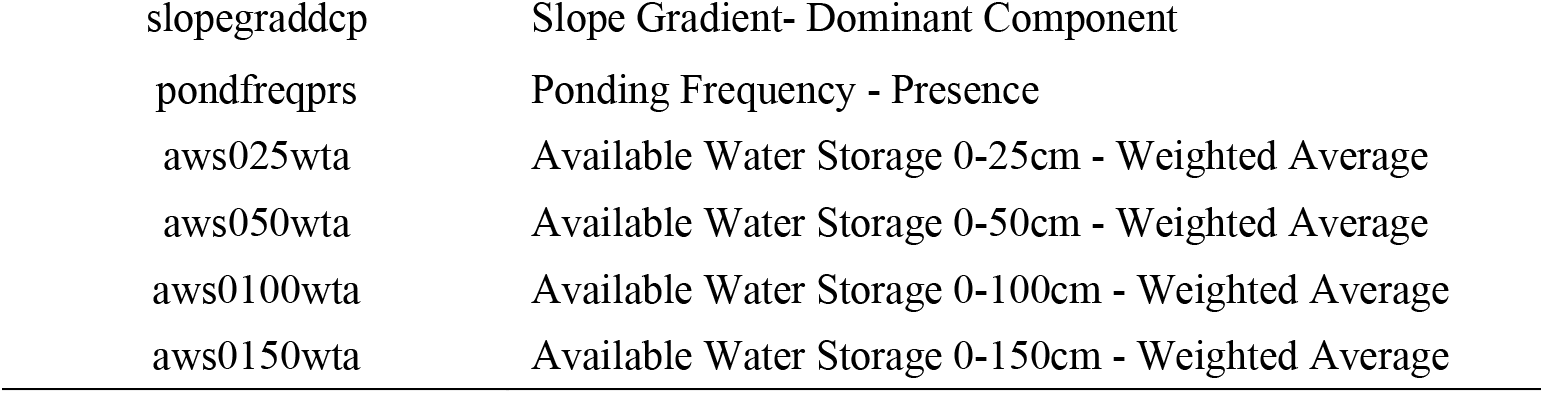
Fifteen soil variables from SSURGO. The first nine soil variables were calculated as an average based on all soil layers between 0 and 180 cm below ground. The remaining soil variables were a single value extracted from SSURGO.

Many fewer syntenic regions were discovered between HPraH1 and HPraH2, covering only around 33% (slightly above 1 Gbp) of the assembled genome, indicating the haplotypes found in the individual sampled for this reference genotype are more differentiated structurally (Figure 4). In all, 315 inversions (of which 55 inversions were >500 Kbp), 33,533 translocations (size range 199 - 43,727 bp), and 30,454 indels were found between haplotypes (Table 3).

The between-species genome assembly comparison at the nucleotide level for HPraH2 and HDebH2 revealed a small number of syntenic regions (0.33 - 0.37 Gbp), representing approximately 10% of the genome (Figure 4). Eighty-one inversions (out of 324) were above 500 Kbp, with many translocations (size range 199 - 187,907 bp) and indels (Figure 4; Table 3). Between-species comparisons with GENESPACE using homologous genes (Figure S13) revealed the same 81 inversions above 500 Kbp. GENESPACE result also revealed and visualized several large inter-chromosomal translocations, some of which are also inverted (Figure S13). These are on average longer in length and captured substantial number of genes and are likely to be strongly underdominant. We used inversion breakpoint coordinates from the nucleotide level comparison for downstream analyses.

The final number of ‘other’ inversions after filtering for inversions >500 Kbp and removal of locally adapted inversions identified with *lostruct* results in 17, 53 and 75 inversions (145 in total) from comparisons of HDebH1 - HDebH2, HPraH1 - HPraH2 and HPraH2 - HDebH2, respectively (Table S3).

### 2.5 Comparison of locally adapted and ‘other’ inversions

Wilcoxon rank-sum tests showed that locally adapted inversions were significantly longer (*P* = 9.1 x 10^-8^; Figure 5a), contained more genes (*P* = 6.4 x 10^-7^; Figure 5b), and were more divergent between haplotypes (*P* = 2.7 x 10^-6^; Figure 5d) than ‘other’ inversions. Note there is a strong correlation (r = 0.841, p-value < 2.2e-16) between inversion length and number of genes.

**Figure 5.**
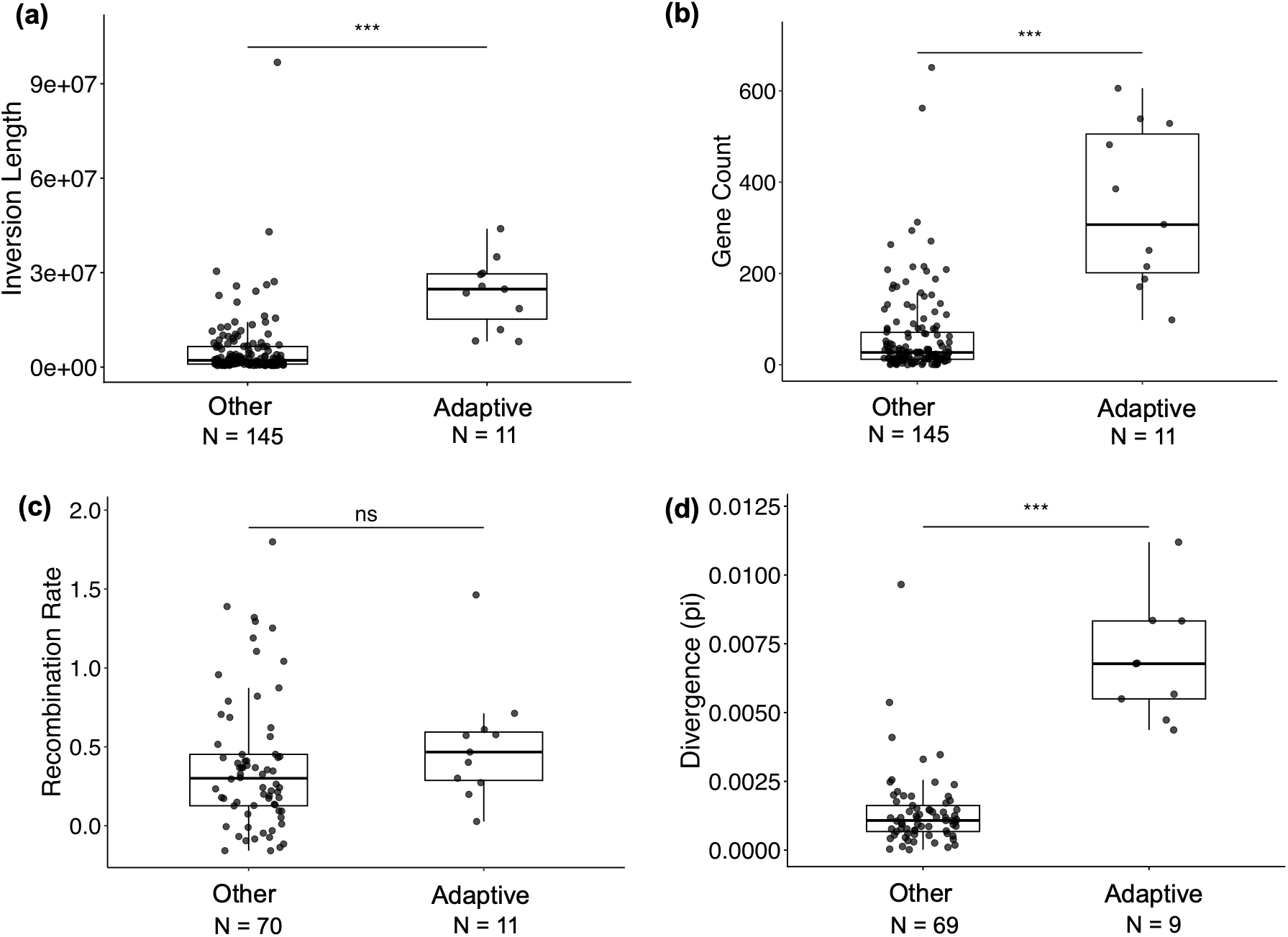
Comparison of inversion length (a), gene count (b), recombination rate (c), and divergence (d) for locally adapted versus other inversions using boxplots. Only inversions > 500 Kbp were included in the comparisons to match the size limit for inversion detection by *lostruct*. Each point represents one inversion, significance level of the Wilcoxon rank-sum test is represented by: *** P<0.001, ** 0.001<P<0.01, * 0.01<P<0.05, ns P>0.05. Number of samples for the non-adaptive group varies in each comparison because some included all 145 “other” inversions, whereas recombination rate and divergence comparisons only included the 70 inversions that were detected between haplotypes within each species. Of these, one inversion was excluded from the divergence comparison as its alignment length was too short, less than 100 Kbp.

Inferred recombination rates were not different between the two groups (*P* = 0.11; Figure 5c). Logistic regression showed that inversion length was a significant predictor of pericentric status, with longer inversions being more likely to overlap the centromere (*P* = 5.16 x 10 ^-5^). In contrast, inversion type was not a significant predictor, after controlling for length (*P* = 0.082).

Most of the inversions in the ‘other’ category display low divergence, with a few highly divergent exceptions (Figure 5d). The latter may well be locally adapted inversions that were not detected by *lostruct*, perhaps because they were small or occurred at low frequency or were overdominant.

## 3. Discussion

Through a combination of haplotype-resolved genome assemblies and population genomics analyses in the wild sunflower system of *H. debilis* and *H. praecox*, we gained a better understanding of the taxonomic classification among their eight subspecies and identified three major genetic clusters. We also detected 11 putative locally adapted inversions and a total of 145 ‘other’ inversions above 500 Kbp. These detected inversions further supported our first hypothesis that locally adapted inversions are more likely to evolve when gene flow is present among heterogeneous environments, such as in sympatric or parapatric populations in Texas relative to Florida. Lastly, we also tested our second hypothesis by comparing predicted characteristics between locally adapted and ‘other’ inversions and confirmed that, as predicted by theory, locally adapted inversions are likely to be longer, capture more genes (highly correlated with inversion length), and exhibit greater divergence between haplotypes. On the other hand, predictions on recombination rates and the likelihood that inversions are pericentric did not show significant differences between the two groups. However, this could be due to poor accuracy of the indirect measure (gene density) we used for estimating recombination rate and the small number of pericentric inversions within the study system.

### 3.1 Three main genetic clusters identified among all *H. praecox* and *H. debilis* subspecies

*H. praecox* and *H. debilis* subspecies were previously characterized based on a combination of crossing relationships (hybrid offspring fertility), phenotypic variation, and the environment to which they are adapted (Heiser et al. 1969). Molecular phylogenetic studies have consistently identified the *debilis* subspecies in Florida as a distinct cluster, but relationships among the Texas subspecies of *H. debilis* and *H. praecox* are less clear and varied among analyses (Rieseberg 1991; Stephens et al. 2015).

Our genetic cluster analyses add to previous taxonomic understanding of these subspecies, confirming that the *H. debilis* subspecies in Florida are distinctively different from the remaining subspecies in Texas (Figure S1, K = 2; Figure 2). The situation in Texas is more complex (Figure 2). While most populations of *H. praecox* form a single cluster (Figure 2, cluster B), one population of *H. praecox* ssp. *hirtus* (PH2) and several populations of *H. praecox* ssp. *runyonii* (PR1-3) clustered with the *H. debilis* subspecies from Texas: *H. debilis ssp. cucumeriflolius* (DC) and *H. debilis* ssp. *silvestris* (DS). This is likely due to misidentification of PH2 and PR1-3 (Figure 2b,c), possibly as a consequence of admixture. DC and PR are sympatric and hybridize in the coastal prairies of southern Texas, where both subspecies intergrade phenotypically. However, it might also result from challenges in species delimitation based on phenotypic and crossing data.

Taxonomic identification of subspecies, even those belonging to different species, is challenging in this system. For example, PR and PH are identified mainly by differences in the density of hairs on their stems, which can lead to misidentification because of continuous variation in this trait. Indeed, our genetic evidence revealed the likely misidentification of the PH2 population near Carrizo Springs, Texas; it is not placed in the same genetic cluster as its neighbour population PH1 but instead with *H. debilis ssp. cucumeriflolius* (DC) and *H. debilis* ssp. *silvestris* (DS) in cluster C (Figure 2 and S1).

Previous literature also mentioned that allopatric *H. praecox* ssp. *praecox* (PP) from Galveston Island, Texas, is morphologically similar to *H. debilis* ssp. *vestitus* (DV), which occurs along the west coast of Florida. Both are procumbent, with the latter less densely pubescent (Heiser et al. 1966). However, they are not sister taxa genetically. Thus, the procumbent habit likely evolved independently due to similar coastal environmental pressure, as growing low and flat along the ground can improve wind resistance, increase stability when on sandy soil, and reduce transpiration (Hesp 1991).

### 3.2 Identification of locally adaptive and ‘other’ inversions

An initial analysis of all 416 samples with *lostruct* failed to detect polymorphic haploblocks; we suspected that this was due to population structure. Therefore, we analyzed the three major genetic clusters independently. As cluster B and C are sympatric in Texas, we also pooled both clusters and found an additional haploblock, pra05.01 (35 Mbp), as well as three haploblocks that overlapped with ones from cluster B and C when examined separately. However, due to population structure introduced by combining cluster B and C, the heterozygosity signal of intermediate genotypes from the haploblock region in the PCA is masked. We kept all inversions from the combined cluster, as these were validated by the *lostruct* analyses of individual clusters, as well as from assembly comparisons. But this observation, as well as our failure to find haploblocks when all samples were combined, raises the concern that complex population structure can result in false negatives. While not all population genomic signatures of inversions were detected for all inversions, validation from assembly comparisons indicates that the putative locally adapted inversions detected in Texas by *lostruct* are real.

Two locally adaptive inversions found in cluster A did not show up in the assembly comparison. This is not surprising given the low heterozygosity of the reference individual. Despite lack of sequence validation, both putative inversions (deb10.01 and deb12.01) were retained for further analyses, given that almost all haploblocks detected by *lostruct* in the sunflower system have ultimately proven to result from inversions (Todesco et al. 2020; Huang et al. 2020, 2023, 2025), including those reported in the present paper. Loci within these putative inversions were also found to be associated with several environmental factors: summer temperature, precipitation, soil water availability, and soil nutrient content.

We lack population-level data to determine whether any inversions in the ‘other’ category are locally adaptive, but several might be, given high divergence between haplotypes. We also note that the inversions detected here are only the ‘tip of the iceberg’, as many more inversions will be revealed if the genomes of more individuals are sequenced and assembled.

### 3.3 Inversion frequency in sympatric versus allopatric species/subspecies

Existing literature on inversion establishment suggests that inversions can act as recombination modifiers to preserve sets of locally adaptive genes, and that they will be favoured at geographic locations where there is gene flow between heterogeneous environments (Kirkpatrick and Barton 2006). Evidence from our analyses suggests that there are more inversions detected in genetic clusters that include taxa with more geographic range overlap and that occupy more heterogeneous haplotypes.

The initial evidence comes from our analyses of locally adaptive inversions. Nine out of 11 of them were found among sympatric Texas populations belonging to different species/subspecies found in diverse habitats, including barrier islands, coastal beaches, and a variety of inland habitats. By contrast, only two locally adaptive inversions were found in Florida populations, where the subspecies are largely allopatric, and all are found on coastal beaches. This implies that local adaptation to heterogeneous environments in the presence of gene flow in Texas is more likely to favour polymorphic inversions.

The second layer of evidence lies in the number of ‘other’ inversions detected through assembly comparison. Haplotypes comparison within *H. debilis* ssp. *tardiflorus* revealed only 17 inversions >500 Kbp, whereas this number in *H. praecox* ssp. *hirtus* is more than tripled to 53 inversions. A caveat is that many of the ‘other’ inversions are likely to be near-neutral or directly adaptive, so their establishment fate would be unaffected by geographical relationships among populations. It is also possible that the *H. praecox* ssp. *hirtus* reference by chance combined more divergent haplotypes and thereby captured more inversions. But these haplotypes will still have to be present in a sympatric or parapatric system to be picked up in our sample, supported by the admixed signal in its sampled population PH1 (Figure 2d). Overall, our results support the hypothesis that a system with ongoing gene flow between populations in different habitats will accumulate more inversions, but additional comparisons in other species in this system and others are needed to confirm this observation.

Additionally, 75 ‘other’ inversions > 500 Kbp in length were found between the reference genomes for *H. praecox* ssp. *hirtus* and *H. debilis* ssp. *tardiflorus*. Future studies could further explore when these inversions arose and whether they represent fixed differences between species.

### 3.4 Characteristics of locally adaptive inversions

We found that adaptive inversions are longer, contain more genes, and exhibit higher levels of divergence compared to the ‘other’ inversions. This is consistent with predictions that selection favouring inversions that are indirectly adaptive will be stronger for inversions that contain more genes and suppress recombination between more genetically distant loci (Kirkpatrick and Barton 2006; Wellenreuther and Bernatchez 2018). Likewise, locally adaptive inversions are expected to be under balancing selection, possibly accounting for their greater divergence. On the other hand, we did not find that adaptive inversions occurred in higher recombination genomic regions as expected (Charlesworth and Barton 2017), possibly due to the low accuracy of our inferred recombination rate based on gene density. A caveat with these comparisons that the inversion categories were identified using different methods. While we employed thresholds to control for possible methodological biases, we cannot rule them out.

We also found that there are many more paracentric than pericentric inversions in the study system; this is not surprising as paracentric inversions are less strongly underdominant and are more likely to successfully establish (Coyne et al.1991). In addition, longer inversions were likely to be pericentric, whether found in the locally adaptive or ‘other’ inversion sets. Thus, our results show no evidence that pericentric inversions are more or less likely to contribute to local adaptation.

## 4. Conclusion

We conclude that three major genetic clusters are present in the wild sunflower system, comprising mixed eco-geographic populations of *H. debilis* and *H. praecox* in Texas and Florida, USA. Within this system, locally adaptive and other types of inversions support our first hypothesis that inversions are more likely to evolve in heterogeneous environments with ongoing gene flow than in allopatry. Our results further indicate that putatively locally adaptive inversions, compared with other types of inversions, are generally larger, contain more genes, and showed greater divergence between haplotypes. Collectively, these findings provide empirical evidence supporting the hypothesis that gene flow promotes inversion establishment and highlight that adaptive inversions are commonly associated with distinct genomic characteristics.

## 5. Materials and Methods

### 5.1 Taxonomy and geographic distributions

*Helianthus. debilis* consists of five subspecies, mostly found in Texas and Florida. *H. debilis* ssp. *debilis*, ssp. *vestitus,* and ssp. *tardiflorus* occur on the sandy beaches of east, west and northwestern Florida, respectively. The species have allopatric distributions, with ssp. *tardiflorus* further isolated from the other *debilis* subspecies by a flowering time barrier. The other two subspecies, *H. debilis* ssp. *silvestris* and *H. debilis* ssp. *cucumerifolius,* are endemic to eastern Texas, with some range overlap. Subspecies *silvestris* is known as forest sunflower, as it’s found in the sandy soil of pine-oak forests of northeastern Texas, whereas ssp. *cucumerifolius* occupies the open sand in southeastern Texas, where it is sympatric with *H. praecox*.

*Helianthus praecox* is comprised of three subspecies, which have parapatric distributions in southeastern Texas. *H. praecox* ssp. *praecox* is restricted to sandy soils around Galveston Island and the adjacent mainland; this subspecies is morphologically similar to *H. debilis* ssp. *vestitus* but is less densely pubescent (Heiser et al. 1969). Subspecies *hirtus* is rare and only occupies a small range in the sand soils of Carrizo Springs in southwestern Texas. Lastly, *H. praecox* ssp. *runyonii* can be found in the sandy, coastal prairies of southern Texas, where it intergrades phenotypically with *H. debilis* ssp. *cucumerifolius*, possibly due to hybridization (Heiser 1956; Heiser et al. 1969).

### 5.2 HiFi and Hi-C library preparation and sequencing

The individual used to assemble a haplotype-resolved reference genome for *H. praecox* was selected from *praecox* ssp. *hirsutus* (PI 468850) collected from southwest Texas. For *H. debilis*, one individual was selected from *debilis* ssp. *tardiflorus* (PI 468691), collected along the sandy beaches in northwest Florida. Geographic coordinates of both accessions can be found in Table S1. Germplasm was provided by USDA-ARS GeneBank and plants were grown at INRAE Toulouse, where leaf tissue was collected for high-molecular-weight DNA extraction (detailed protocol in Supplement Document 1). HiFi library preparation was done according to manufactural protocol for PacBio sequencing and sequenced on the PacBio Sequel II instrument by the Gentyane Platform (INRAE Clermont-Ferrand, France; Pubert et al. 2024). Hi-C library preparation protocols follow Hirabayashi et al. (2025). Given the high proportion of repetitive sequences in sunflower genomes, an enzymatic repeats depletion was performed on completed Hi-C libraries, similar to that used in Todesco et al. (2020). Briefly, Hi-C libraries were denatured, re-annealed at high temperatures (70 °C), treated with a duplex-specific nuclease (DSN; Evrogen, Mocow, Russia) to remove repetitive sequences (which are more abundant and can reanneal more efficiently), and then re-amplified. Hi-C reads were sequenced using a NovaSeq X machine at Canada’s Michael Smith Genome Science Centre.

### 5.3 Haplotype-resolved reference genome assemblies and quality checks

HiFi and Hi-C reads were quality filtered and were initially used to construct contig-level draft genome assemblies using hifiasm (v0.16.1; Cheng et al. 2021) with parameter “-s 0.4” and keeping default values for the remaining parameters. For *H. debilis,* Hi-C reads were mapped to the draft assembly using Juicer (v.9.9, Durand et al. 2016a) to transform raw reads into Hi-C contact positions. 3D-DNA (v80419, Dudchenko et al. 2017) was then used to scaffold mapped reads into a near-chromosome-level 3D de-novo assembly. For *H. praecox*, the above method resulted in overly fragmented assemblies for both H1 and H2. Therefore, we repeated the scaffolding step with another software, YaHS (v.2.2, Zhou et al. 2023), which is better at distinguishing real interaction signals from mapping noise; this resulted in a much more complete assembly. Scaffolded assemblies were then reviewed in Juicebox (v.1.11.08, Durand et al. 2016b) to generate a chromosome-level assembly for each haplotype. This review process included manual sorting and reorientation of contigs to resolve abnormal Hi-C contact signals.

To further align the order and orientation of the haplotype-resolved assemblies, we aligned both haplotypes for each individual using minimap2 (v2.28, Li et al. 2018), SyRI (v1.6, Goel et al. 2019) and plotSR (v1.0, Goel and Schneeberger 2022) to guide a second round of revision in Juicebox. Additionally, a third round of manual curation in Juicebox was performed to adjust any possible misassigned reads in each haplotype. To do so, we re-mapped the Hi-C reads to produce a haplotype-aware contact map including both haplotypes for each individual using 3D-DNA without misjoin correction (-r 0).

Genome assembly quality statistics were assessed using BUSCO (v6.0.0; Manni et al. 2021) in genome mode with the eudicotyledons_odb12 lineage dataset. The proportions of complete, duplicated, fragmented, and missing BUSCO genes were used to evaluate assembly completeness. Candidate intact LTR retrotransposons were identified using LTRharvest and LTR_FINDER_parallel following the recommended LTR_retriever pipeline. The curated intact LTR retrotransposon set was subsequently used to calculate genome-wide LTR Assembly Index (LAI) scores using LTR_retriever (v2.9.0.; Ou and Jiang 2018; Ou et al. 2018).

A small portion of contigs were not scaffolded into the final genome assembly; these unplaced contigs mainly consist of repeats (see below). We named the finalized haplotype reference genomes as HDebH1 (H1: haplotype 1), HDebH2 (H2: haplotype 2), HPraH1 and HPraH2.

### 5.4 Genome annotation, centromere identification, and visualization

We generated gene and repeat annotations, and identified centromeres, on the finalized genome assemblies. Transposable elements and other repetitive sequences were annotated using a *de novo* repeat identification strategy. RepeatModeler (v2.0.7; Flynn et al. 2020) was used to construct species-specific repeat libraries with the LTRStruct pipeline enabled, and the resulting consensus libraries were subsequently used by RepeatMasker (v4.1.9; Smit et al. 2015) to annotate and soft-mask repetitive regions in each genome assembly and in the unassigned contigs separately. Additionally, we identified putative centromeres using RepeatOBserver (Elphinstone et al. 2025) inferred by “MinRepeatAbund”. These predicted centromeres corresponded to regions enriched in sunflower-specific centromeric repeats that had been identified though ChIP-seq of CEN3 histones in cultivated sunflower (*H.* a*nnuus*; Nagaki et al. 2015).

Gene annotation was performed using BRAKER (v3.0.6; Gabriel et al. 2024) with combined transcriptomic and protein homology evidence for each genome assembly and the unassigned contigs separately. Illumina RNA-seq reads (described in Table S4) were aligned to the reference genomes using STAR (v2.7.11b; Dobin et al. 2013) and long-read transcriptomic sequences were aligned using minimap2 (v2.26; Li 2018, 2021). The resulting RNA-seq alignments, together with Viridiplantae protein sequences as external protein evidence, were supplied to BRAKER for gene prediction. Genome assemblies with soft masked repeats were used as input for annotation. Gene annotation completeness was assessed using BUSCO (v6.0.0; Manni et al., 2021). Predicted protein sequences generated by BRAKER3 were used as input, and the proportions of complete, duplicated, fragmented, and missing BUSCO genes were used to evaluate annotation completeness.

Visualization of the above annotations between haplotypes in a Circos plot (Krzywinski et al. 2009) were generated using TBtools-II (v2.441, Chen et al. 2023).

### 5.5 Population-level plant material

Seeds of 47 populations from three subspecies of *H. praecox* and five subspecies of *H. debilis* were provided by the USDA-ARS GeneBank (Figure 2a, coordinates in Table S1). Seeds were germinated in the lab and grown in a soil mix composed of 3 parts of potting mix, 2 parts of medium grit sand, and 1 part of medium perlite in the University of British Columbia greenhouses in Vancouver, Canada. Young leaves from up to 10 plants per population were collected for DNA extraction.

### 5.6 GBS library preparation and sequencing

DNA for each individual was extracted using a modified CTAB protocol (Zeng et al. 2002) and used to prepare genotype-by-sequencing (GBS) libraries, for a total of 451 individuals: 130 *H. praecox* samples from 13 populations and 321 *H. debilis* samples from 34 populations. GBS libraries were prepared using a standard two-enzyme protocol (Poland et al. 2012), except for the addition of a duplex-specific nuclease (DSN) treatment, which reduces the proportion of high-copy fragments (Moyers et al. 2017). The library was then sequenced to an average of 1.89 million reads per individual by Novogene using Illumina sequencing technology.

### 5.7 Alignments, variant calling, and downstream filtering

To generate each set of single nucleotide polymorphisms (SNPs), raw GBS reads were first aligned against the newly assembled reference genomes (see section 2.3) using bwa-mem2 (v2.2.1, Li et al. 2013). Then variant calling was performed with the Genome Analysis Tool Kit (GATK; v4.2.4.0; DePristo et al. 2011) followed by the following variant filtering pipeline. First, sites and samples with low sequencing quality were filtered with gatk VariantFiltration “QD < 2|| FS > 60|| MQ < 40|| MQRankSum < −12.5|| ExcessHet > 54.69” for site level filter, and with bcftools filter “GQ < 30||DP <10” to filter samples with genotype quality lower than 30 and sequencing depth lower than 10. Then, we further removed sites for which alternative alleles were not present after low-quality samples; these sites, along with multi-allelic sites and sites with missing data over 30%, were removed with gatk SelectVariants “--remove-unused-alternates, --restrict-alleles-to BIALLELIC and --max-nocall-fraction 0.3”. Samples with variant coverage (vcftools --missing-indv) missing more than 30% were removed using awk. Selected fields were recoded, SNPs with minor allele frequency <3% were filtered out, with the remaining polymorphic SNPs retained in VCF format for downstream analyses.

We initially aligned reads from all 451 samples to the HPraH2 genome assembly and called a global SNP set using the pipeline described above. Thirty-five samples with missing data >30%, or which did not cluster with other individuals from the same population, were removed (see below; Table S5). We then re-called SNPs for the remaining 416 samples. This data set was used to determine the optimal number of genetic clusters in this system, which was found to be three (see Results). Samples were placed into three sets based on the clustering results (A, B, and C), and SNPs were called independently for each set to reduce reference bias. Set A included allopatric *H. debilis* subspecies in Florida and was called on the HDebH2 genome assembly. Sets B and C were comprised of the remaining subspecies of *H. praecox* and *H. debilis* from Texas and were called on the HPraH2 genome assembly. For the complete list of samples used in each SNP set, see Table S5.

### 5.8 Population structure and principal component analyses

To understand population structure within each SNP set, we pruned out highly correlated SNPs based on LD patterns and physical proximity with PLINK (v1.9, Purcell et al. 2007) “--indep-pairwise 10kb 50 0.2” and kept only one SNP for every 300 bp. ADMIXTURE (v.3.0; Alexander et al. 2009) was used to examine population structure among individuals for the global SNP set, with the number of genetic clusters K ranging from 1 to 13. The most suitable number of genetic clusters was determined by the *K* value with the lowest cross-validation error.

Principal Component Analysis (PCA) was also conducted to further evaluate the genetic structure using snprelate in R 4.2.1 (Zheng et al. 2012; R Core Team 2022). PCA was used to identify individuals that did not cluster with the population they were sampled from, which were then removed. VCFs were updated and separated to minimize population structure while also considering the optimal biological clusters for downstream haploblock detection.

### 5.9 Identification of locally adapted inversions

Haploblocks were detected following Huang et al. (2020) using the local population structure (*lostruct*) software package developed by Li and Ralph (2019). This approach conducts PCA on non-overlapping 50-SNP windows across the genome. Regions of the genome that contribute to local adaptation are expected to have a different population structure than the rest of the genome. If structural variants such as inversions contribute to local adaptation, then numerous linked windows across the rearranged region are expected to show the same outlier pattern. We kept 40 multidimensional scaling (MDS) coordinates and windows with greater than 3 standard deviations (SD) from the across-window mean; adjacent outliers with equal or fewer than four windows between them were kept as a putative cluster. Highly correlated outliers on the same chromosome across multiple MDS coordinates were detected based on Pearson’s correlation coefficient calculated using the genotype matrix; when the coefficient was greater than 0.8, these were collapsed by selecting regions with the larger number of outliers. As linked selection or other evolutionary processes may cause similar patterns, we conducted additional analyses on outlier regions following Todesco et al. (2020). If the haploblocks are caused by structural variants, then a PCA of all SNPs within the outlier regions should display three distinct clusters corresponding to the homozygous genotypes for each haplotype and heterozygotes between the haplotypes. The latter should form an intermediate cluster and have the highest heterozygosity. We did not examine patterns of linkage disequilibrium (LD) within haploblocks due to the limited number of SNPs in GBS data. In addition, we asked whether the *lostruct*-detected haploblocks could be confirmed as inversions through coordinate alignment with genome assembly comparisons. Most *lostruct* outlier regions were confirmed as inversions (see Results), so those haploblocks will be referred to as putative locally adaptive inversions in the next sections.

We further tested if any loci within the putative inversions detected above were also associated with environmental variables, which could potentially account for the local adaptation signal. Environmental data included climate data extracted from ClimateNA (v7.70; Wang et al. 2016) using the ClimateNAr R package (v3.1.0; Schönberger and Wang 2025). Because sunflower is an annual plant, we included 19 annual climate variables and 25 seasonal climate variables. Our selected sunflower populations are also thought to be adapted to different edaphic conditions. Therefore, we extracted 15 soil variables at soil depths between 0 cm and 180 cm from SSURGO using the soilDB R package (v2.9.1; Beaudette et al. 2026). A few populations in Florida lacked soil data due to their proximity to the ocean and recent site modification or submergence due to rising sea levels (Table S1).

All loci were further filtered to retain only MAF >5% to compute allele frequencies per population. For each climate and soil variable, regularized least-squares estimates for Latent Factor Mixed Model (LFMM) using a ridge penalty were calculated with the *lfmm* package (Caye et al. 2019) in R 4.5.0 (R Core Team 2025). The best number of latent factors was inferred by *K* with the lowest cross-validation error in ADMIXTURE, with adjustment to maintain the genomic inflation factor (git) scores close to 1 to ensure residual population structure are not inflating best candidate loci detection using LFMM. This was done because while *K* with the lowest cross-validation error may explain the overall genetic structure, it may over or under-correct for other structure present in the dataset. All putative locally adaptive inversions identified with local PCA were checked to determine if they harboured environmental adaptive loci detected in LFMM. Allele frequencies of selected adaptive inversions were plotted along significant environmental factors as a monotonic fitted line with spearman’s rho of increasing or decreasing relationship.

### 5.10 Identification of ‘other’ inversions

We identified many other inversions by comparing genome assemblies, including between haplotypes (HDebH1 - HDebH2 and HPraH1 - HPraH2) and between species (HDebH2 - HPraH2). Genome assembly comparison for syntenic regions and structural variation identification was conducted at both the nucleotide level and the homologous gene level. The former was generated with the combination of minimap2 and SyRI visualized by plotSR, and the latter was conducted with a combination of Liftoff (Shumate and Salzberg 2021) for homologous gene detection and then visualized using GENESPACE (v1.4; Lovell et al. 2022). Liftoff was used to map the previously annotated HA412v2 genome (Huang et al. 2023), consisting of 44,544 genes, to the newly assembled genomes described herein.

The following comparisons between putative locally adapted inversions and ‘other’ inversions were restricted to inversions of above 500 Kbp. This was done to reduce bias because *lostruct* has low power to detect haploblocks under 400 Kbp using our GBS data and selection criteria.

### 5.11 Comparison of locally adaptive and “other” inversions

#### 5.11.1 Length and number of genes

Inversion length and the number of genes they contain were calculated based on their coordinates in the HDebH2 and HPraH2 assemblies. For comparisons between these two genome assemblies, HPraH2 coordinates were employed. Due to differences in the numbers of locally adapted versus ‘other’ inversions, and the non-normal distribution and presence of outliers, we selected the Wilcoxon rank-sum test to identify significant differences between the inversion categories.

#### 5.11.2 Level of divergence between haplotypes

Our population-level GBS data are sparse and cannot provide a robust measure of divergence for all identified inversions. Therefore, divergence was calculated using the haplotype-resolved reference genomes. First, inversion positions were determined based on their coordinates in the HDebH2 and HPraH2 assemblies. Then, we compared HDebH1-HDebH2 and HPraH1-HPraH2 to find syntenic regions using minimapV2 and SyRI. We used pairwise nucleotide diversity (π) as a measure of divergence between the haplotypes for each candidate inversion. Pairwise nucleotide diversity was calculated with the number of SNPs as the numerator, and the length of the aligned region corresponding to the inversion as the denominator. If the total aligned length was less than 100 Kbp or if no SNPs were found, the inversion was excluded from Wilcoxon rank-sum test for differences in levels of sequence divergence between the ‘locally adapted’ and ‘other’ categories of inversions. Note that for this comparison we excluded inversions that were polymorphic between species only, since in these cases, divergence between inversion haplotypes would be confounded by between-species divergence.

#### 5.11.3 Recombination proxy from gene density

Recombination rate data were not available for the assembled sunflower species. However, numerous recombination maps have been made for a closely related species, *Helianthus annuus*, and average recombination rates have been calculated on a 1 Mbp scale for the HA412 genome (Huang et al. 2023). Using this map, we calculated the nucleotide collinearity between HA412 relative to HDebH2 and HPraH2, respectively, using minimap and SyRI to explore the possibility of transferring the recombination rate between collinear regions; however, this attempt failed because of numerous structural differences between the genomes (see Figure S14 and S15). We then tested if gene density could predict recombination rate at a 1 Mbp scale in HA412. A relatively strong correlation (r = 0.618, p-value <2.2e-16; Figure S16) was found, so we used this relationship and known gene density in 1 Mbp windows to predict recombination rates for HDebH2 and HPraH2 (Figure S17, S18). Finally, we summarized recombination rates for the regions containing inversions.

#### 5.11.4 Paracentric and pericentric inversion comparison

Centromere positions predicted by RepeatOBserver were used to classify whether the identified inversions were paracentric or pericentric. We hypothesize that larger inversions are more likely to overlap a centromere (pericentric) by chance. Therefore, we used a logistic regression to test whether inversion type and inversion length predicted the probability that an inversion was pericentric. The model included inversion type and log-transformed inversion length as predictors, with pericentric status treated as a binary response variable.

## Data Availability

All raw sequences are deposited to xxx (ongoing). The assembled reference genomes are uploaded to NCBI at xxx (ongoing). Code for all the analyses for this manuscript can be found at https://github.com/yueyu27/debilis_praecox_project.

## Acknowledgment

We thank Natalia Bercovich for training on DNA extraction and GBS library preparation lab work; Kaede Hirabayashi for support with Hi-C library depletion; Cassandra Elphinstone for support with RepeatObserver and detection of centromere regions; Zhe Cai for insightful discussions on inversion signals; Kaichi Huang for discussions on the application of local PCA; Gregory Owens for discussions on recombination rates and related topics; Tom Booker and Fernando Hernandez for guidance on the use of the GEA approaches; Tongli Wang for clarifying questions regarding seasonal climate variables in ClimateNA; Laura Marek for assistance in clarifying information related to requested seed accessions and their geographic origins; Alan Stahnke and Josué Aceituno-Díaz for guidance on soil variable selection; and Yu-Xuan Ye for the illustration of the two reference genome species. We also thank the Centre National de Ressources Génomiques Végétales (CNRGV) and the GeT-PlaGe platform for HiFi library preparation and sequencing.

Funding was provided by China Scholarship Council (CSC-202108180006) to Y.Y. and the Natural Sciences and Engineering Research Council of Canada (RGPIN-2022-03002) to L.H.R. Sequencing of the assembled reference genomes was funded by the International Consortium on Sunflower Genomics (ICSG).

## Supplementary Table Captions

**Table S1**: Location, population and sequencing information on all sequenced populations and reference genomes.

**Table S2**: Latent Factor Mixed Model (LFMM) outlier SNPs and its corresponding adaptive inversions for genetic cluster A, B, C, and B plus C.

**Table S3**: Inversions identified with SyRI for comparisons of HDebH1 - HDebH2, HPraH1 - HPraH2 and HPraH2 - HDebH2. Rows in red are after filtering for inversions >500 Kbp.

**Table S4**: Illumina RNA sequence information.

**Table S5**: Complete list of samples used in each SNP set.

## Supplementary Document Caption

**Supplement Document 1:** Protocol details on extraction of High Molecular Weight (HMW) DNA extraction and quality check for the two reference genomes.

## Supplementary Figure Captions

**Figure S1:** Admixture plot K = 1 to 12 for all 416 samples. Each column representing an individual, with each corresponding population ordered from west to east.

**Figure S2:** Inversion plot for deb10.01

**Figure S3:** Inversion plot for deb12.01

**Figure S4:** Inversion plot for pra11.01

**Figure S5:** Inversion plot for pra17.01

**Figure S6:** Inversion plot for pra17.02

**Figure S7:** Inversion plot for pra08.01

**Figure S8:** Inversion plot for pra08.02

**Figure S9:** Inversion plot for pra05.01

**Figure S10:** Inversion plot for pra17.03

**Figure S11:** Inversion plot for pra08.03

**Figure S12:** Inversion plot for pra11.02

**Figure S13:** GENESPACE plot between species: HDebH2 - HPraH2

**Figure S14:** Syntenic regions and structural variations between HA412 and HDebH2

**Figure S15:** Syntenic regions and structural variations between HA412 and HPraH2

**Figure 16:** Correlation between number of gene count (x axis) and recombination rate (y axis) for Ha412. Correlation shown 0.618 with p-value < 2.2e-16.

**Figure 17:** Recombination rate for HDebH2 inferred from gene density for all 17 chromosomes, predicted based on the recombination rate – gene density relationship from HA412.

**Figure 18:** Recombination rate for HPraH2 inferred from gene density for all 17 chromosomes, predicted based on the recombination rate – gene density relationship from HA412.

