## Supplementary Figure S1-S18 for "Genomic and eco-geographic features of locally adapted inversions in wild sunflowers"

Figure S1: Admixture plot K = 1 to 12 for all 416 samples. Each column representing an individual, with each corresponding population ordered from west to east.

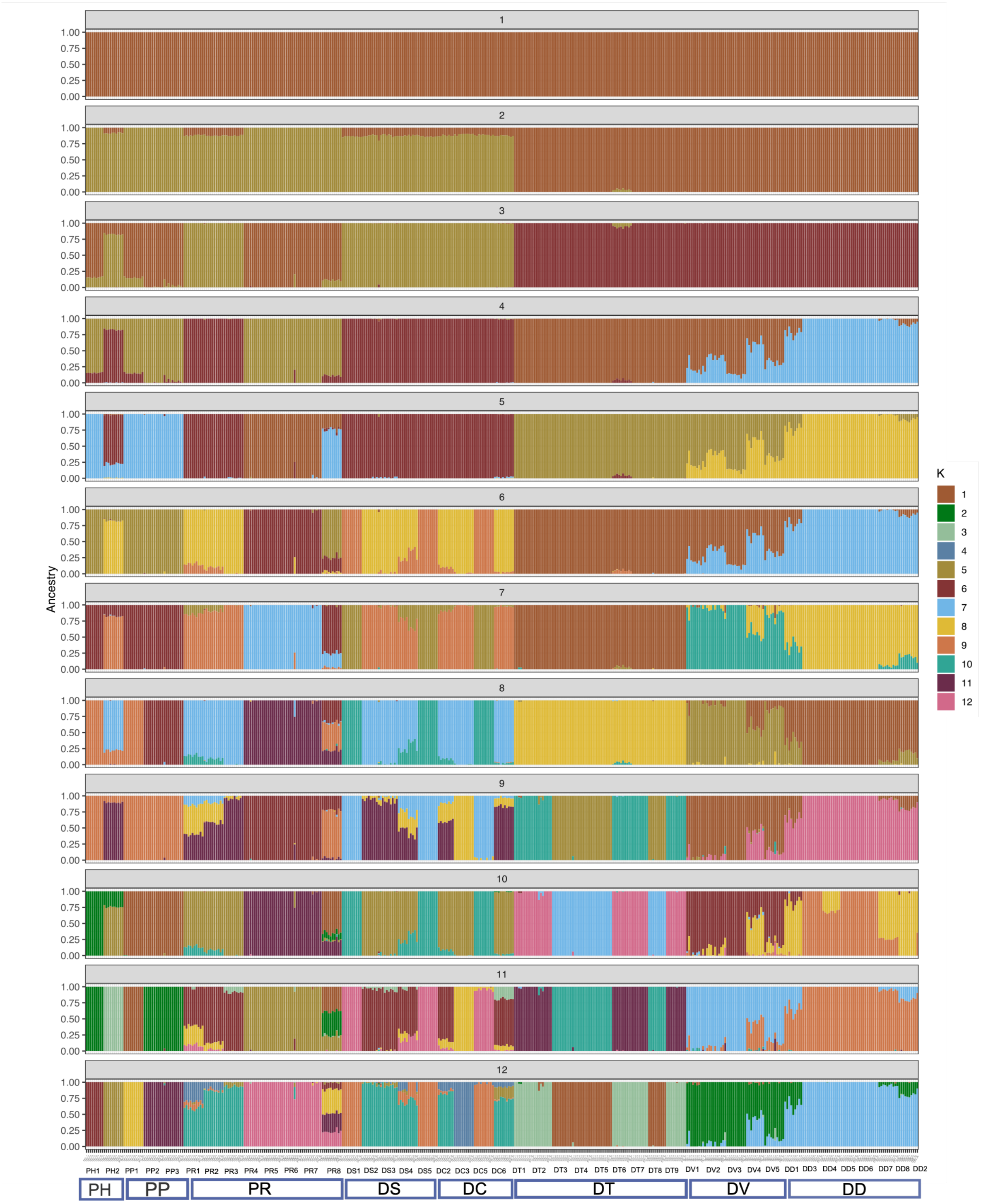

Figure S2: Inversion plot for deb10.01

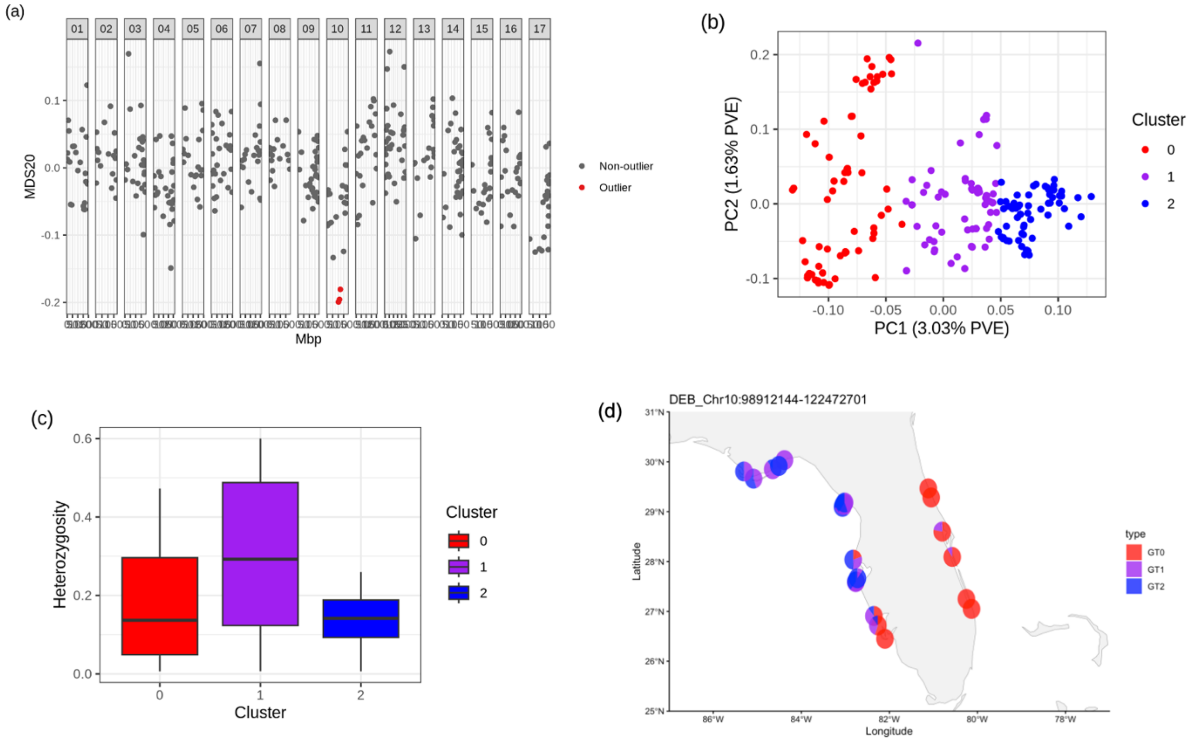


Figure S3: Inversion plot for deb12.01

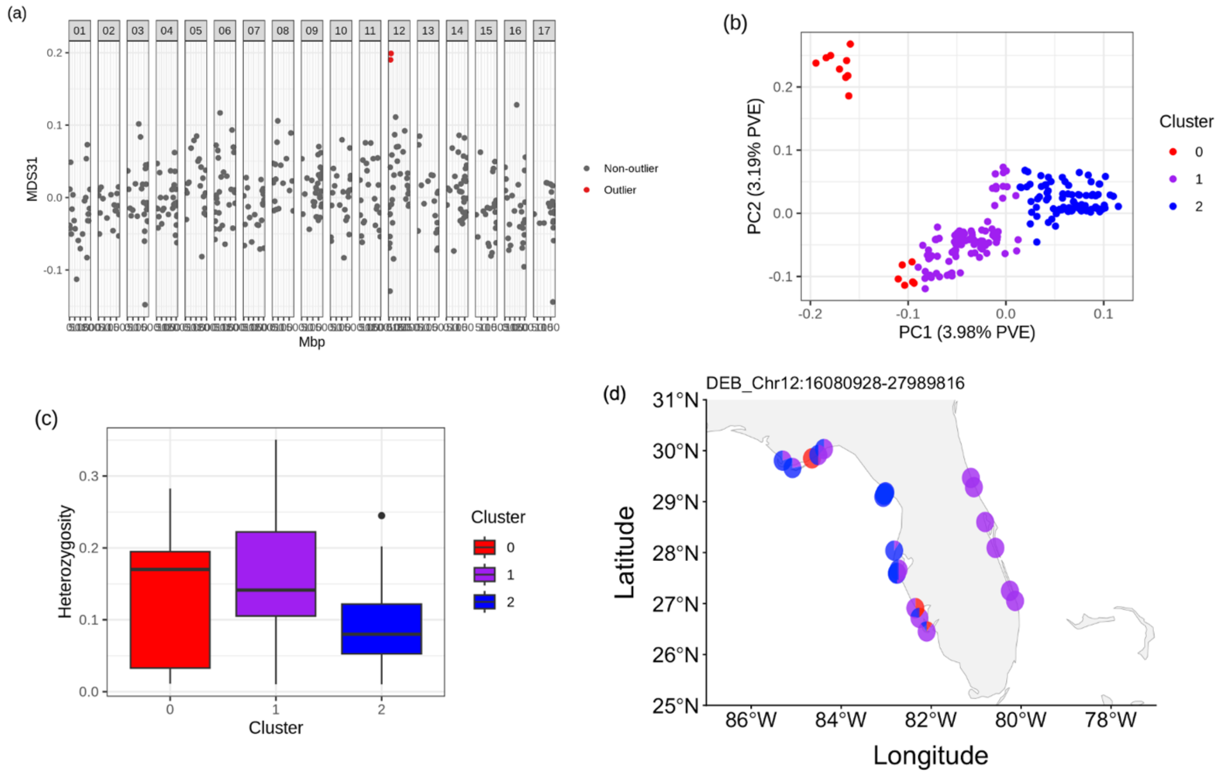

Figure S4: Inversion plot for pra11.01

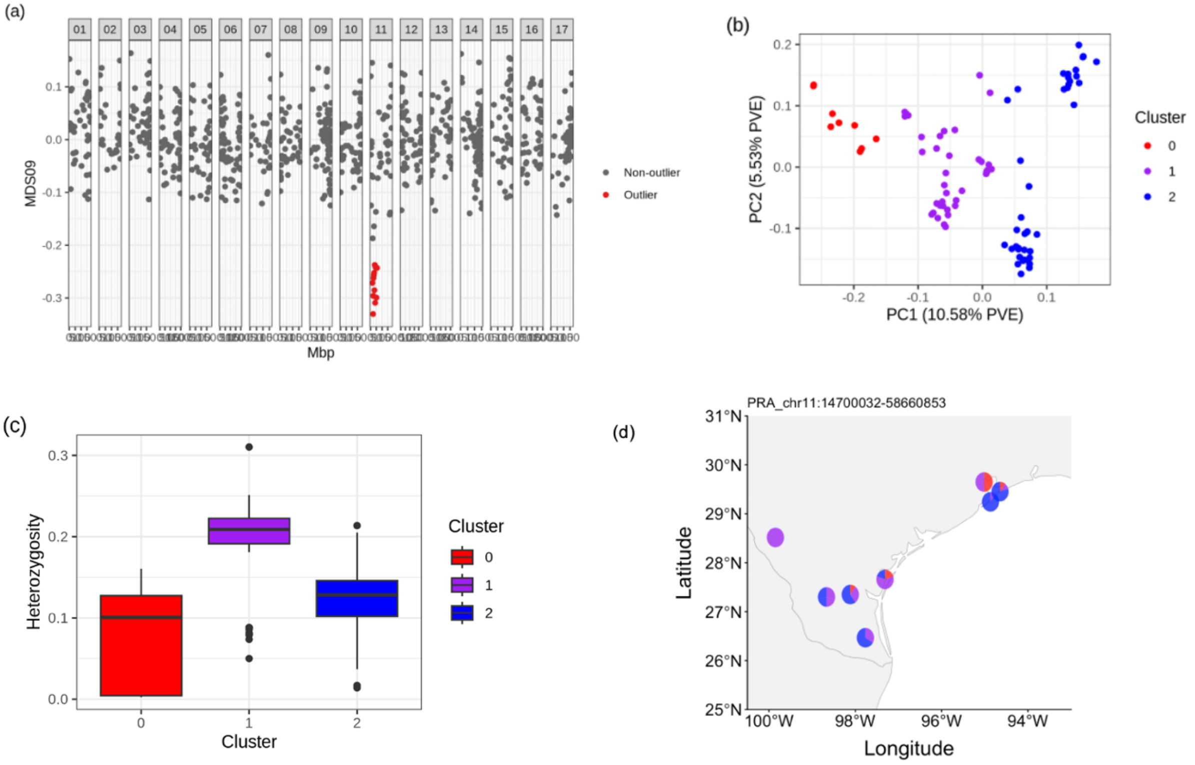


Figure S5: Inversion plot for pra17.01

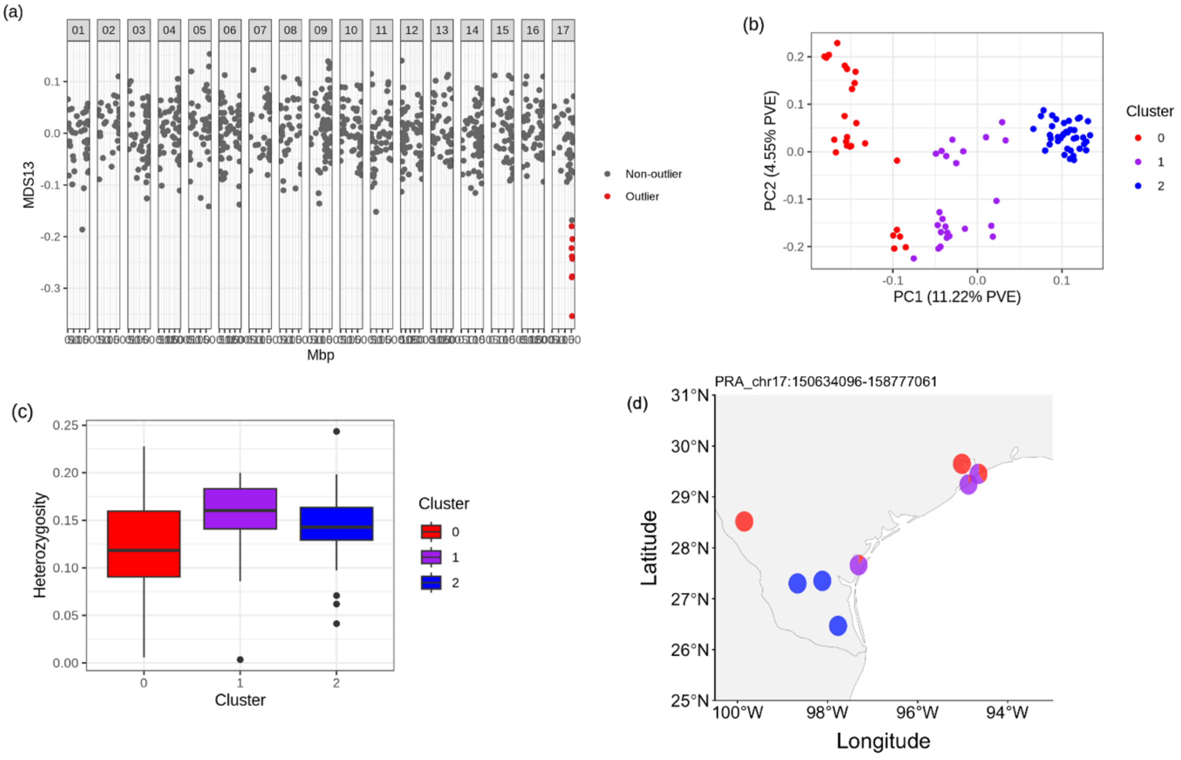

Figure S6: Inversion plot for pra17.02

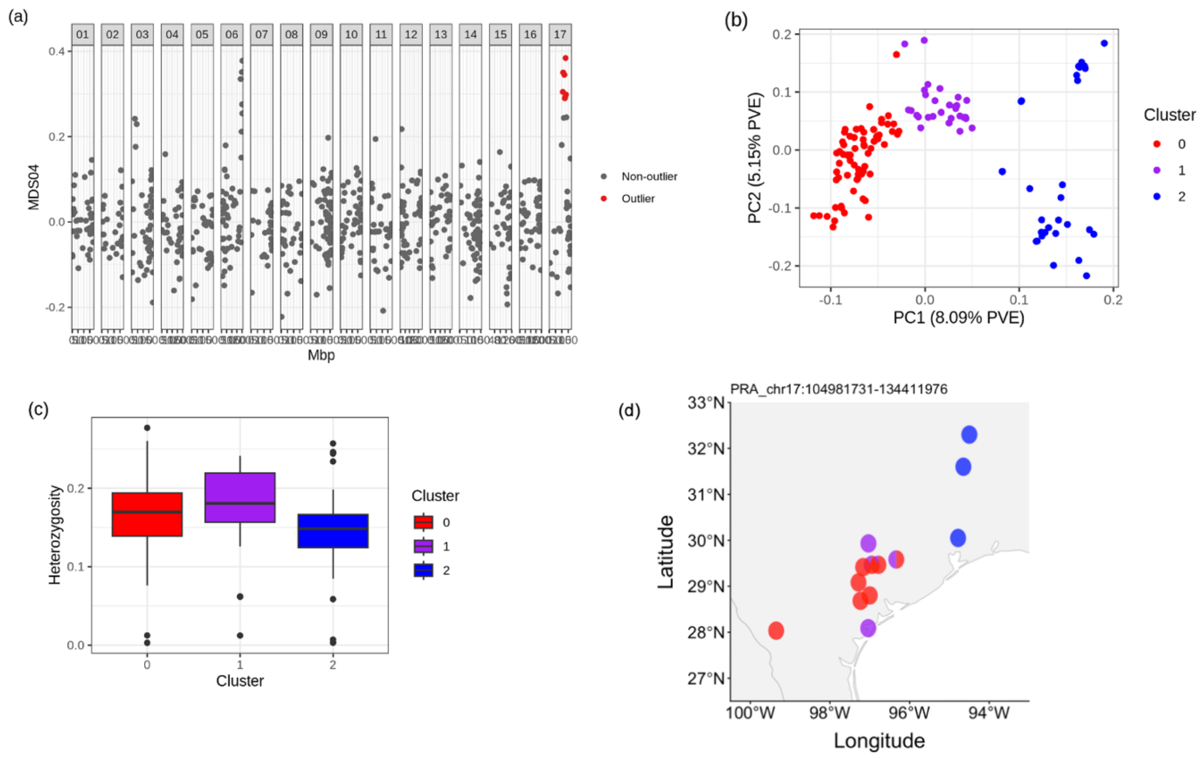


Figure S7: Inversion plot for pra08.01

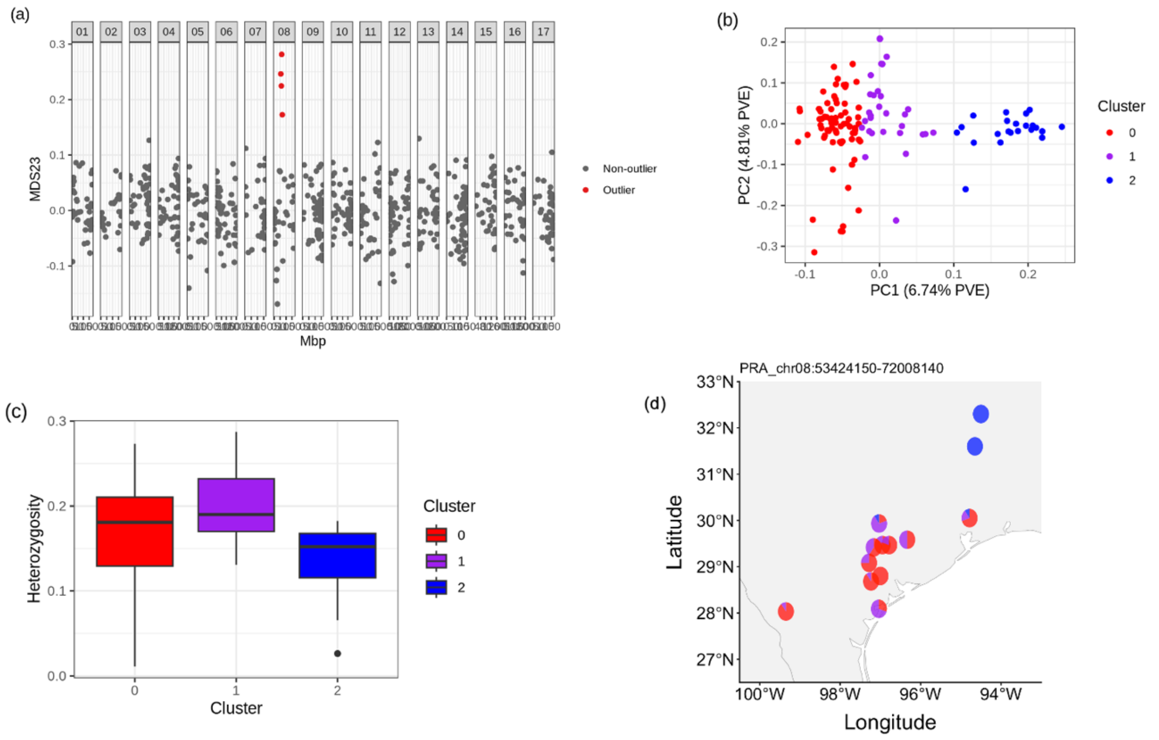

Figure S8: Inversion plot for pra08.02

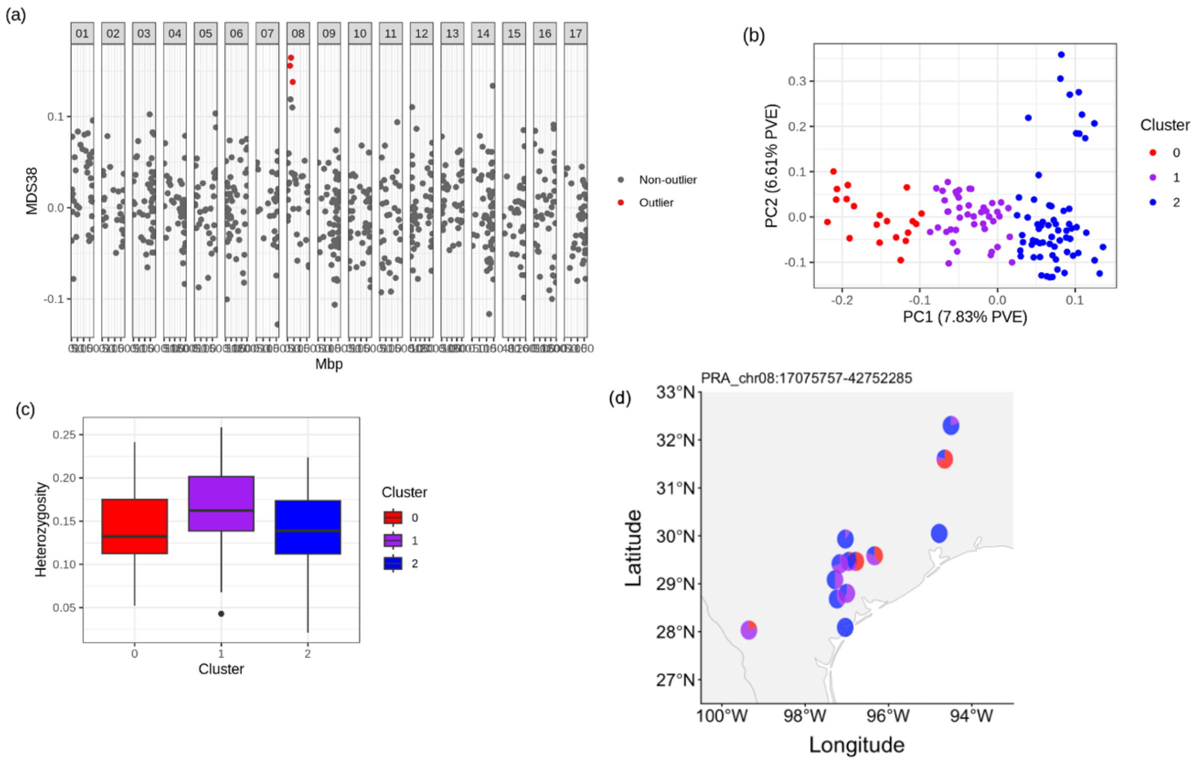


Figure S9: Inversion plot for pra05.01

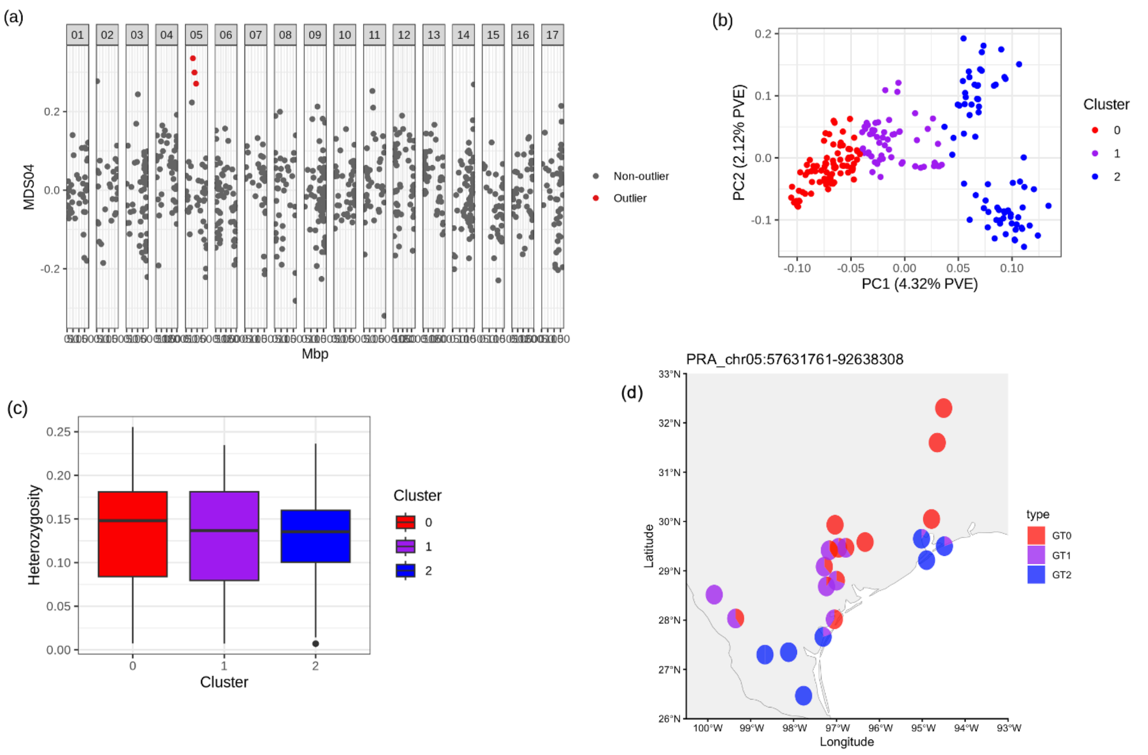

### Figure S10: Inversion plot for pra17.03

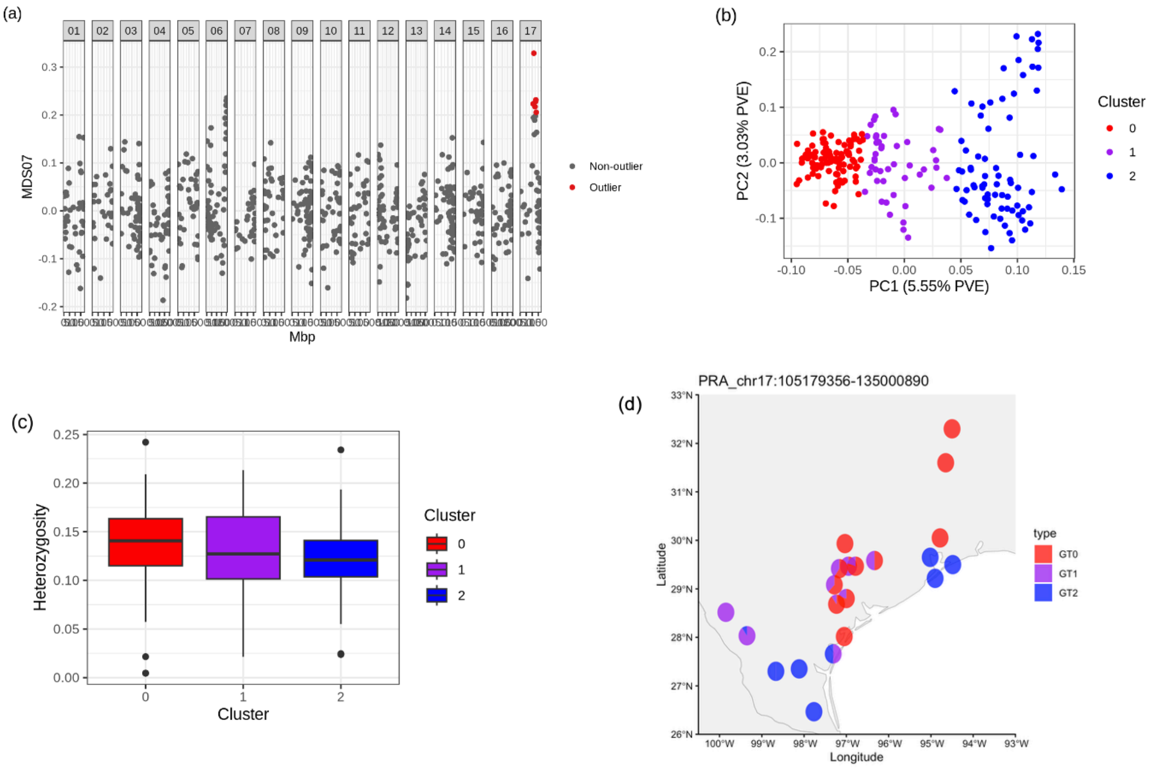


### Figure S11: Inversion plot for pra08.03

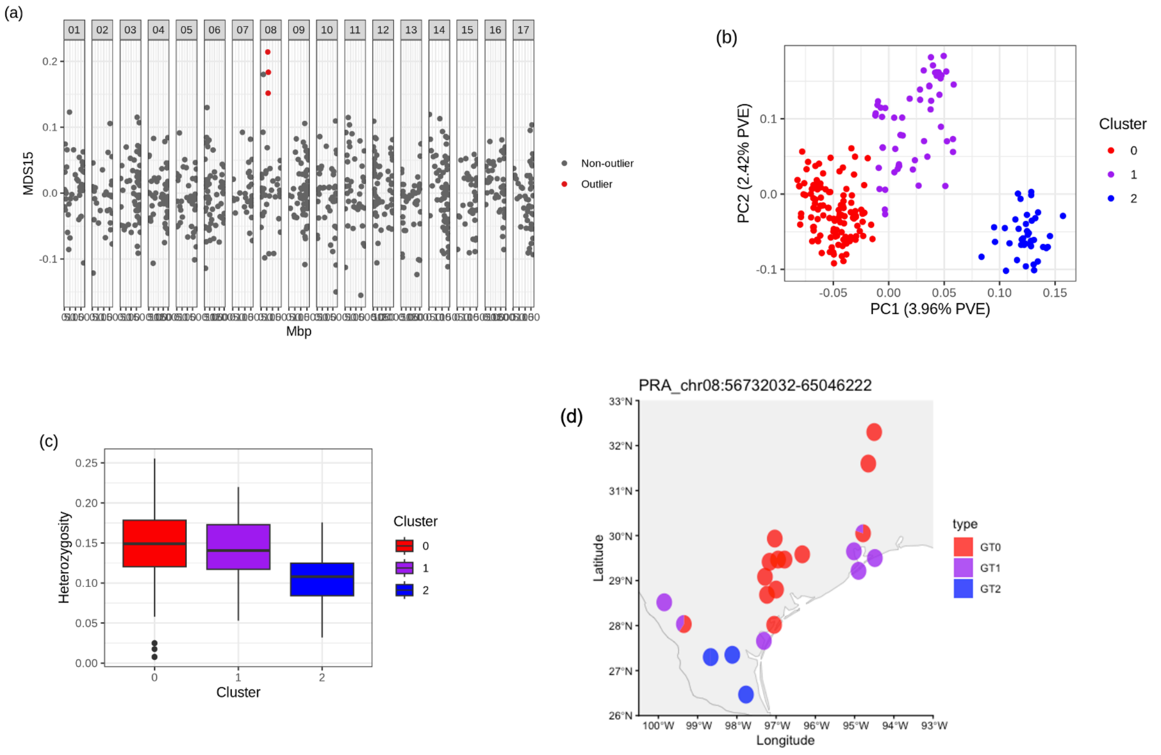

Figure S12: Inversion plot for pra11.02

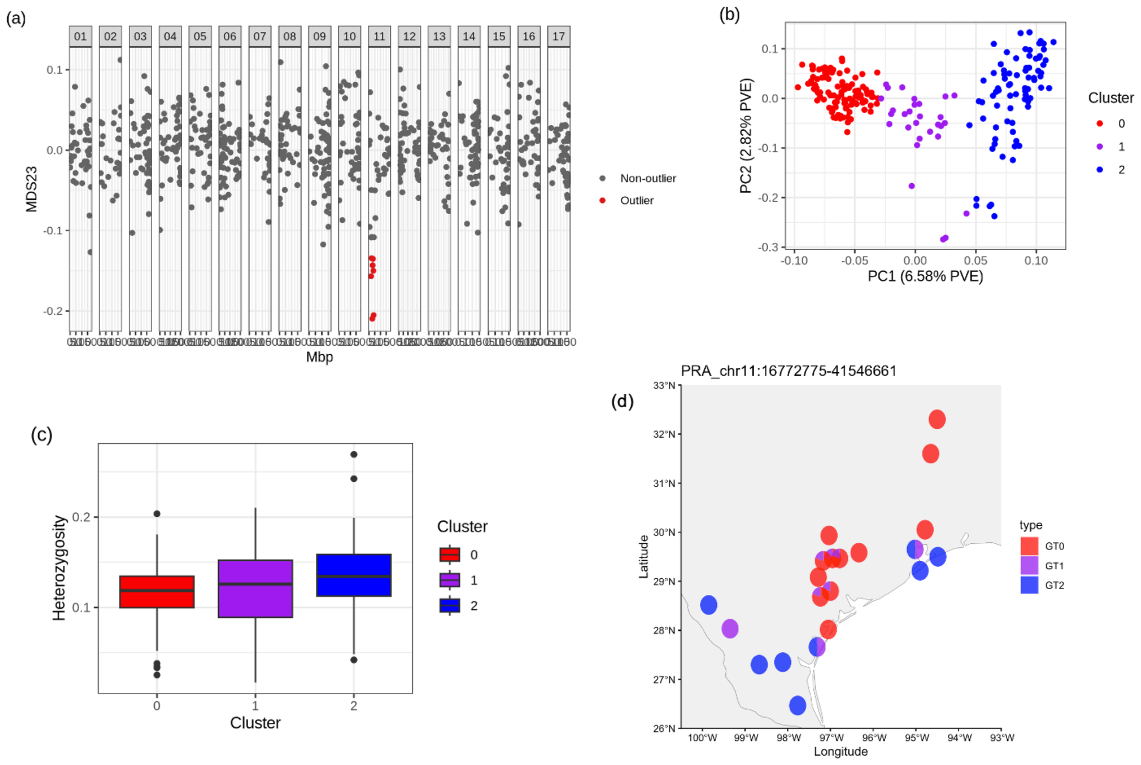


Figure S13: GENESPACE plot between species: HDebH2 - HPraH2

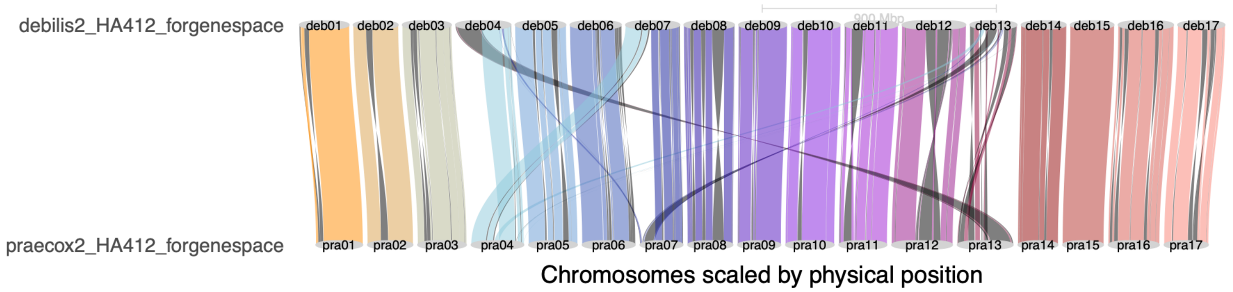

Figure S14: Syntenic regions and structural variations between HA412 and HDebH2.

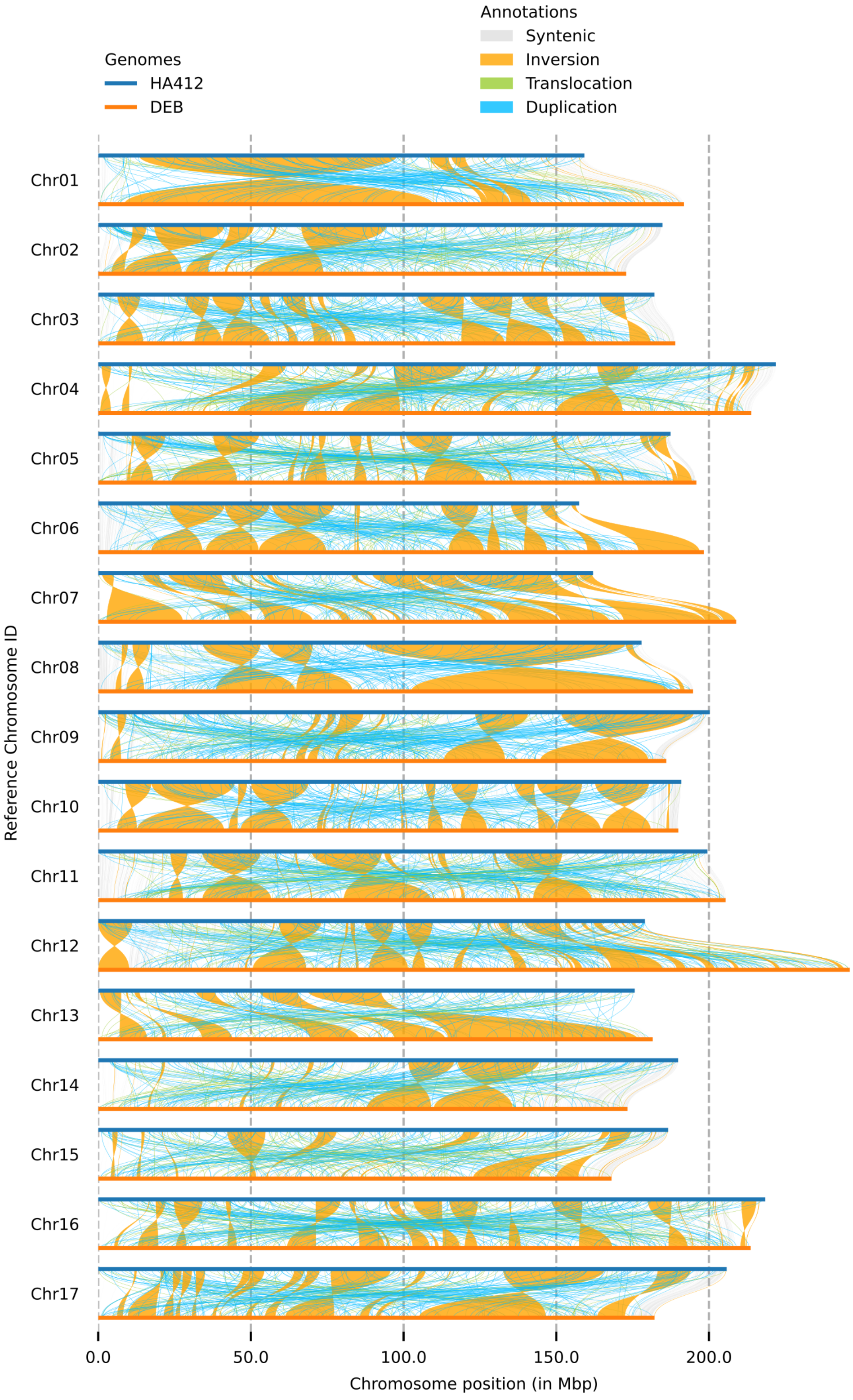

Figure S15: Syntenic regions and structural variations between HA412 and HPraH2.

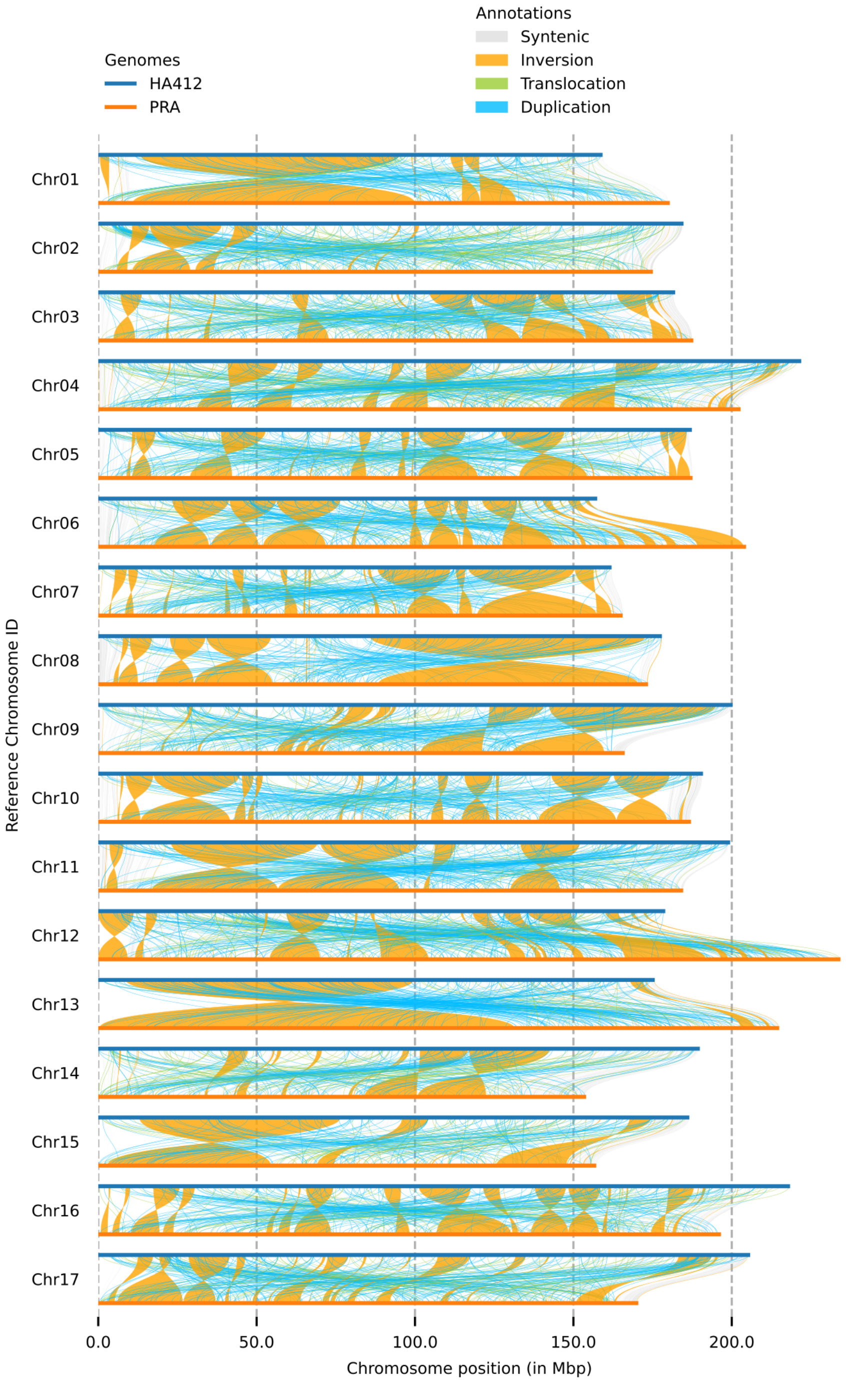

Figure 16: Correlation between number of gene count (x axis) and recombination rate (y axis) for Ha412. Correlation shown 0.618 with p-value < 2.2e-16.

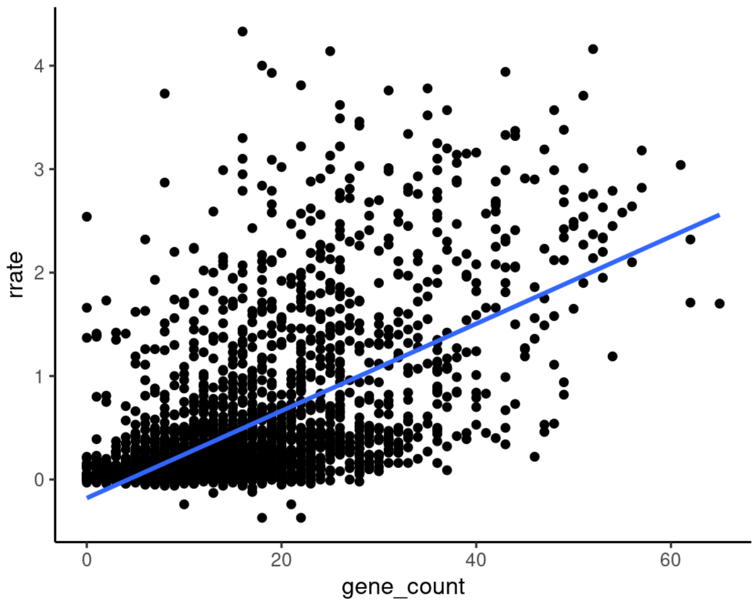

Figure 17 Recombination rate for HDebH2 inferred from gene density for all 17 chromosomes, predicted based on the recombination rate – gene density relationship from HA412.

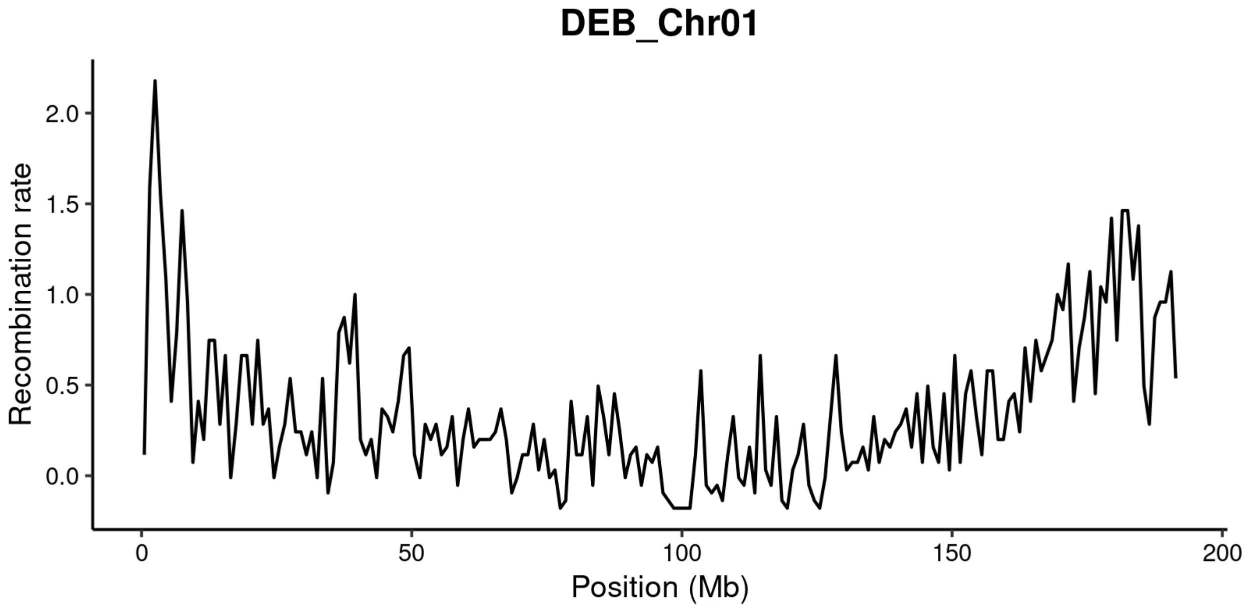


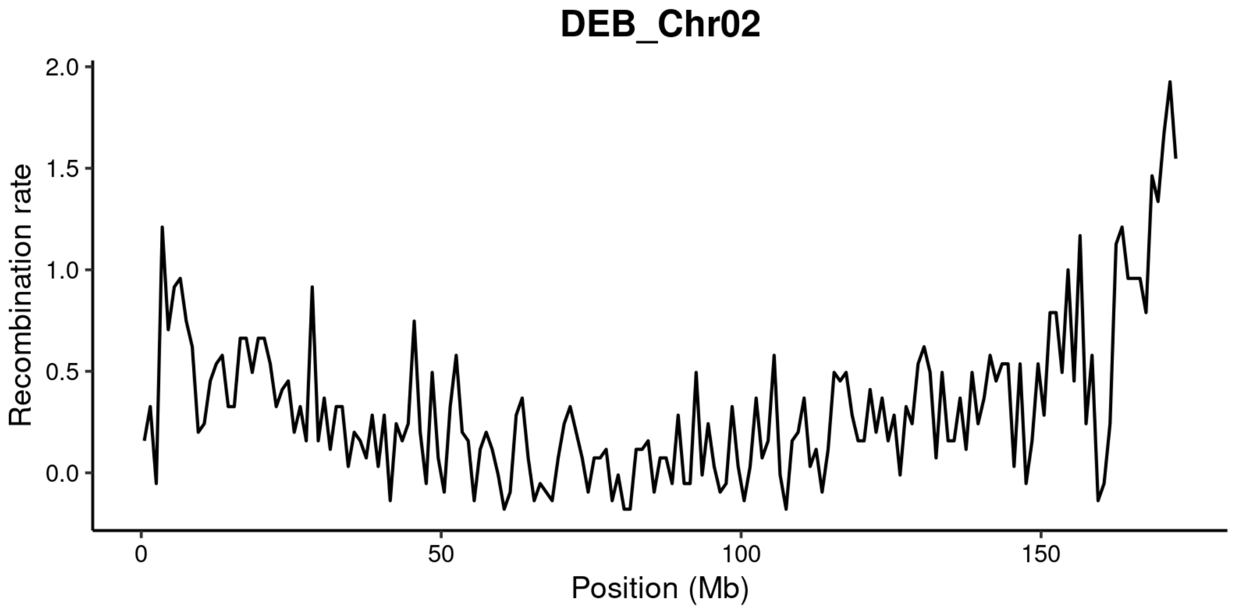

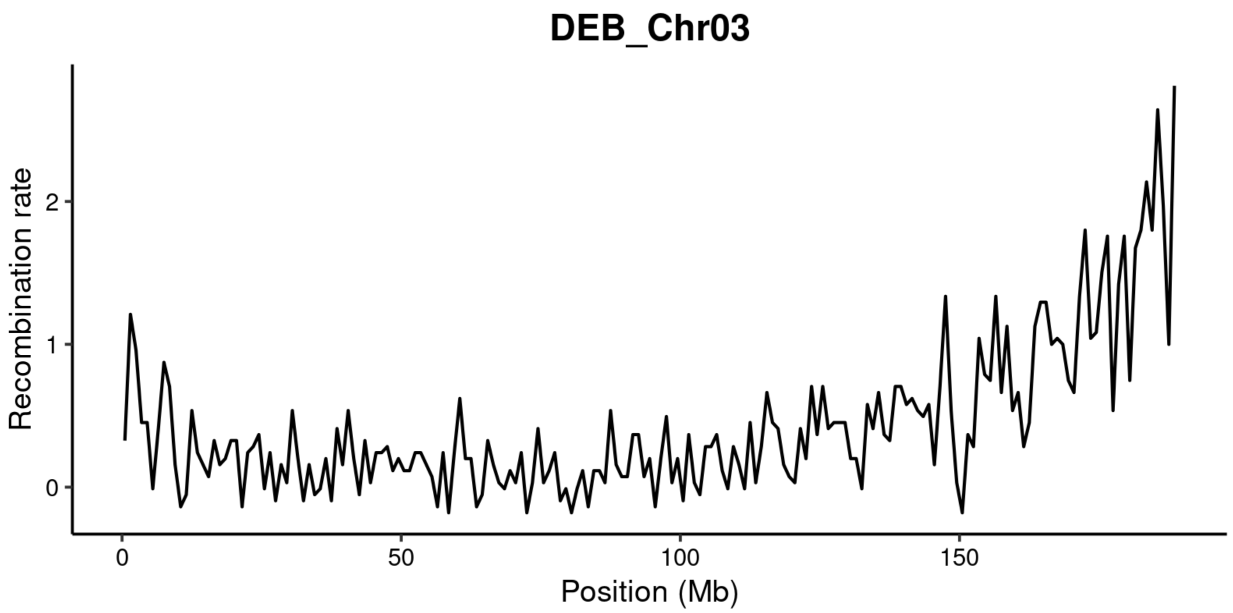
DEB_Chr03
Recombination rate
0
1
2
0
50
100
150
Position (Mb)

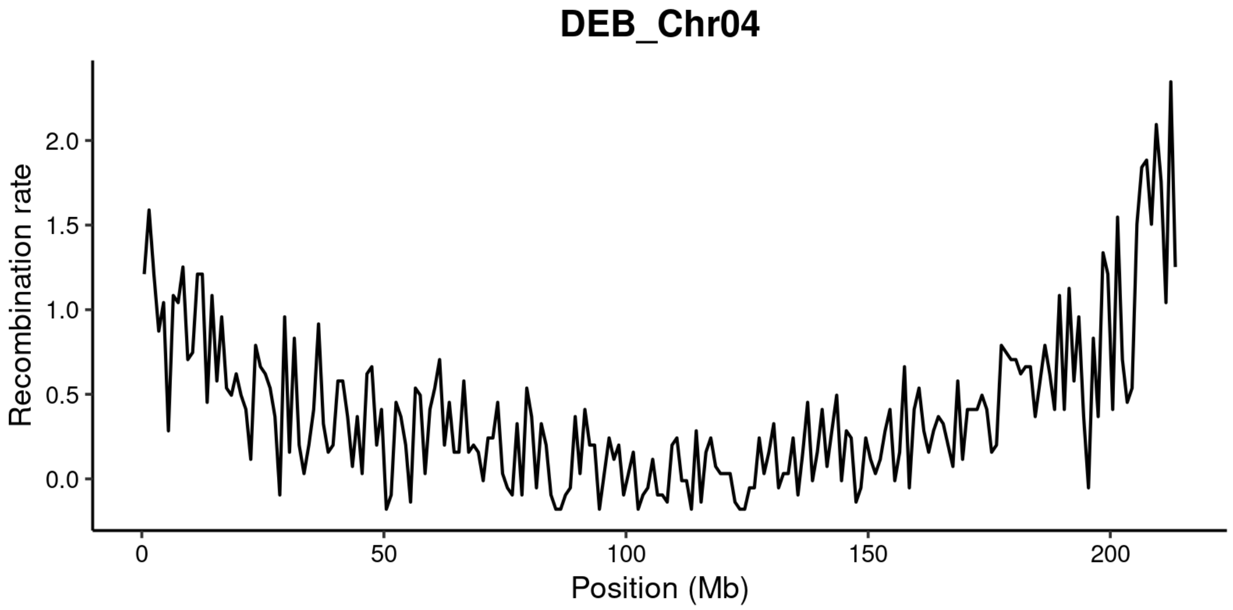
DEB_Chr04
Recombination rate
0.0
0.5
1.0
1.5
2.0
0
50
100
150
200
Position (Mb)

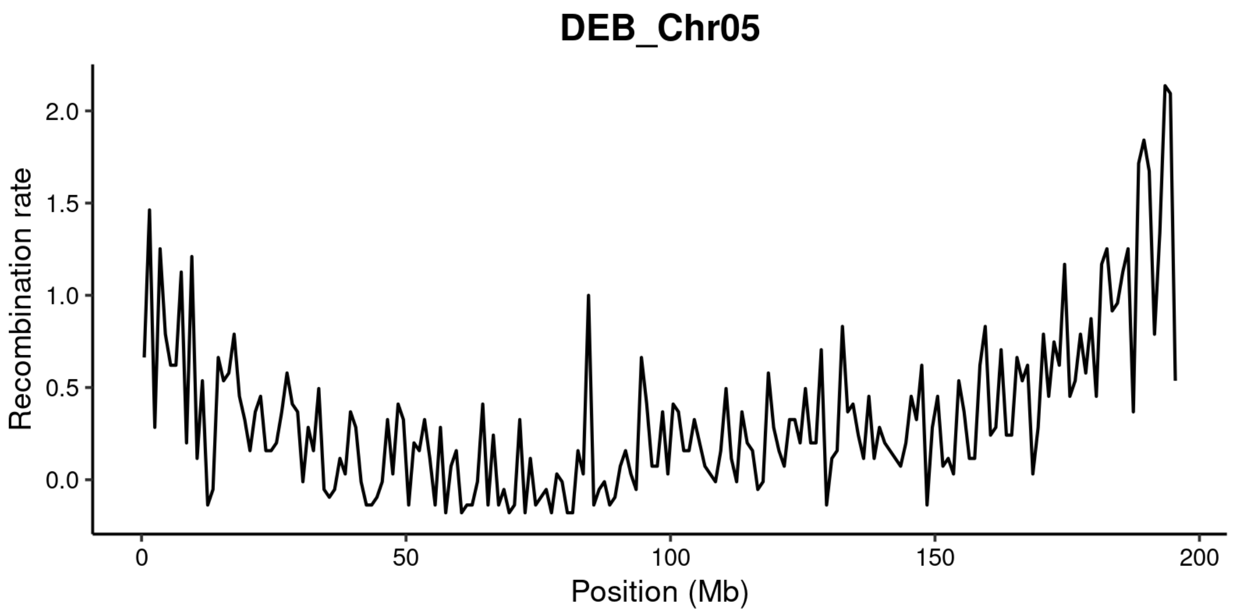
DEB_Chr05
Recombination rate
2.0
1.5
1.0
0.5
0.0
0
50
100
150
200
Position (Mb)

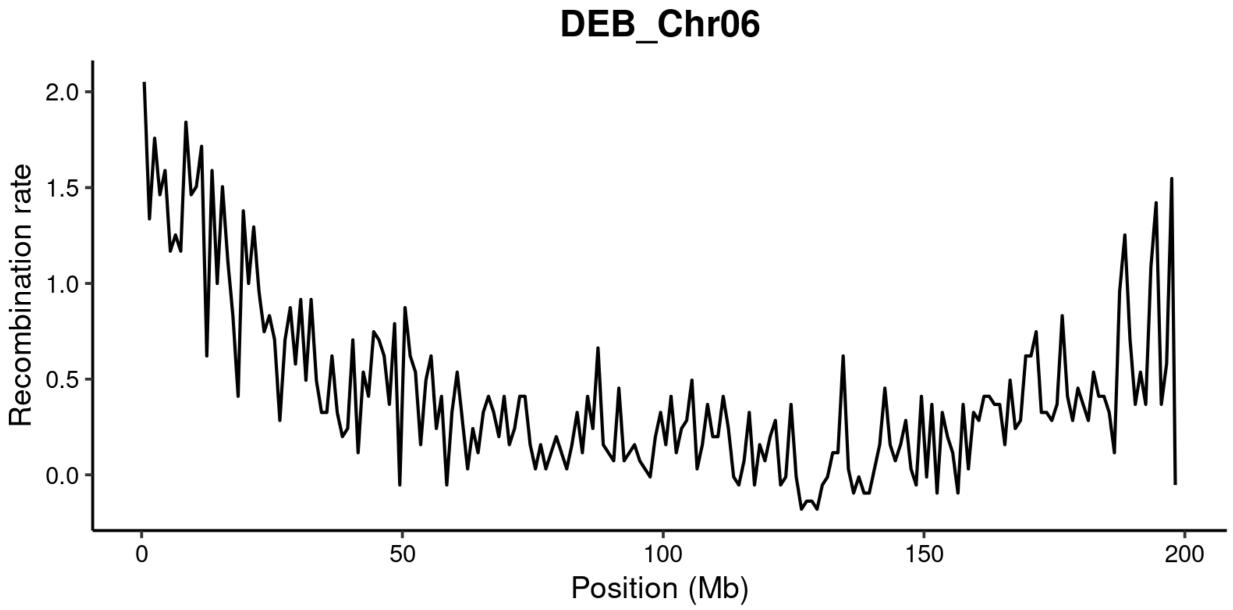
DEB_Chr06
Recombination rate
2.0
1.5
1.0
0.5
0.0
0
50
100
150
200
Position (Mb)

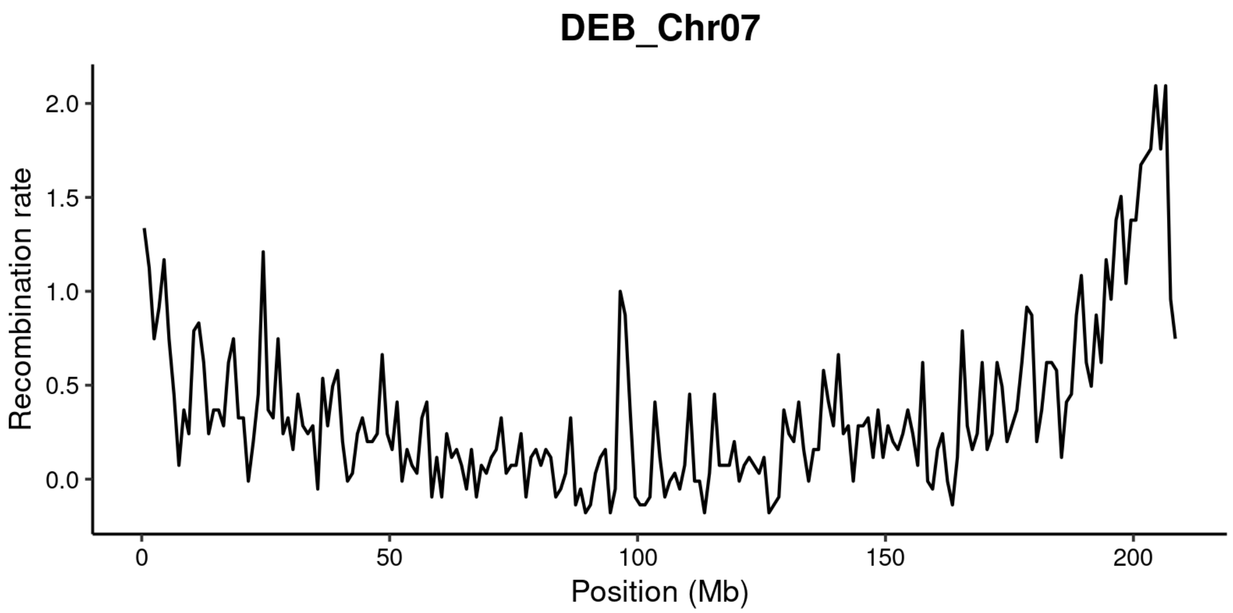
DEB_Chr07
Recombination rate
2.0
1.5
1.0
0.5
0.0
0
50
100
150
200
Position (Mb)

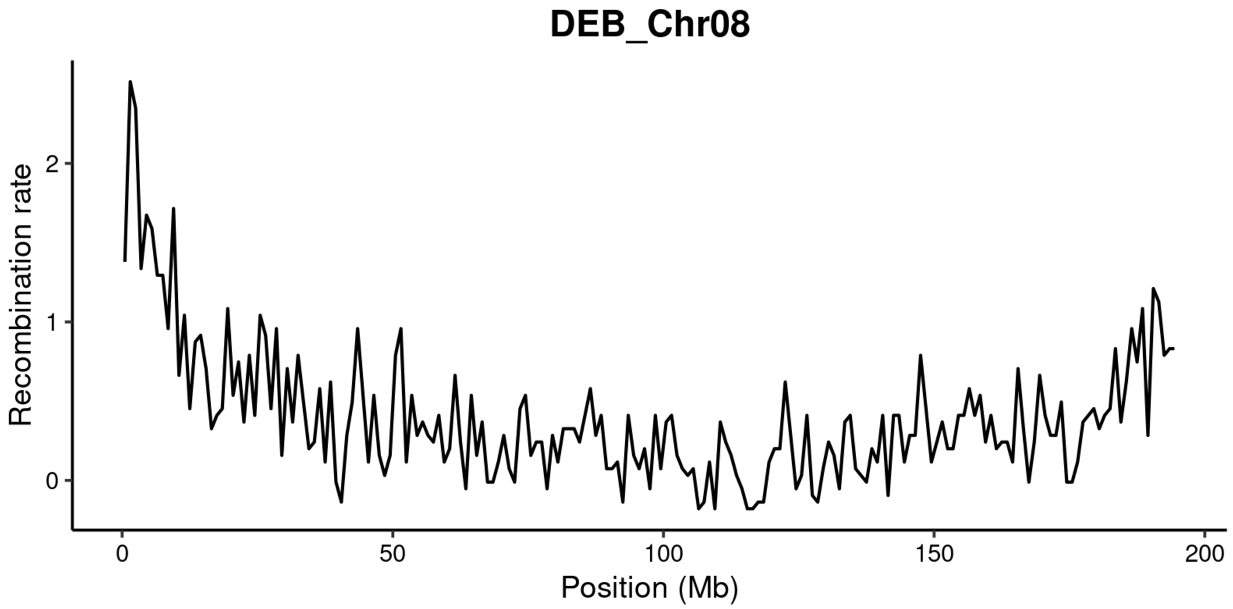
DEB_Chr08
Recombination rate
2
1
0
0
50
100
150
200
Position (Mb)

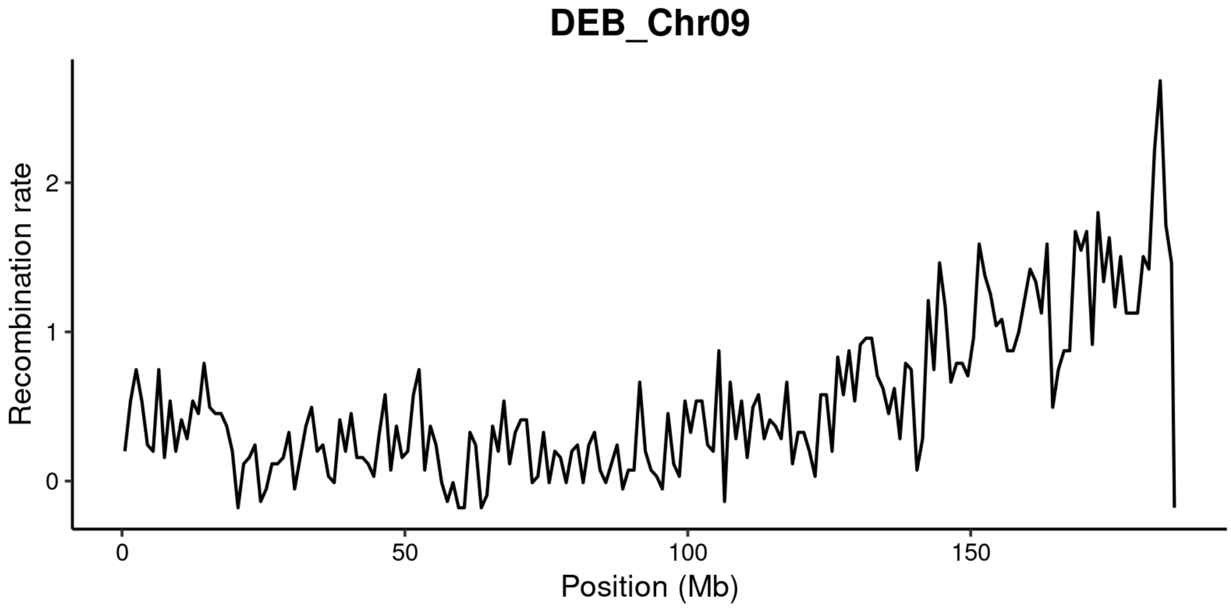
DEB_Chr09
Recombination rate
2
1
0
0
50
100
150
Position (Mb)

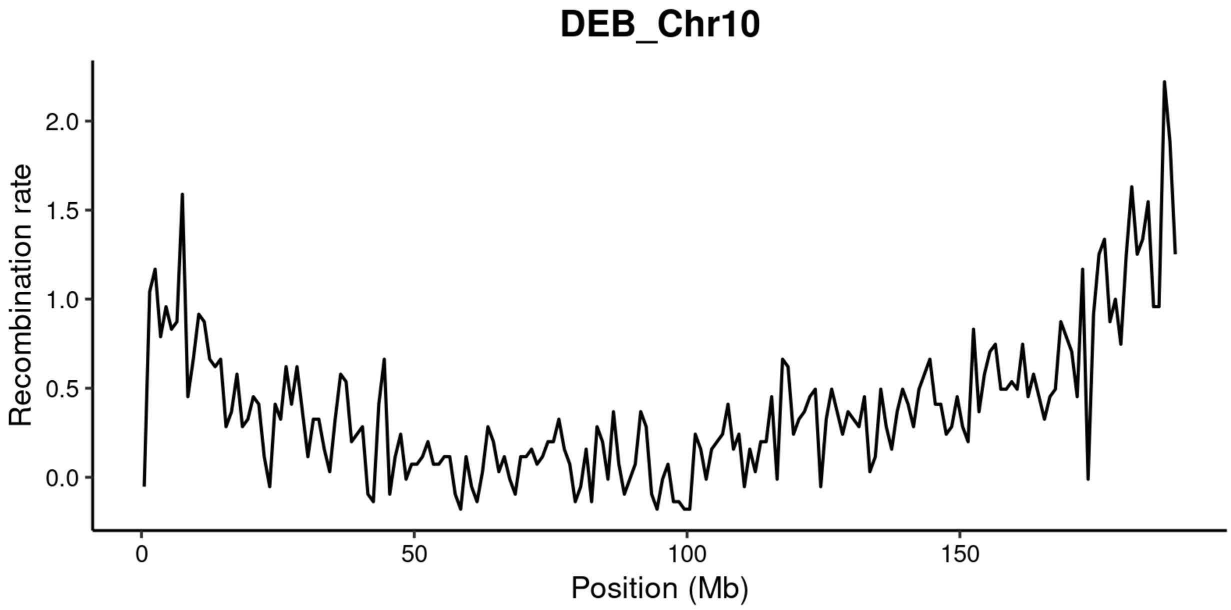
DEB_Chr10
Recombination rate
2.0
1.5
1.0
0.5
0.0
0
50
100
150
Position (Mb)

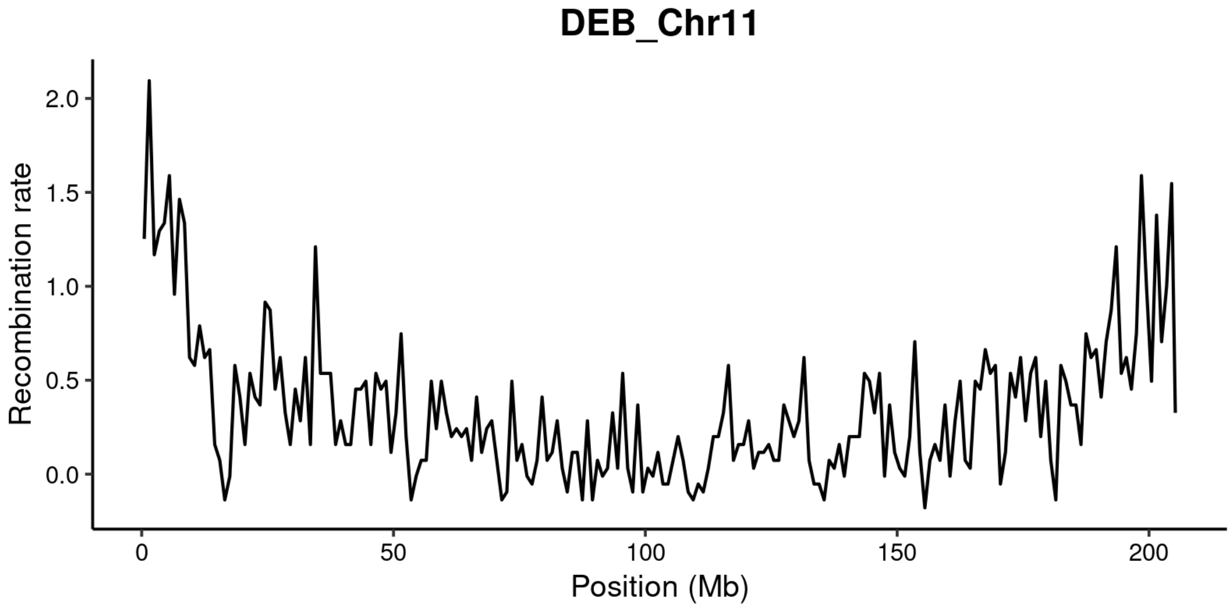
DEB_Chr11
Recombination rate
2.0
1.5
1.0
0.5
0.0
0
50
100
150
200
Position (Mb)

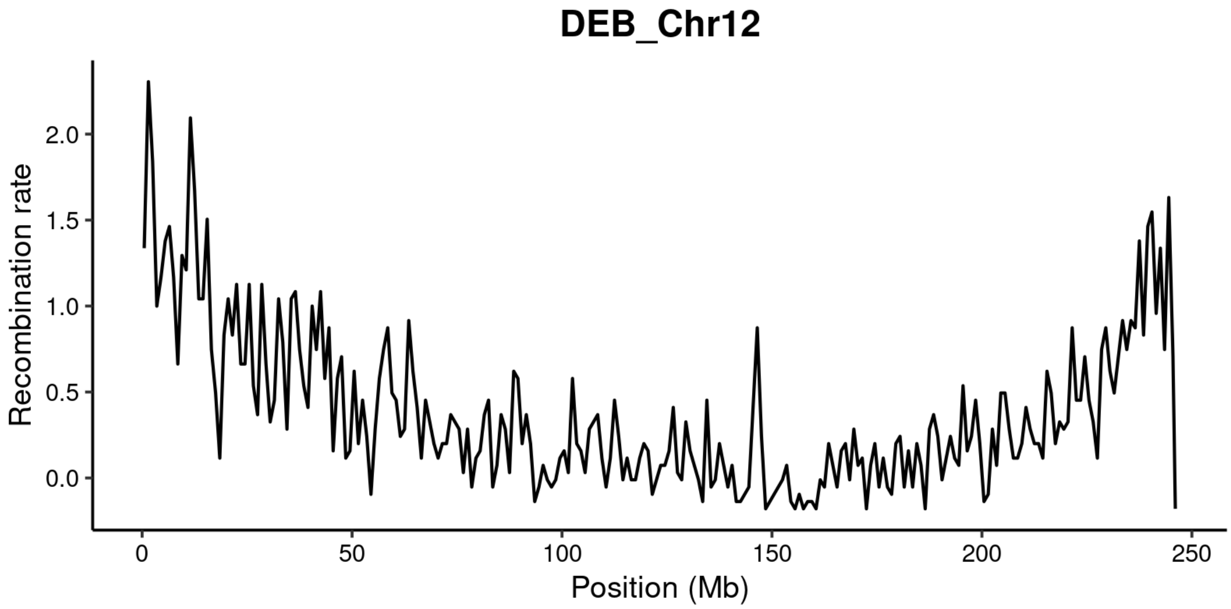
DEB_Chr12
Recombination rate
2.0
1.5
1.0
0.5
0.0
0
50
100
150
200
250
Position (Mb)

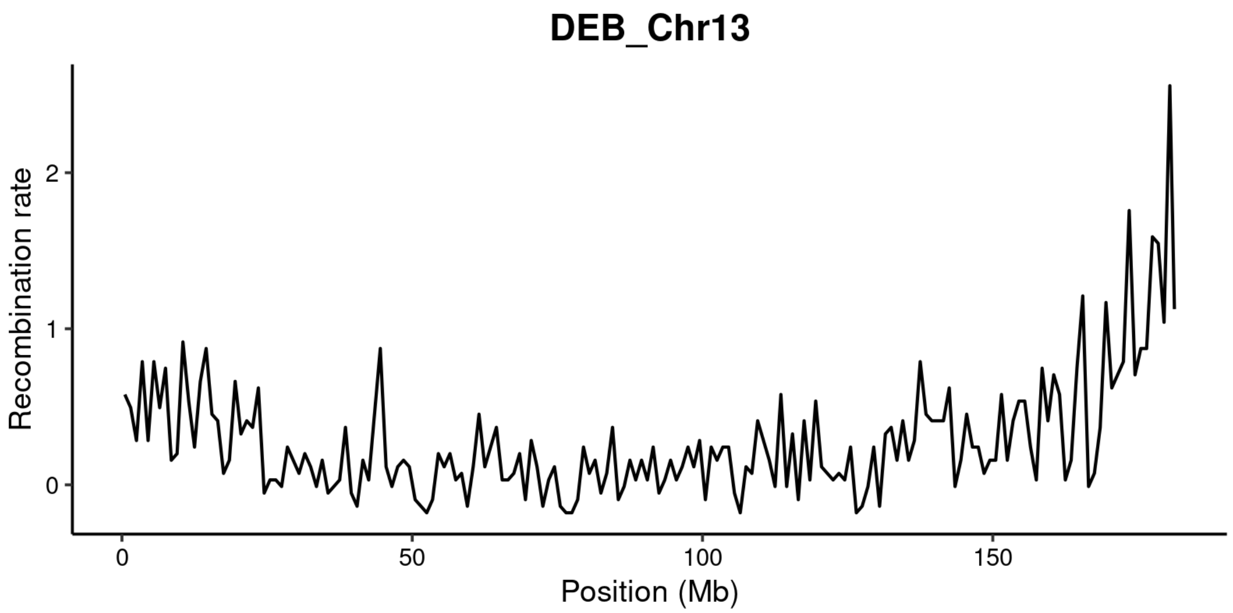
DEB_Chr13
Recombination rate
0
1
2
0
50
100
150
Position (Mb)

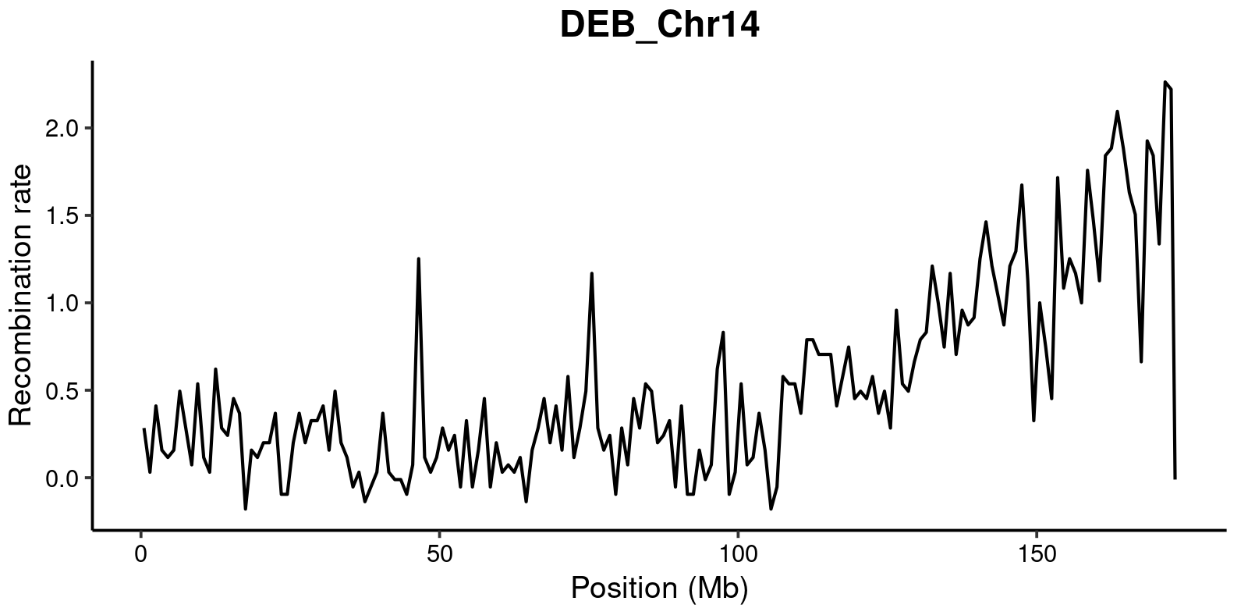
DEB_Chr14
Recombination rate
0.0
0.5
1.0
1.5
2.0
0
50
100
150
Position (Mb)

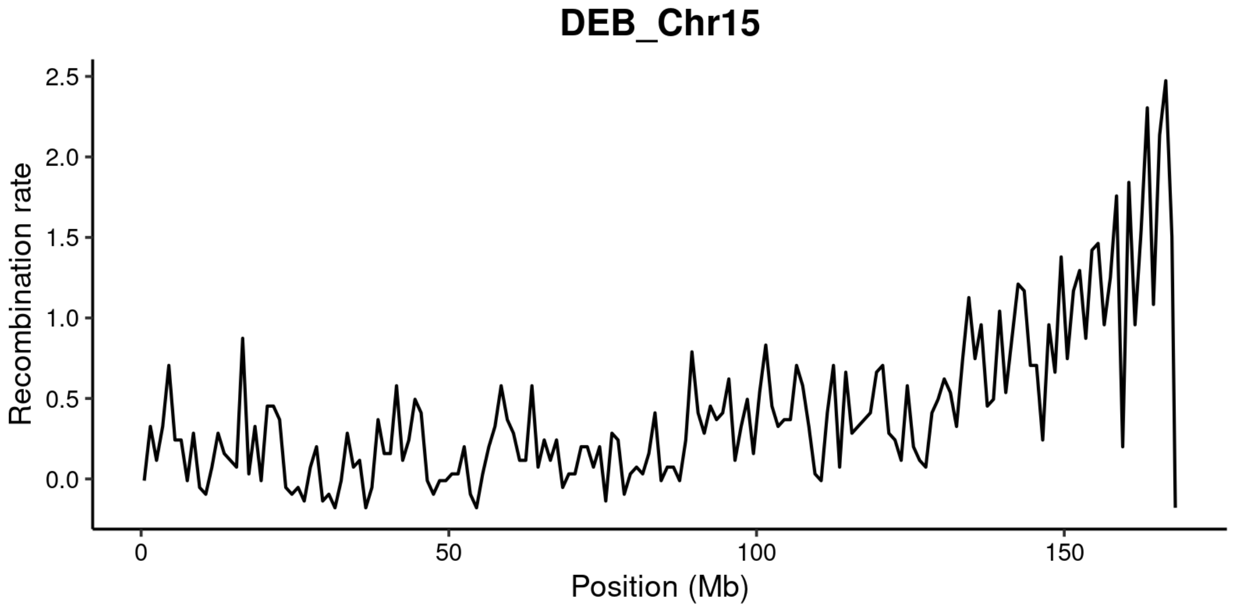
DEB_Chr15
Recombination rate
2.5
2.0
1.5
1.0
0.5
0.0
0
50
100
150
Position (Mb)

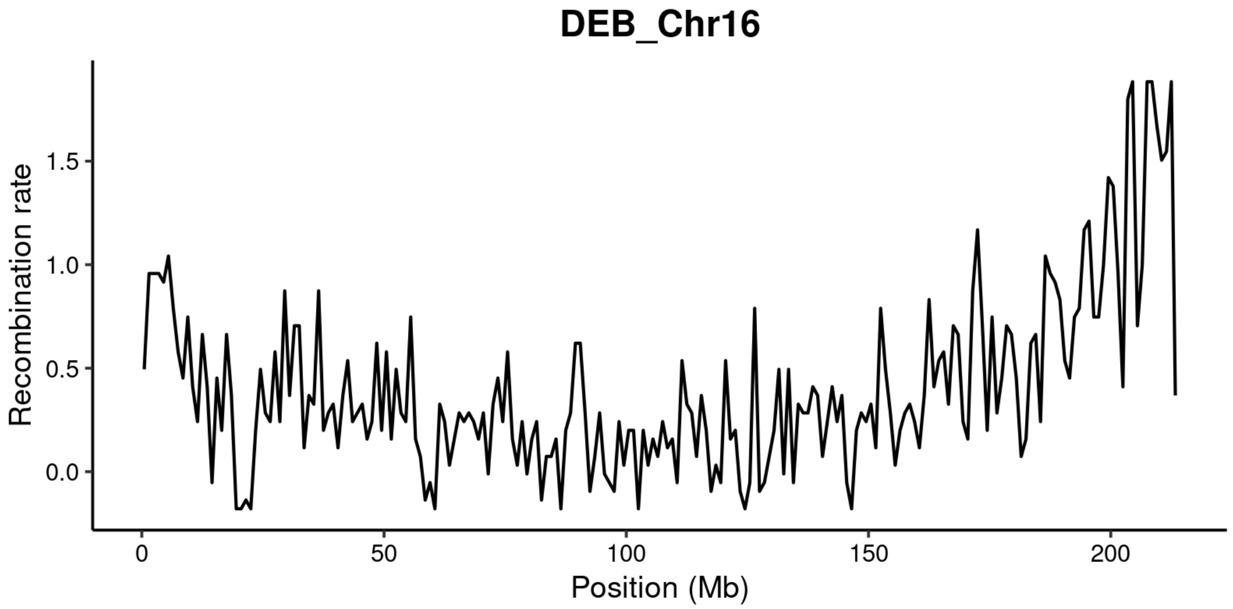
DEB_Chr16
Recombination rate
1.5
1.0
0.5
0.0
0
50
100
150
200
Position (Mb)

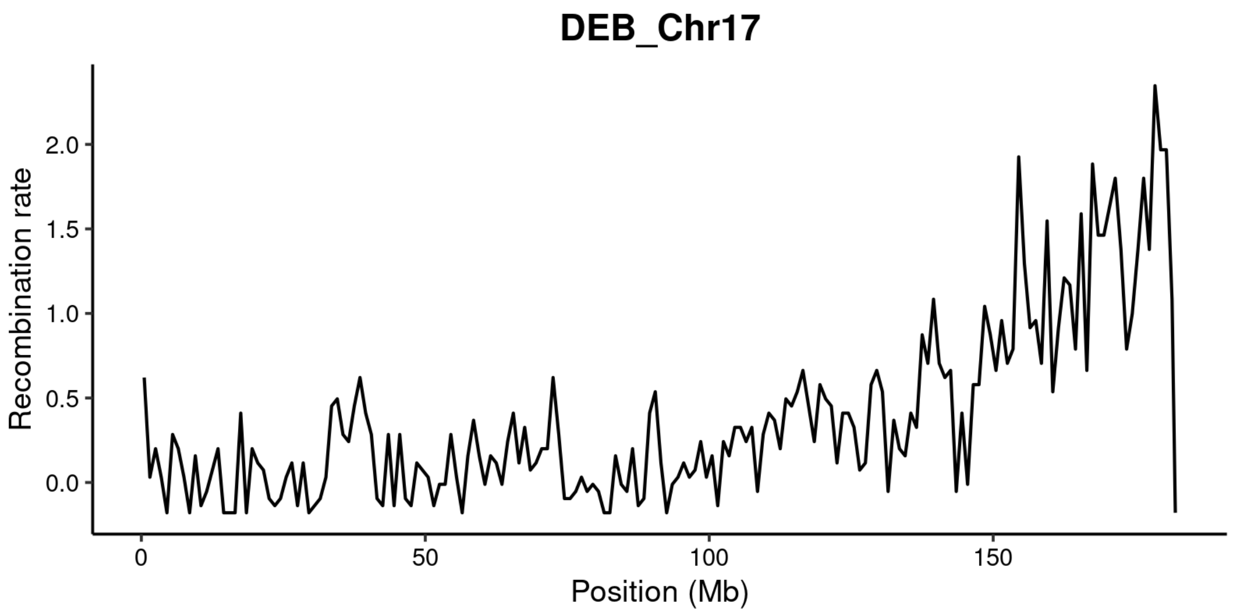

DEB_Chr17
Recombination rate
2.0
1.5
1.0
0.5
0.0
0
50
100
150
Position (Mb)

Figure 18 Recombination rate for HPraH2 inferred from gene density for all 17 chromosomes, predicted based on the recombination rate – gene density relationship from HA412.

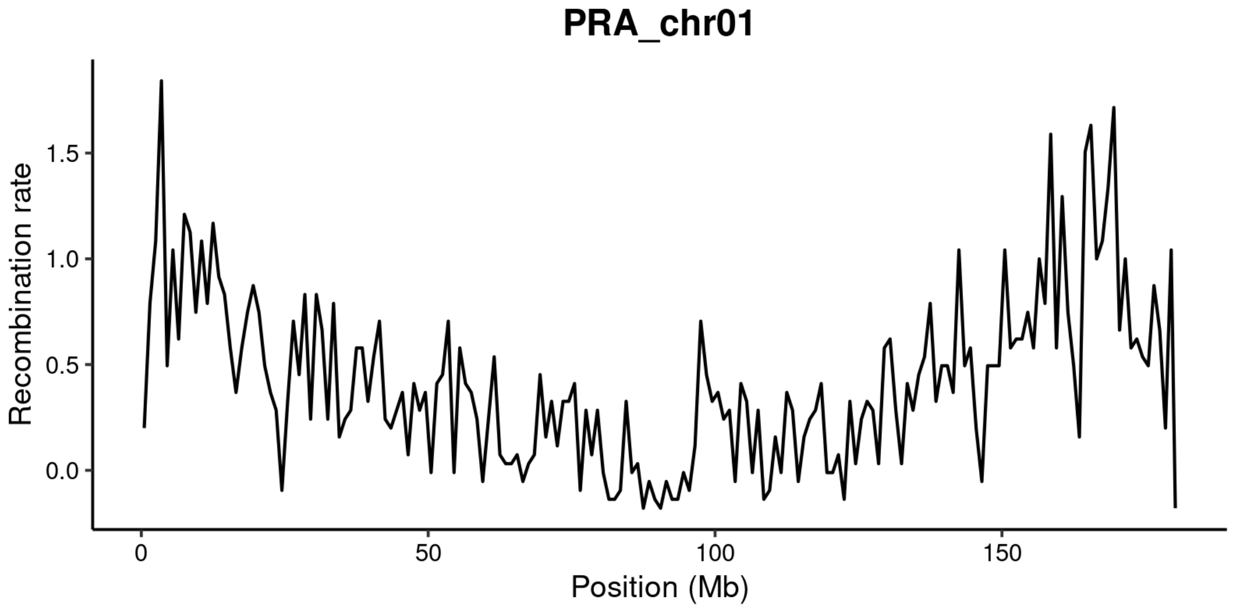


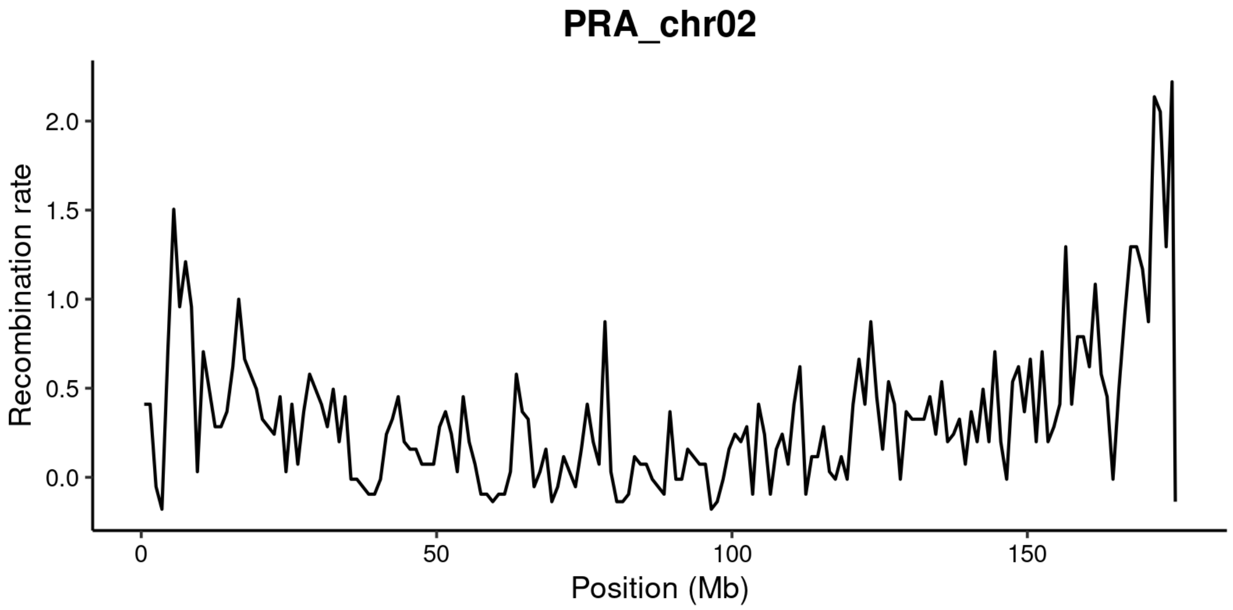

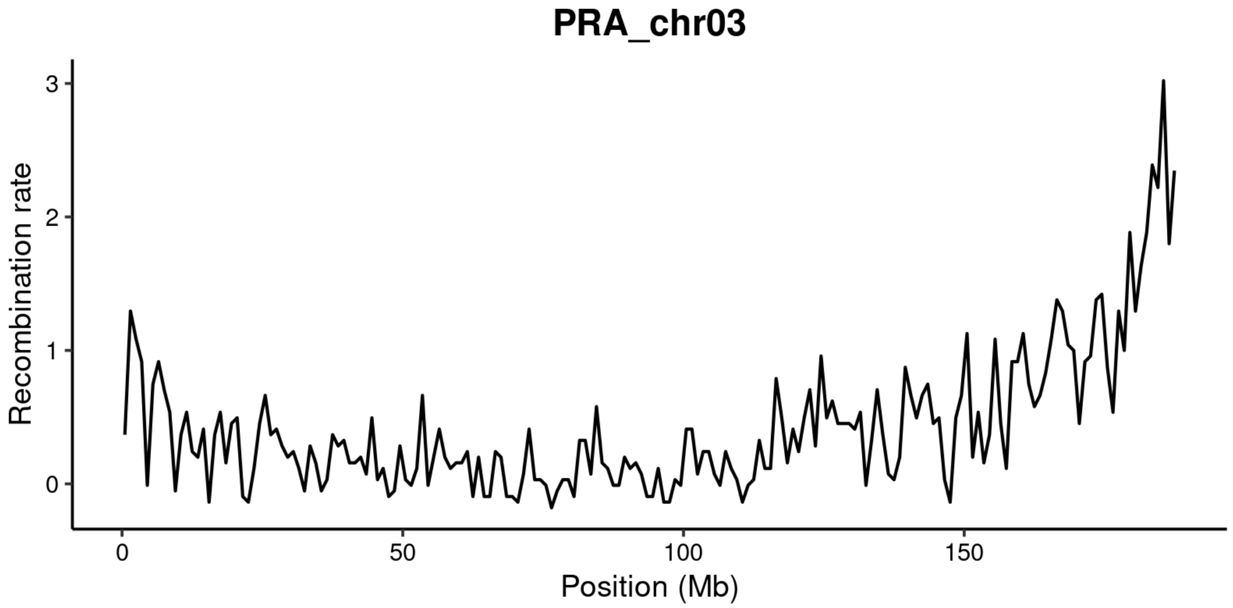
PRA_chr03
Recombination rate
0
1
2
3
0
50
100
150
Position (Mb)

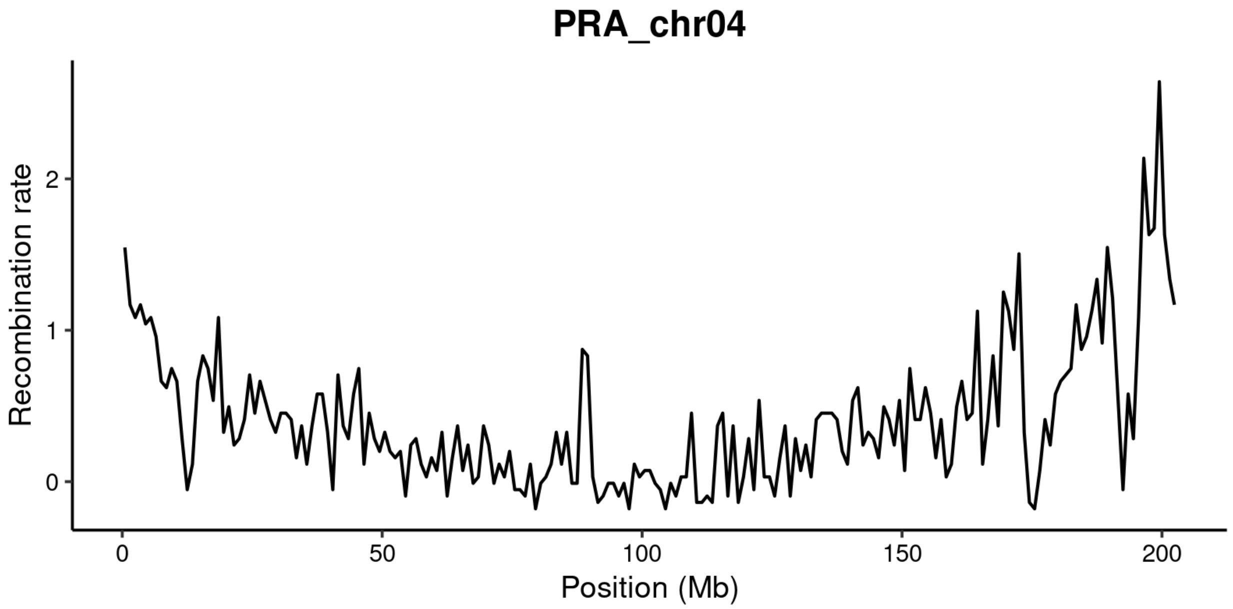
PRA_chr04
Recombination rate
0
1
2
0
50
100
150
200
Position (Mb)

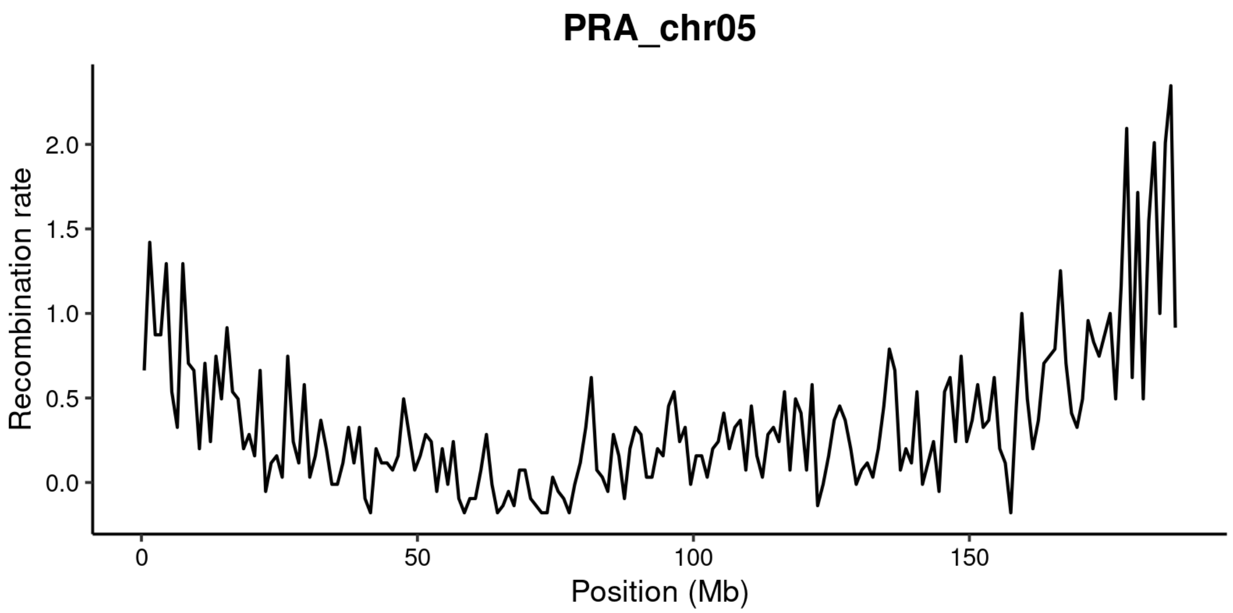

PRA_chr05
Recombination rate
0.0
0.5
1.0
1.5
2.0
0
50
100
150
Position (Mb)


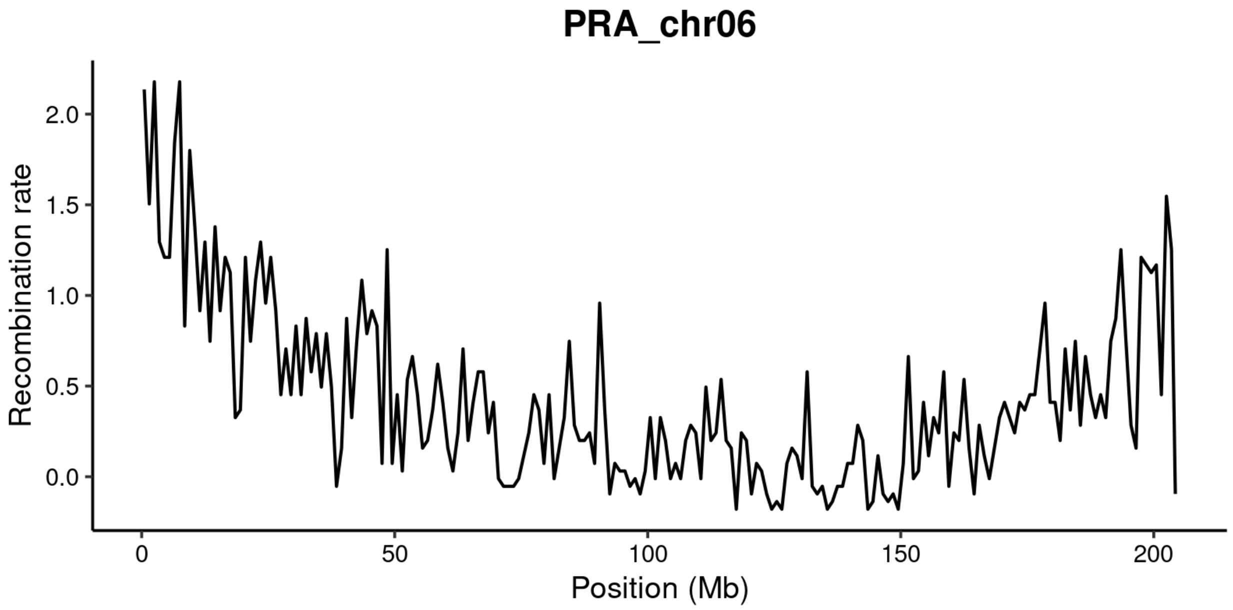

PRA_chr06
Recombination rate
0.0
0.5
1.0
1.5
2.0
0
50
100
150
200
Position (Mb)

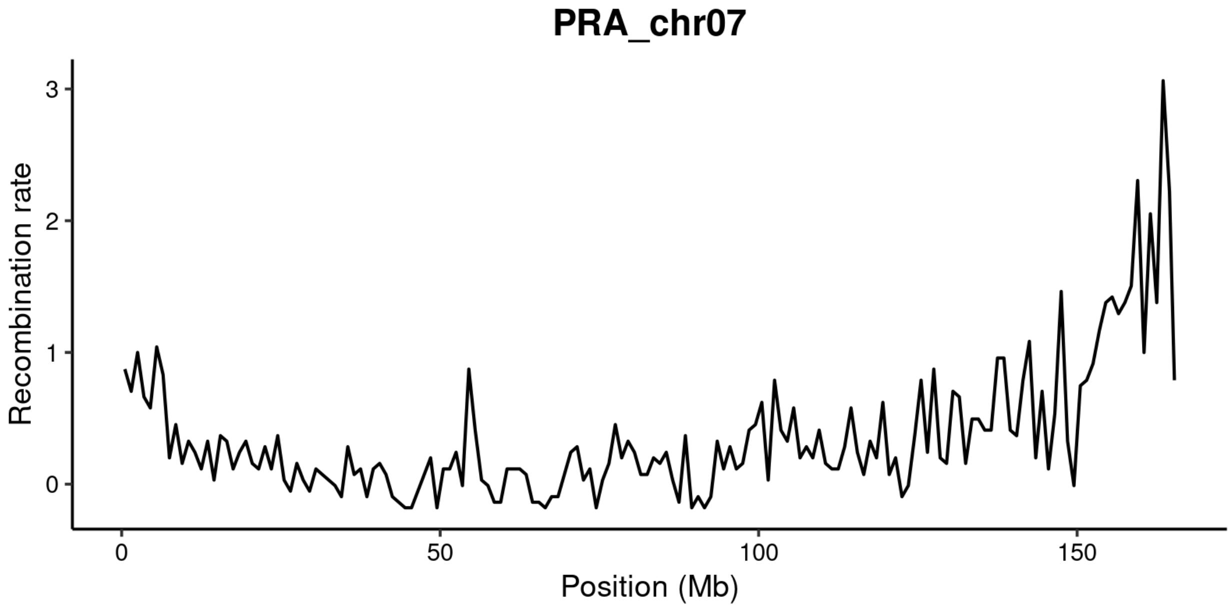
PRA_chr07
Recombination rate
0
1
2
3
0
50
100
150
Position (Mb)

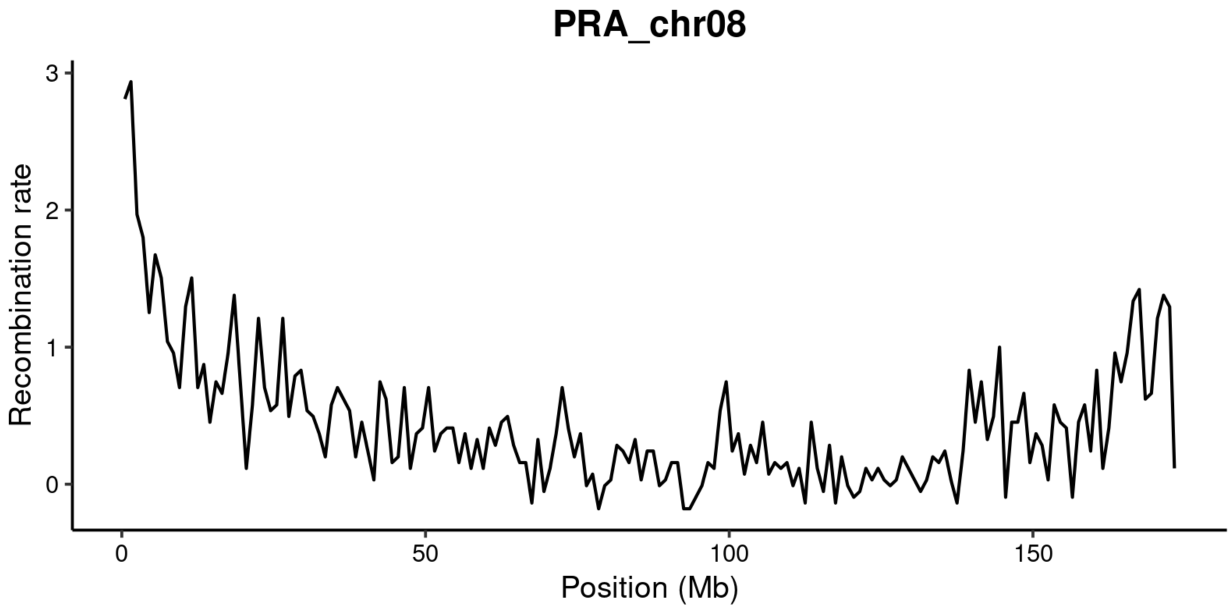
PRA_chr08
Recombination rate
0
1
2
3
0
50
100
150
Position (Mb)

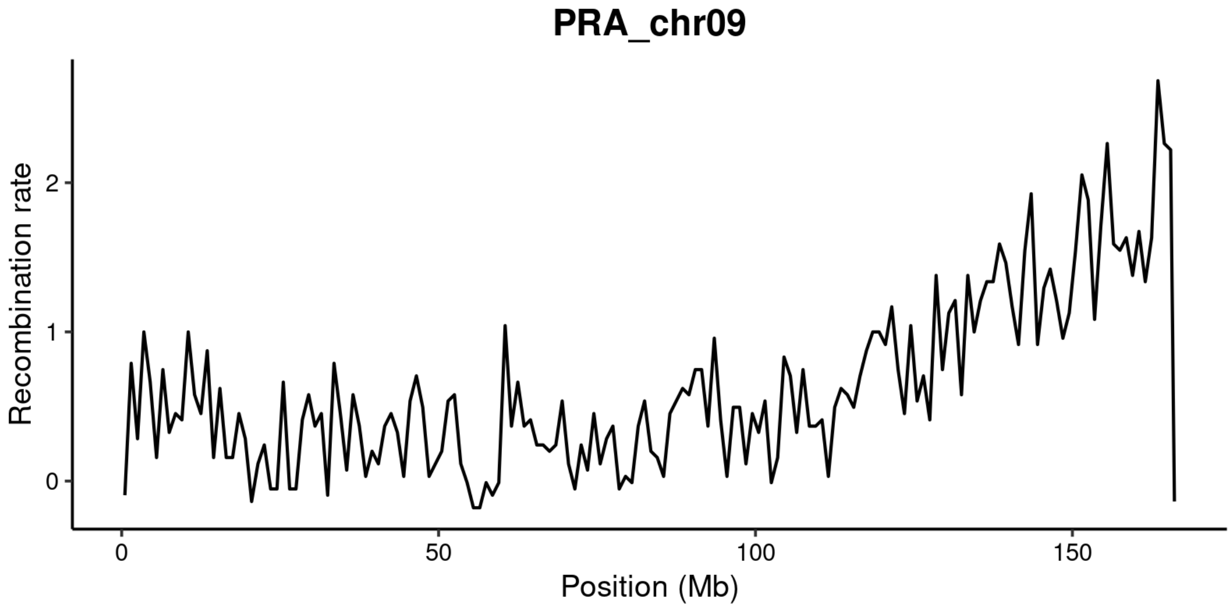
PRA_chr09
Recombination rate
0
1
2
0
50
100
150
Position (Mb)

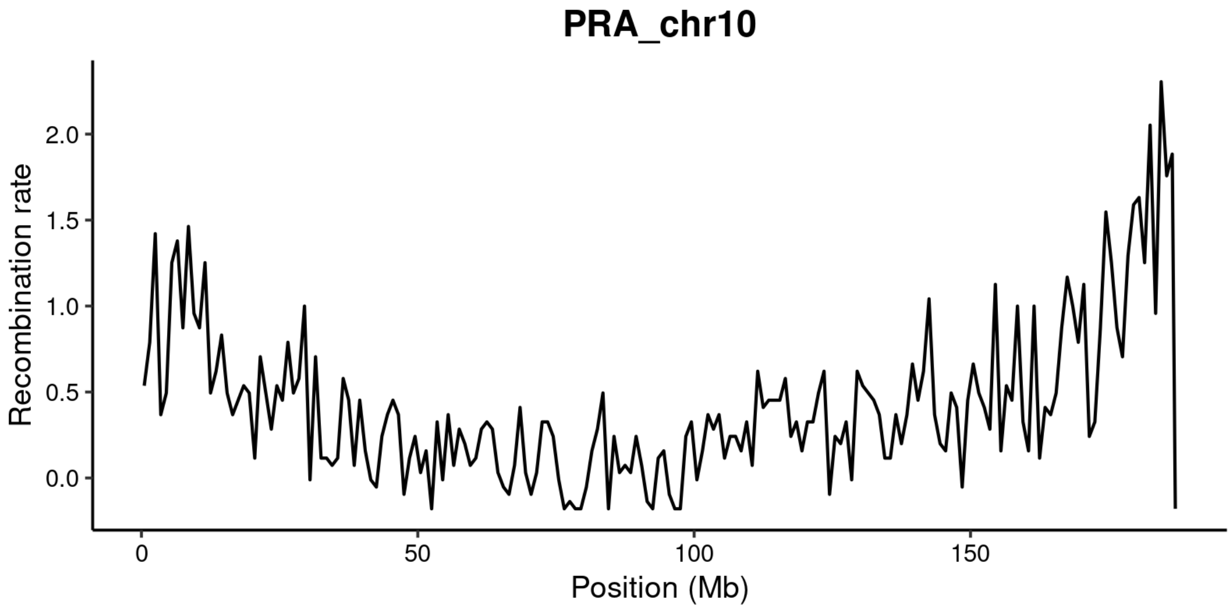
PRA_chr10
Recombination rate
0.0
0.5
1.0
1.5
2.0
0
50
100
150
Position (Mb)

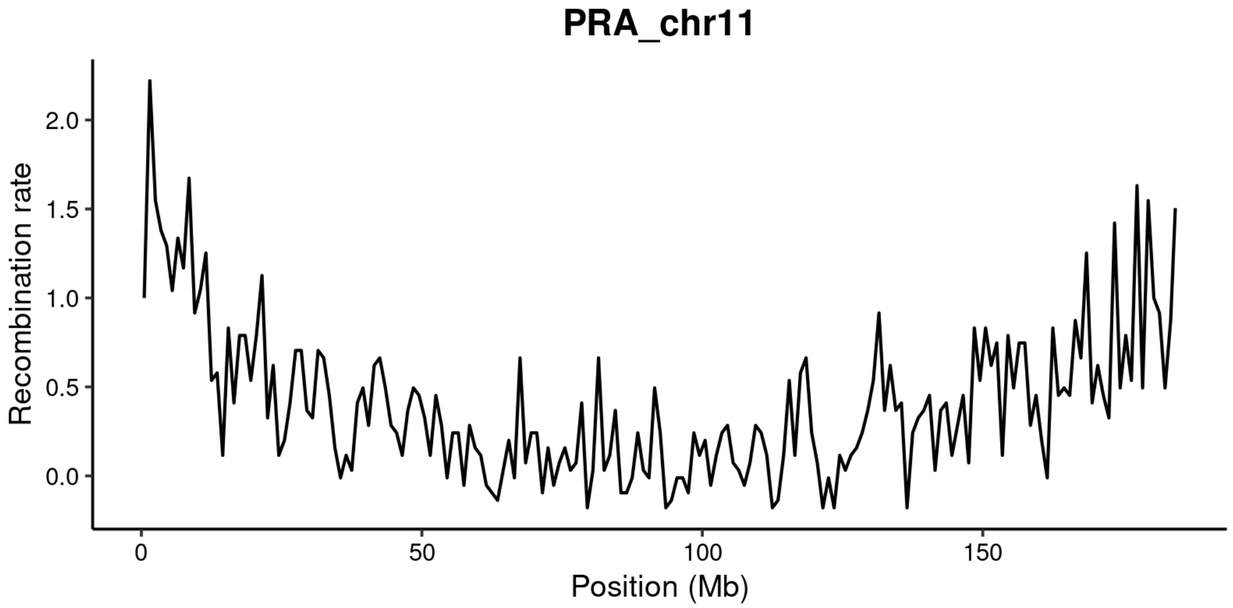
PRA_chr11
Recombination rate
2.0
1.5
1.0
0.5
0.0
0
50
100
150
Position (Mb)

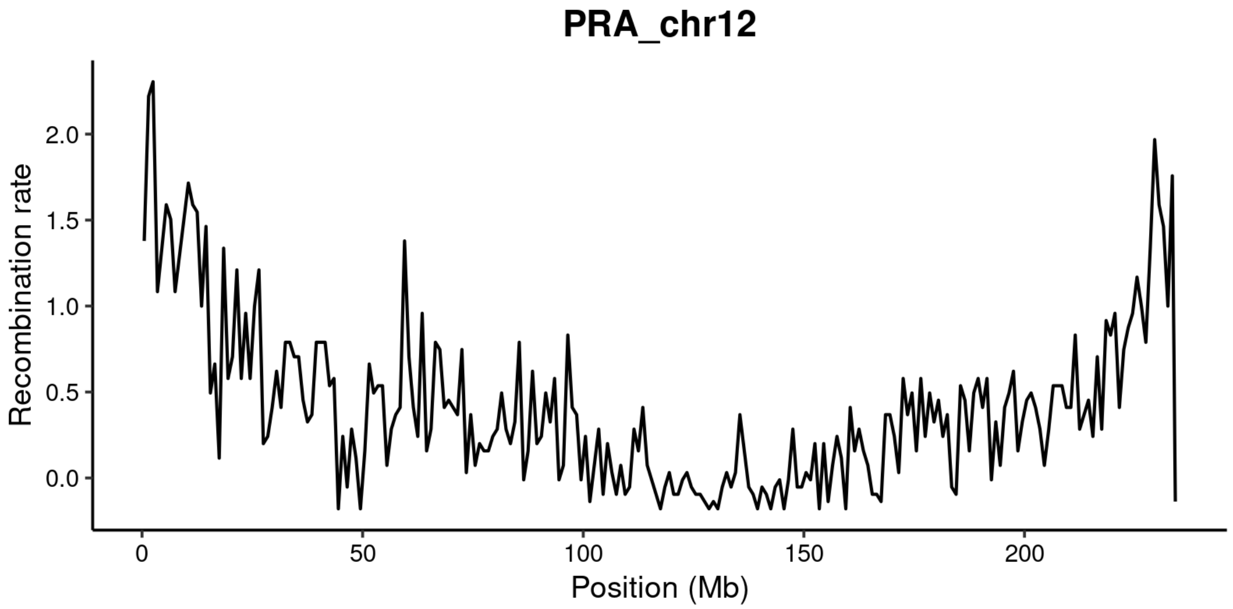
PRA_chr12
Recombination rate
2.0
1.5
1.0
0.5
0.0
0
50
100
150
200
Position (Mb)

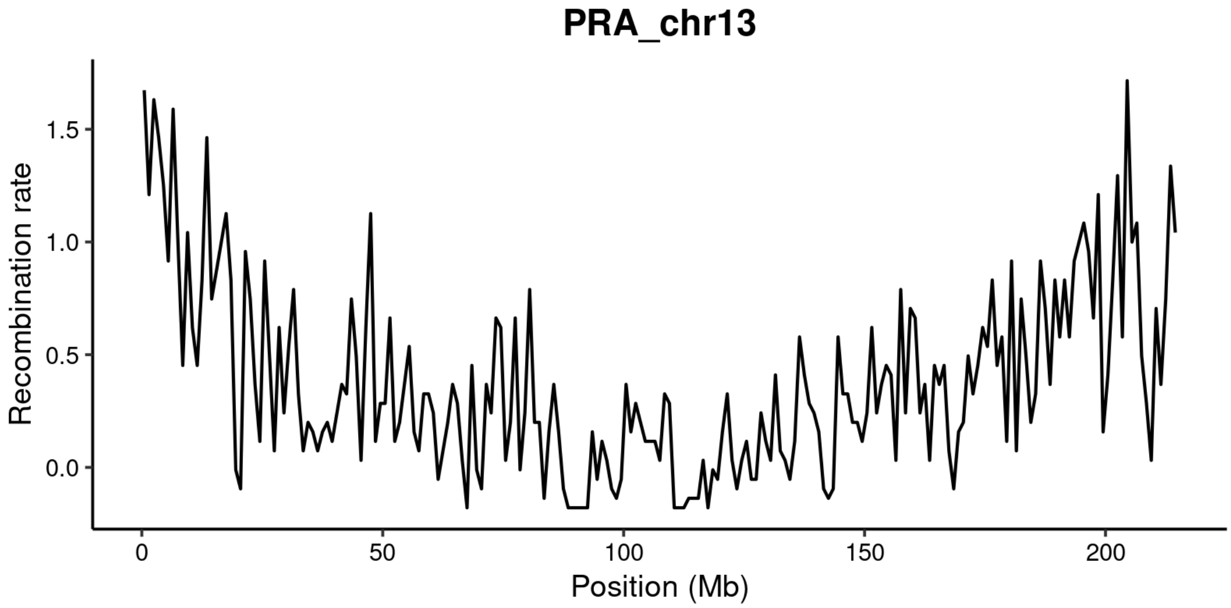
PRA_chr13
Recombination rate
1.5
1.0
0.5
0.0
0
50
100
150
200
Position (Mb)

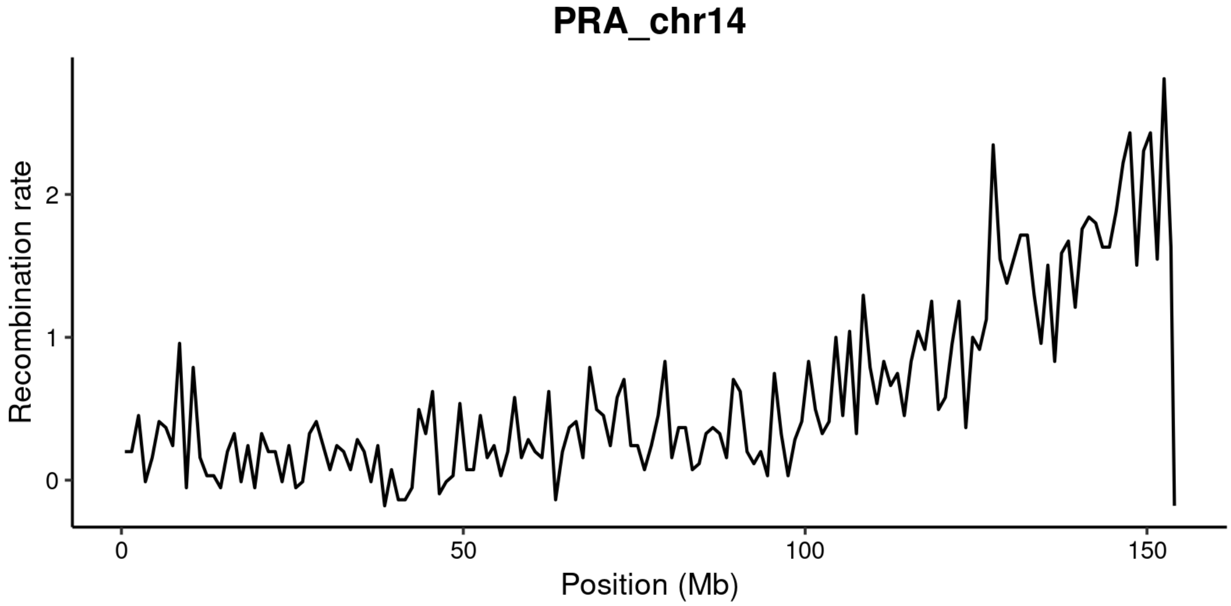
PRA_chr14
Recombination rate
2
1
0
0
50
100
150
Position (Mb)

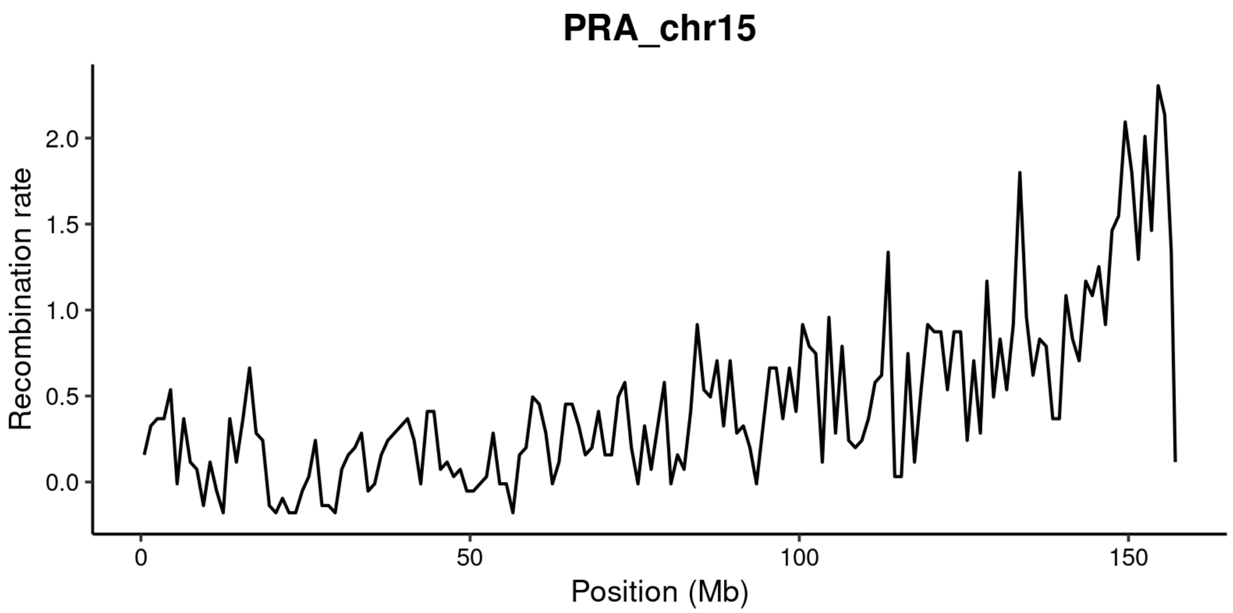
PRA_chr15
Recombination rate
2.0
1.5
1.0
0.5
0.0
0
50
100
150
Position (Mb)

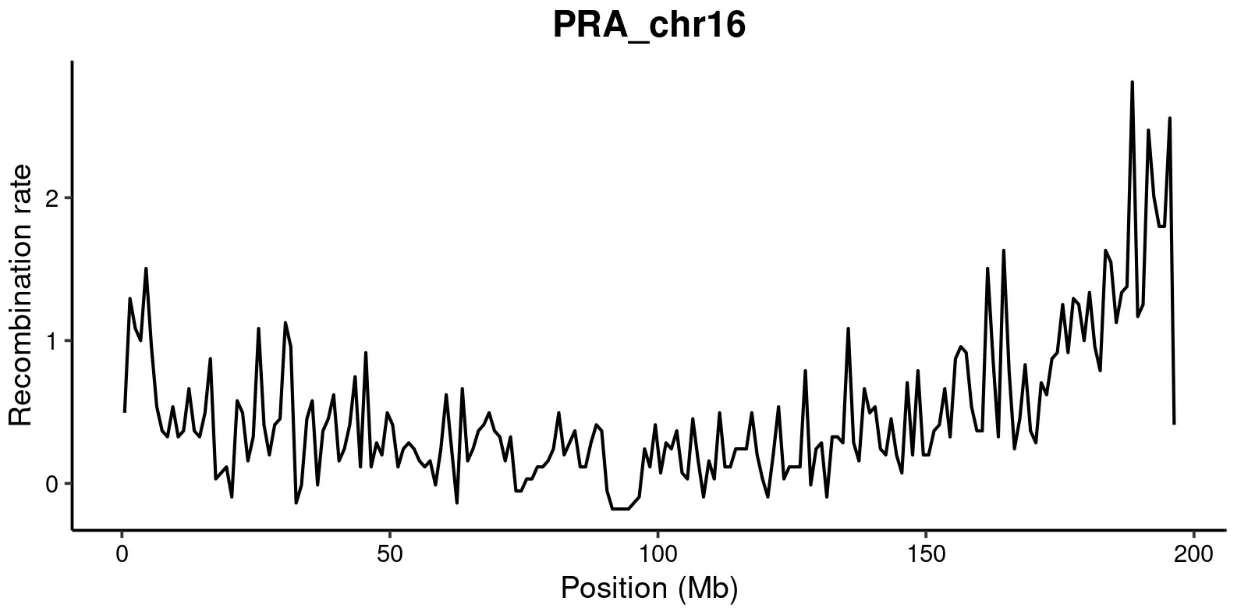
PRA_chr16
Recombination rate
2
1
0
0
50
100
150
200
Position (Mb)

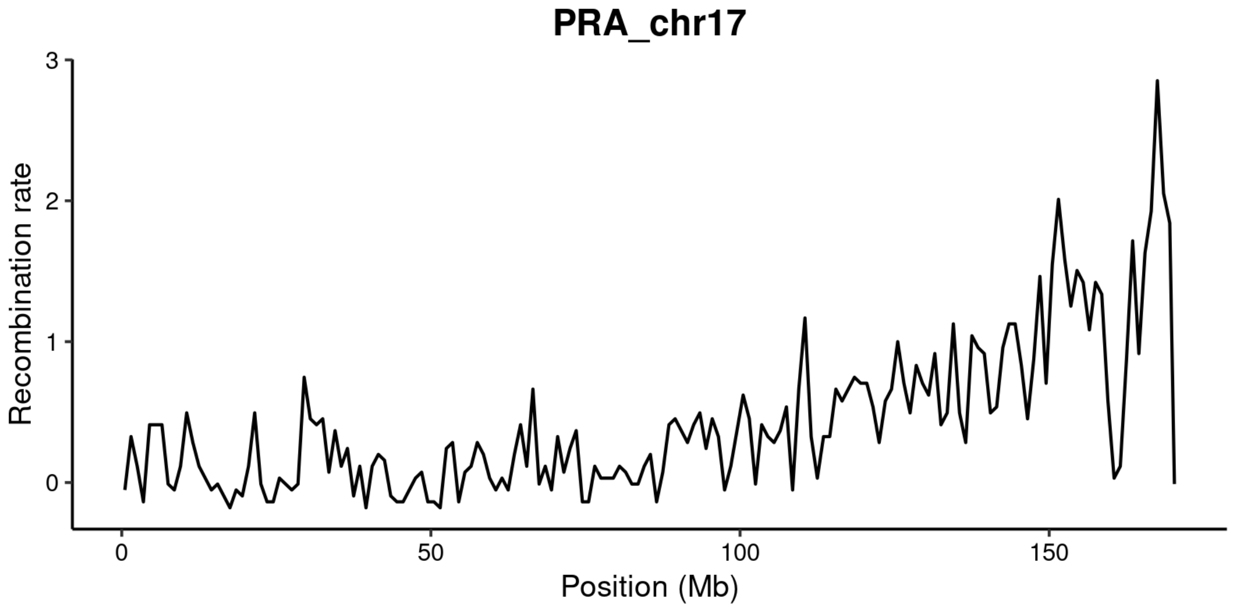
PRA_chr17
Recombination rate
3
2
1
0
0
50
100
150
Position (Mb)
