## Supplementary Document S1 for "Genomic and eco-geographic features of locally adapted inversions in wild sunflowers"

**Supplement Document 1:** Protocol details on extraction of High Molecular Weight (HMW) DNA extraction and quality check for the two reference genomes
(by CNRGV: INRAE-CNRGV, French Plant Genomic Resource Centre,
doi: 10.15454/1.5572367923221042E12 - http://cnrgv.toulouse.inrae.fr/)

Leaves from 3 weeks-old plantlets are harvested and frozen in liquid nitrogen, they are stored at -80°C. In the first step of the DNA extraction, leaves are ground into fine powder by manual crushing with pre-cooled mortar and pestle in liquid nitrogen. Then, a buffer containing SDS is used to lyse cells walls (NaCl 0.5M final, TrisHCl pH8.0-100mM final, EDTA pH8.0-50mM final, SDS 1.25% final, PVP40 1% final, sodium metabisulfite 1% final, β-mercapto-ethanol2% final). To increase the yield of DNA, this buffer must be heating before used, at 65°C-30min. Samples are incubated at 55°C for 20min. To inhibit the activity of RNA during the DNA extraction, add RNase A 0.45mg/ml final [Qiagen-1007885] and incubate on a rotator at 10rpm for 10min. After the lysis step, the precipitation of proteins and polysaccharides is performed by adding 5M potassium acetate (1/3 of lysis buffer volume), and 1 volume total of phenol:chloroform:IAA 25:24:1, and incubate at 20rpm for 10min. Contaminants are eliminated by centrifugation. Transfer upper phase and add 1 volume of chloroform:IAA 24:1 and incubate at 20rpm for 10min. After centrifugation, transfer upper layer to perform the extraction, be sure not to carry over interphase and lower layer. The third step consist to purify DNA using carboxylated magnetic beads (SpeedBeads magnetic carboxylate modified particles, GE Healthcare). Add 1 volume of beads solution previously prepared at 2%. Incubate at 10rpm for 1h at room temperature. The last step corresponding to two wash steps, with 70% ethanol freshly prepared, using the magnetic rack to trap beads and DNA. Finally, elution was carried out in buffered solution Tris-EDTA (tris 10mM, EDTA 0.1mM) pH8.0. It is recommended to let the tube in a magnetic rack for hours or even overnight. It is important to transfer the DNA in a new tube with a cut tip, to conserve the high quality of DNA.

With a spectrophotometric method, we can evaluate the purity of DNA samples. The ratio A260/280 > 1.8 indicates that DNA no containing protein. The ratio A260/230 > 2.0 indicates that DNA no containing any contaminants such as EDTA. UV spectrophotometry does not have the sensitivity to accurately measure low concentration of DNA. Using a fluorescent intercaling dye based method, such as Qubit Fluorometer (Thermofischer Scientific), we can do accurate measurements because of the binding dye to the selected molecule. The size of HMW DNA is evaluated using Femto pulse system of Agilent society.
